# Best Practices for Modeling Arthropod Lifetimes in a Bayesian Framework

**DOI:** 10.64898/2025.12.16.694707

**Authors:** Piper O. Zimmerman, Leah R. Johnson

## Abstract

Most arthropods are ectothermic, with multiple performance traits being constrained by environmental temperature. Through the effects on traits, temperature therefore constrains when, where, and how large are arthropod populations. Because many arthropods are of relevance to humans, for example as pests or disease vectors, biologists have spent substantial time trying to understand the mechanistic relationship between temperature and traits. For example, in mosquito and other vectors species, the lifespan of vectors - the number of days the adult survives – is important in determining both the size of the vector population and whether or not vectors will likely be able to transmit pathogens. Often thermal traits such as lifetime are modeled using Thermal Performance Curves (TPC) – functions that describe the relationship between temperature and traits mathematically. Many functional forms with many possible shapes have been proposed as TPCs. However, the effects of the distribution of data around these shapes and of common data transformations on how well we can infer the TCPs has been relatively ignored. In this paper we use simulated data on vector lifespans, inspired by mosquito data, to explore the reliability of inference under different assumptions about the data on our ability to accurately infer a known TPC. Using a Bayesian approach, we are also able to quantify the effects of data assumptions and transformations on uncertainty in estimates. Our results suggest that mismatches between a true and assumed distribution of data around a TPC can greatly increase uncertainty. Further, some transformations of the data before analysis are more likely to lead to biased results than others. Based on our results, we make suggestions for best practices in the analysis of arthropod thermal trait data such as lifespan and related traits.

## 1. Introduction

Temperature is a fundamental driver of ectotherm biology, influencing how organisms reproduce (Ciota et al., 2014), grow and develop (Verant et al., 2012; Ciota et al., 2014), and survive (Milosavl-jevíc et al., 2020). Because ectothermic organisms depend primarily on external heat sources to regulate their body temperature (Andrade et al., 2015), many physiological traits, or measures of physiological performance that can have an impact on fitness (McGill et al., 2006; Nespolo et al., 2014), are largely dependent on temperature. These traits have been studied in a range of contexts, from host–pathogen relationships (Blanford and Thomas, 1999) to fungal growth and disease susceptibility (Verant et al., 2012; Voyles et al., 2017). Temperature also shapes broader ecological processes such as herbivory rate (Lemoine et al., 2014), metabolic efficiency (Little et al., 2020), and flight behavior (Kenna et al., 2021), highlighting its influence in determining how organisms function.

The relationship between traits and temperature is especially critical in the context of climate change. As global temperatures rise, ectothermic organisms, including many disease-vectors, are experiencing shifts in their population dynamics (Iwamura et al., 2020). Because the species that transmit pathogens responsible for diseases such as dengue, West Nile virus, malaria, and more are ectothermic (WHO, 2014), minor thermal changes can lead to global health crises (Sinka et al., 2020). Each year more than one billion people are infected and more than one million people die from vector-borne diseases, while many more are left with permanent complications (WHO, 2014). Warming temperatures are expected to expand the ranges suitable for transmission of vector-borne diseases. For example, as many as 1.33 billion new people may be shifted into areas that are suitable for Zika virus transmission by 2050 (Ryan et al., 2021) and similar trends are also predicted for other *Aedes*-borne diseases (Ryan et al., 2019; Iwamura et al., 2020). To better predict these patterns and the areas that will be affected by them, it is necessary not just to understand temperature-trait relationships but also to quantify them.

A popular way to quantify the response of an ectotherm trait across a range of temperatures is with a thermal performance curve (TPC). TPCs are inherently nonlinear (Huey and Stevenson, 1979; Huey and Kingsolver, 1989; Bulte and Blouin-Demers, 2006; Cohen et al., 2020), as well as generally unimodal and asymmetric (Angilletta, 2009; Clarke, 2017; Kontopoulos et al., 2024). For unimodal TPCs there is an “optimal” temperature where the trait is maximized (or minimized, depending on the trait). TPCs are often bounded at critical thermal limits (Angilletta, 2006) that may be also be determined by measures of tolerance or mortality (Kenna et al., 2021). There are many different functions used to model these relationships, such as polynomials (Palaima and Spitze, 2004; Kellermann et al., 2019), Gaussians (Verant et al., 2012; Angilletta, 2006; Huey and Kingsolver, 1993), or Brieres (Briere et al., 1999).

TPCs are often fit using parameter estimation methods such as non-linear least squares (NLLS) (Shi and Ge, 2010; Low-Décarie et al., 2017; Rebaudo et al., 2018; Padfield et al., 2021) due to the inherent nonlinearity of these relationships at the more extreme temperatures (Briere et al., 1999). Recently, using Bayesian methods has become more popular (Johnson et al., 2015; Ryan et al., 2021; Childress and Letcher, 2017; Landry Yuan et al., 2021; von Schmalensee et al., 2021). Using a Bayesian framework allows for incorporation of prior knowledge of parameters, while also allowing quantification of uncertainty around these parameters (Voyles et al., 2017). This combination is particularly helpful with sparse data (Johnson et al., 2015; Angilletta, 2006) since prior knowledge about similar species can be used to constrain estimation.

The focus of researchers is generally using these thermal performance curves to quantify the relationship between temperature and some physiological trait. There has also been a focus on finding the “best TPC”, or a set of guidelines to find the best model (Kontopoulos et al., 2024; Rezende and Bozinovic, 2019). However, before the functional form of the TPC is chosen, i.e. before this “best fit” is considered, it is important to consider the underlying “true” distribution of the data itself, sometimes referred to as the data-generating mechanism. Parameter estimates, as well as which TPC equation is selected, can be impacted by assumptions about the data-generating mechanism and how the data are handled. These aspects are often overlooked in the discussion of TPC fitting. Little work has been done to explore the original data-generating mechanisms for physiological traits, nor to explore the consequences if this is assumed incorrectly.

For example, the lifetime of an arthropod, defined as the number of days an individual survives as an adult, is one of the many physiological traits affected by temperature (Loeb and Northrop, 1917; Pearl, 1928; Milosavljevíc et al., 2020). Lifetime is an important trait because it can have impacts on other traits such as total fecundity (Djawdan et al., 1996; Cator et al., 2020), or on summary metrics such as population growth (Huxley et al., 2021). When it comes to lifetime data itself, the two typical distributional assumptions when modeling are that the lifetime is distributed according to an exponential distribution, or that it is distributed according to a Weibull distribution (Lawless, 2011; Kalbfleisch and Prentice, 2011). These contradictory assumptions about the data-generating mechanism are an example of considering the distribution of the data first. Whether these data follow an exponential or a Weibull distribution could potentially affect subsequent analyses.

Another consideration that can affect an analysis is the form of the data. Many studies rely on historical or previously published data, which often present only summary statistics such as means or medians rather than individual-level data (Ciota et al., 2014). Because individual-level data are often unavailable, many temperature-dependent trait analyses utilizing past data must rely on reported means (Deutsch et al., 2008). Additionally, some researchers transform these data by inverting the lifetimes to model mortality rate, or the inverse of lifetime, directly, as mortality rate is often more relevant for calculating population dynamic measures such as maximum population growth rate (Huxley et al., 2022).

With different types of data and different assumed distributions, this creates many ways to represent ectotherm lifetime data. Rather than attempting to find the golden standard or the best TPC equation while modeling physiological traits, the aim of this study is to explore the impact of a contradicting true and assumed data-generating process, as well as the structure of the data used, on the quality of known TPCs fit to ectotherm lifetime data. We use simulated lifetime data to assess the consequences of a researcher making the incorrect assumption about the distribution that the data came from. Quantifying the impacts of this will help to ensure a unified understanding among researchers regarding the data-generating mechanism in these analyses.

We detail the simulation experiment that we used to investigate the consequences of assuming a mismatched data-generating mechanism when fitting a thermal performance curve. We introduce the conceptual framework of how the exponential and Weibull distributions differ and the various forms of data that researchers may model and how these two considerations can impact one’s analysis. Then, we describe our simulation including the TPC equation, the basis of how data were generated for each simulation experiment, and specific settings used in these experiments. Afterwards, we present several visuals and quantification of each method’s ability to recover key features of the curve, as well as summaries of parameter coverage to compare our methods. Finally, we discuss these results and make suggestions for best practices under different availability regimes and possible repercussions of an incorrect distributional assumption in a TPC analysis.

## 2. Conceptual Framework

Often lifespan or lifetime is summarized across a population by a single number, such as a life expectancy or average lifetime. However, different species or populations can exhibit disparate patterns. For example, organisms such as frogs and fish produce thousands of offspring, most of whom die immediately before mortality levels out (Rauschert, 2010), leaving a very small proportion of those who survive to adulthood (Herreid and Kinney, 1966; Brockelman, 1968; Calef, 1973). In contrast, some species, such as humans and sheep, have the majority of their individuals surviving to adulthood and dying at older ages rather than just after birth (Rauschert, 2010). One can imagine that if a lifetime summary from the first case was used to make a prediction about numbers of individuals surviving over time assuming a pattern more like the second case, then the predictions would be less accurate than using the relevant assumptions and vise versa. Thus, being able to quantify the distributional assumptions for lifetimes and incorporate these into models should provide better predictions.

These different patterns of lifetime can be represented mathematically by probability distributions. Two relatively common distributions for this purpose across domains (e.g., epidemiology, demography, sociology, or medical research (Liu, 2012)) are the exponential and Weibull distributions. Each of these corresponds to different assumptions about how likely deaths are across ages, and thus can lead to very different predicted distributions of lifetimes, even when a summary statistic (like expected lifetime) is the same.

The exponential distribution is “memory-less”. That is, it assumes equal risk of death at each time increment. For example, the probability of an arthropod dying in the next day remains the same after 15 days as it does after just 1 day. This results in a pattern where most individuals die quickly, but a few survive for a long period of time. This can be represented mathematically by the probability density function (PDF) of the exponential distribution:

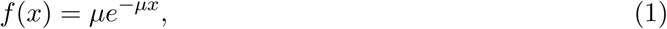

where *µ* is the rate and *x* is the random variable of interest, which in our case is the observed lifetime.

One advantage of the exponential distribution is its relative simplicity. It has only one parameter, which can be estimated in a straightforward way. Suppose that we observe *n* identical, independently distributed samples from an exponential distribution with rate parameter *µ* (i.e., 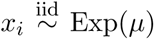 for *i* = 1*, …, n*), we can estimate the rate parameter with the maximum likelihood estimator, 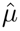, given by

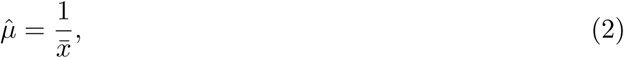

where *x̄* is the mean of the sample. The ease of this estimation makes it appealing for modeling; many models that use estimated rates, such as ordinary differential-equation (ODE) frameworks, assume exponential waiting times (Anderson and May, 1991; Diekmann et al., 2013).

In contrast, the Weibull distribution allows the risk of death to depend on age, so the chance of survival is not constant at each time increment. Specifically, it assumes that the probability of death starts low and increases over time. This difference in assumption in the risk of death affects the frequency of observing very short lifetimes. Thus, it typically predicts lifetimes that are more concentrated around intermediate values (although it is flexible enough to capture other patterns). The PDF of the Weibull distribution is:

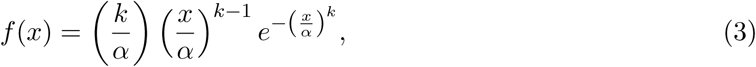

where *k* is the shape, *α* is the scale, and *x* is the random variable of interest. Unless the value of *k* is known, the Weibull distribution does not have a simple, closed form maximum likelihood estimator like the exponential distribution (Cohen, 1965), making it slightly more complicated to estimate the parameters.

We illustrate the differences between the exponential and Weibull distributions using a simple example where the exponential and Weibull mean are the same (Figure 1). Although the mean is the same for both distributions, the actual range and pattern of values around the mean can vary greatly depending on the choice of the parameters. For the values chosen, the pdf of the exponential distribution (Figure 1a) and thus the samples from the distribution (Figure 1c) include a very high proportion of very short lifetimes compared to the Weibull. Interestingly, the exponential also has more long lifetimes than the Weibull in this case. These kinds of differences will affect the estimation and prediction of the models. Thus, even if our analysis manages to capture the true value of this mean, we would make different predictions about the proportion of long or short lifetimes under the two models due to the difference in shape between the two distributions. For example, if an exponential distribution is assumed during fitting but the data are from a Weibull distribution, we may be able to accurately estimate the mean but not the other moments, as the exponential distribution is not flexible enough to capture the observed patterns. The resulting fit will make poor predictions about the range and potential values of lifetimes. Since the amount of time an adult vector survives can impact the probability that it will transmit pathogens (Cator et al., 2020), poorly estimating the distribution of adult lifetimes will result in poor estimates of transmission risk.

**Figure 1:**
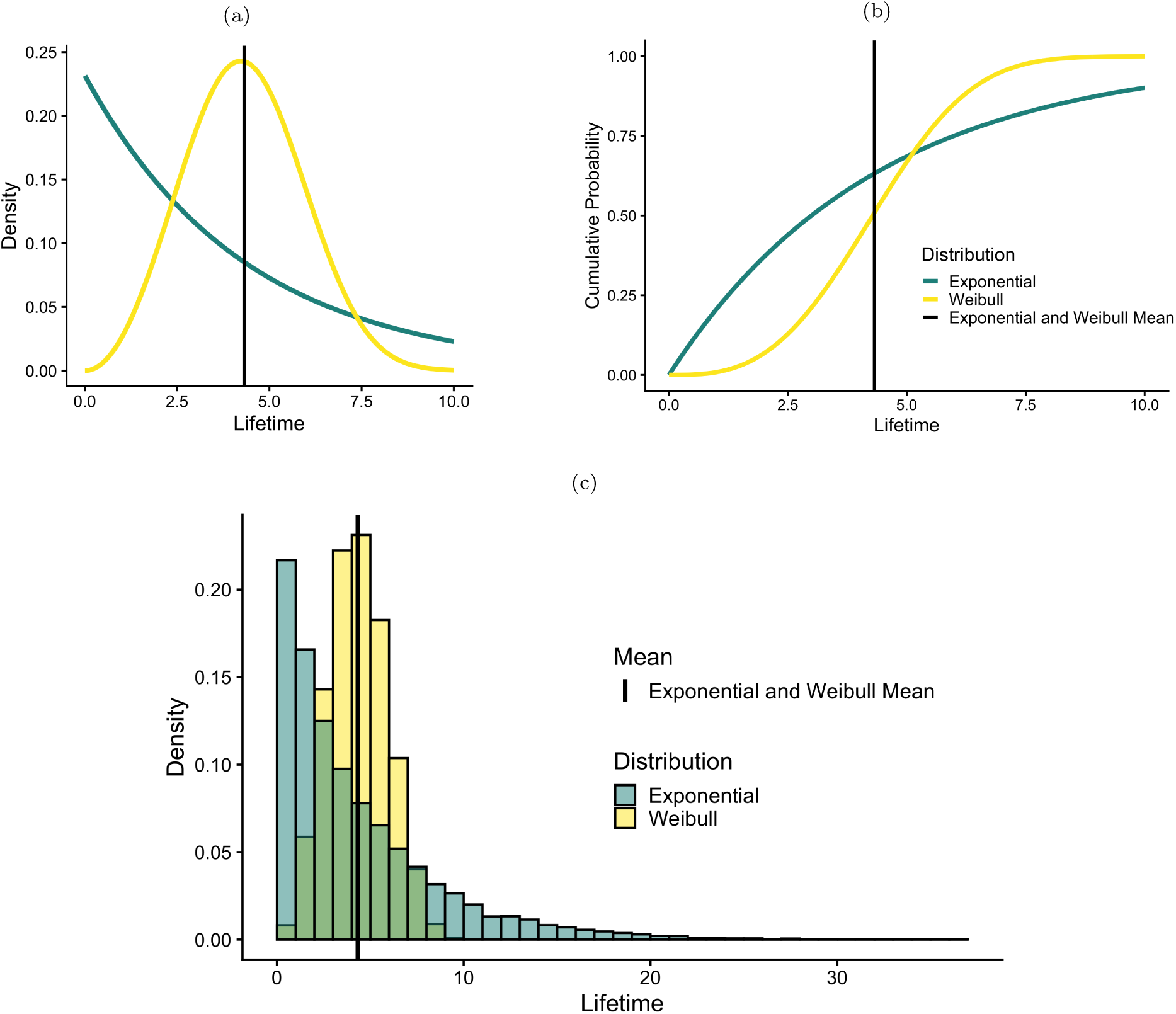
Comparison of exponential and Weibull distributions with identical means. 1a PDFs of the exponential (teal line) and Weibull (yellow line) distributions. 1b CDFs of the exponential (teal line) and Weibull (yellow line) distributions. 1c Histograms of sample lifetimes drawn from the exponential (teal) and Weibull (yellow) distributions. For both distributions, we assume a mean of 4.32 (solid vertical line in all subfigures). More specifically, the rate of the exponential distribution is 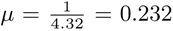, while the scale of the Weibull distribution is *α* = 4.84, and the shape is *k* = 3.

This effect on estimation and predictions has a knock-on effect when we fit a thermal performance curve (TPC). When fitting a TPC, we assume that the data at each temperature will have a distribution where the mean (or median) will shift with temperature according to the functional form of the TPC. For both the exponential and the Weibull, the variance/pattern shifts with the mean (e.g., for the exponential the variance is 1*/µ*^2^). Thus, the difference in patterns between the two distributions will not even be the same across temperatures, resulting in predictions that diverge more at some temperatures than others.

The effects on estimation and prediction assume that we are able to estimate parameters directly from individual lifetime data. However, individual lifetime data are not always available. Instead, data are frequently transformed, aggregated, or summarized. If a researcher is modeling something other than the raw individual lifetimes, the distributions of these summaries will not be the same as the raw data and will further depend on both the underlying distribution and the chosen summary. Thus, inference is expected to be poor or misleading if the same model assumptions are used for summarized and unsummarized data.

We have seen four common data transformations: (1) taking the mean of the data (mean data); (2) taking the inverse of the individual data to obtain a “rate” (inverse data); (3) data that are averaged and then inverted (inverse of the mean (IoM) data); and (4) individual data inverted and then averaged (mean of the inverse (MoI) data). For most of these transformations, there are no closed-form solutions of their distributions (but see the Supplementary Material for a subset for which closed form distributions or approximations exist). Thus, it is unclear how these transformations may impact the inference of an underlying TPC, including the uncertainty in the TPC and the variability of the predictions. In the remainder of the paper we will explore how the combination of the underlying distribution of lifetimes (exponential or Weibull) together with the summarization of data impacts inference using a simulation experiment.

## 3. Simulation Experiments

In this section, we describe the details of the simulation experiments that we use to explore how the underlying data-generating mechanisms and the approaches to summarizing the data from Section 2 jointly impact the inference of the parameters of the TPC model. We first detail the specific TPC function that we use and how it relates to the parameters in both the exponential and the Weibull distributions. We then describe the simulated datasets used in the simulation experiments and review the summaries of the data. Next, we introduce the models we fit to our simulated data, including those where we fit to the raw, individual-level lifetime data, as well as those we fit to a transformation or summary of these data. As the models are fit using a Bayesian approach, we also detail the prior distributions on all model components and specific model settings. Finally, we outline the methods and metrics that we use to compare between the different models.

### 3.1. TPC Specification

When we generate data from an exponential distribution, at each temperature we assume that the mortality rate, *µ*, is given by

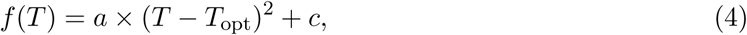

where *T* is temperature, *a* controls the width of the function, *T*_opt_ is the temperature at which the mortality rate is minimized, and *c* is the minimum mortality rate (the inverse of the maximum lifetime) at the optimal temperature. To ensure the curve is concave up, reflecting higher mortality rates as temperatures move away from optimum, we constrain *a* to be positive. In addition, we constrain parameter *c* to be positive to ensure a positive mortality rate. Note that since the rate parameter in the exponential distribution is the inverse of the scale parameter, *f* (*T*) in this case corresponds to the inverse of the lifetime. Thus, at each temperature, the data are exponentially distributed with rate parameter equal to *f* (*T*), with the mean of the distribution given by 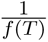. Modeling in terms of the mortality rate is common in a variety of mathematical frameworks, including dynamical systems. Thus, this simple quadratic mortality rate aligns our simulation framework with these common practices.

The Weibull distribution uses a scale parameter rather than a rate parameter, that is, it’s parameterized in terms of a value that scales with the mean instead of the inverse. In our context, this distinction is important because it changes how we relate the distribution to the quadratic TPC in Equation 4. Specifically, to use the mortality function to determine the central tendency of the Weibull model, we invert *f* (*T*) to get *f* (*T*)^−1^, and equate this to the *median* lifetime at each temperature. This median is then used to construct the scale parameter of the Weibull distribution,

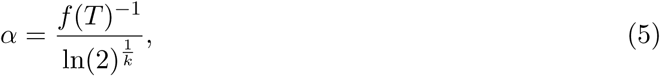

where *k* is the shape parameter. When simulating data here, we choose *k* = 3 so that lifetimes are slightly skewed but distributions are distinct from the corresponding exponential distribution. Thus, at each temperature, the data follow a Weibull distribution with a shape parameter equal to 3 and a scale parameter given in Equation 5, such that the distribution has a median of 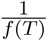.

We choose the parameters that determine the “true” quadratic function for our simulations based on a fitted mortality rate TPC from Figure B.1 (d) of Johnson et al. (2015). The quadratic was fit to *Anopheles gambiae* mortality rate data from Bayoh (2001). Using the estimated values of the parameters *a*, *T*_opt_, and *c* from this curve, the mortality rate TPC for all simulations is assumed to be

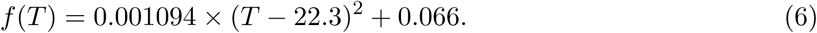

These values indicate that the optimal temperature was 22.3^◦^C with a corresponding minimum mortality rate of 0.066/day. This minimum mortality rate value gives an expected lifetime at the optimal temperature of ∼15.15 days for the exponential distribution and a median lifetime of ∼15.15 days for a Weibull distribution with *k* = 3.

### 3.2. Simulated Data

We ran four simulation experiments involving data generated from the two distributions with a quadratic mortality rate function as described in Section 3.1. The flow of these experiments from data simulation to data transformation is shown in Figure 2. 100 datasets were simulated. Each of these is composed of five unique temperature values, with 100 simulated lifetimes (e.g. as if from 100 mosquitoes) generated from the appropriate distribution at each temperature, resulting in 500 individual observations per dataset. After simulating the raw data, all lifetime values shorter than one day were truncated to one across all datasets. This reflects the assumption that lifetimes less than one day are generally unobservable in most experimental settings and that one day is the minimum observable lifetime in a physical experiment.

**Figure 2:**
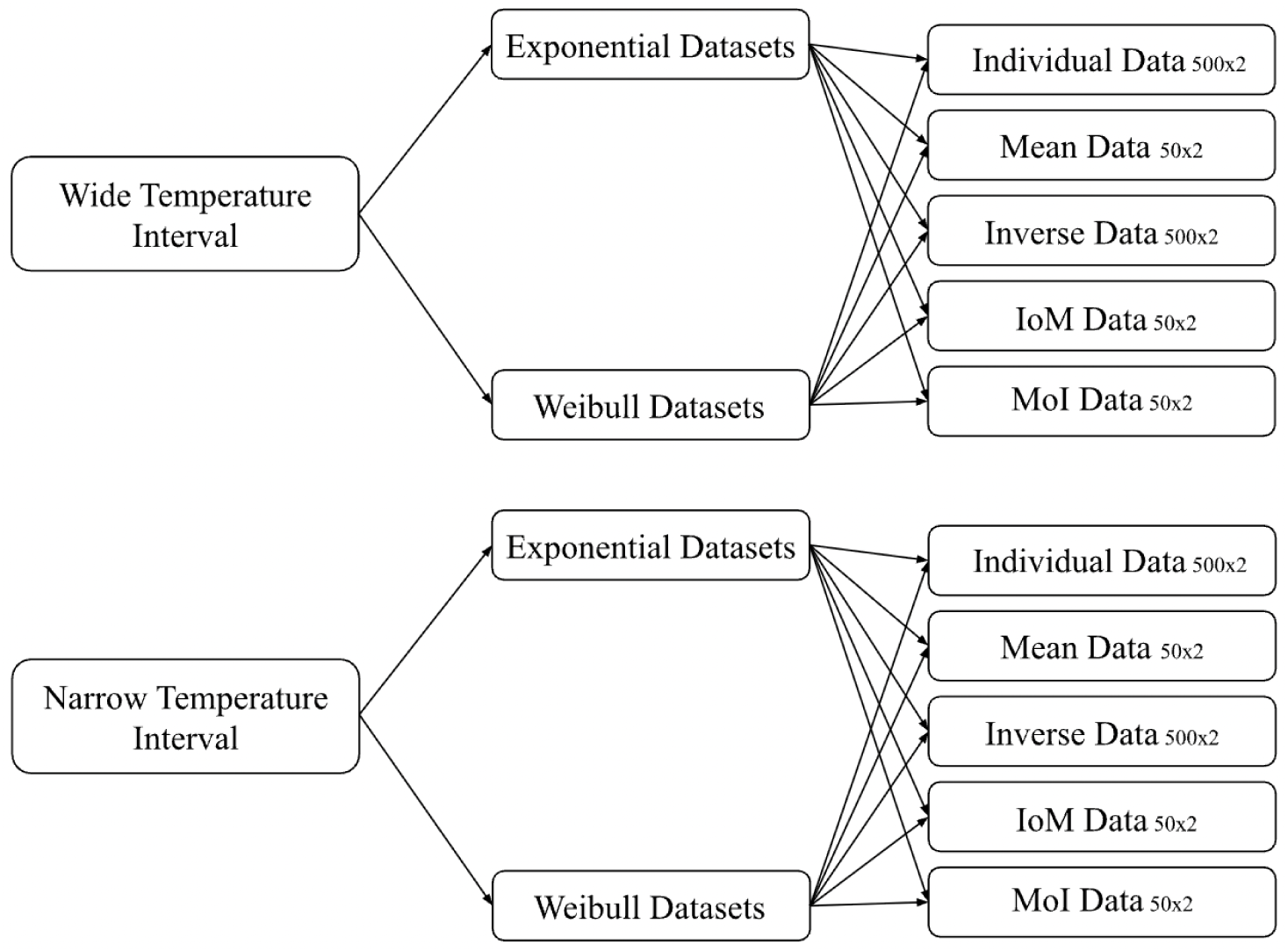
A hierarchical tree diagram showing the breakdown of the simulation experiment from temperature design to data transformation. Data is simulated assuming both a wide (top) and a narrow (bottom) temperature interval. These simulated lifetime data come from either an exponential or a Weibull distribution centered at the TPC. Thus there are four experiments run, one for each combination of temperature interval and distribution. Subsequently, each simulated dataset is summarized/transformed resulting in five possible datasets. The size of the resulting datasets after transformation is given next to the name of the transformation.

We ran simulation experiments with two virtual temperature settings: a wide temperature interval (10, 17, 21, 25, and 32^◦^C), and a narrow temperature interval (13, 17, 21, 25, and 29^◦^C). In experimental design, it is generally preferred to have a wider range of values for the independent variable, as this provides more information about the behavior of the response and decreases uncertainty in parameters. These two settings allow us to explore the impacts of this aspect of experimental design on inference and prediction.

In each simulation experiment, we additionally applied four different data transformations to each of the 100 generated datasets. This results in five distinct forms of the data (four transformations and raw data) to which we fit models.

(a) **Individual Data** uses the raw individual-level lifetime data points that have not been transformed.
(b) **Mean Data** divides the 100 observations at each of the five temperatures into 10 groups of 10, and uses the mean values of those groups to give 50 values in a single dataset.
(c) **Inverse Data** inverts the individual-level lifetime observations to model the “rate” per each individual.
(d) **Inverse of the Mean Data**, or IoM data, divides the 100 observations at each of the five temperatures into 10 groups of 10, takes the mean as before, and then inverts those values.
(e) **Mean of the Inverse Data**, or MoI data, inverts the individual-level lifetime observations, then divides the 100 observations at each of the five temperatures into 10 groups of 10 and uses the mean values of those groups.

All data summaries were created from the original datasets for each simulation to make the methods comparable. An example of these summaries of data generated from an exponential distribution is shown in Figure 3, while an example of data generated from a Weibull distribution is shown in Figure 4.

**Figure 3:**
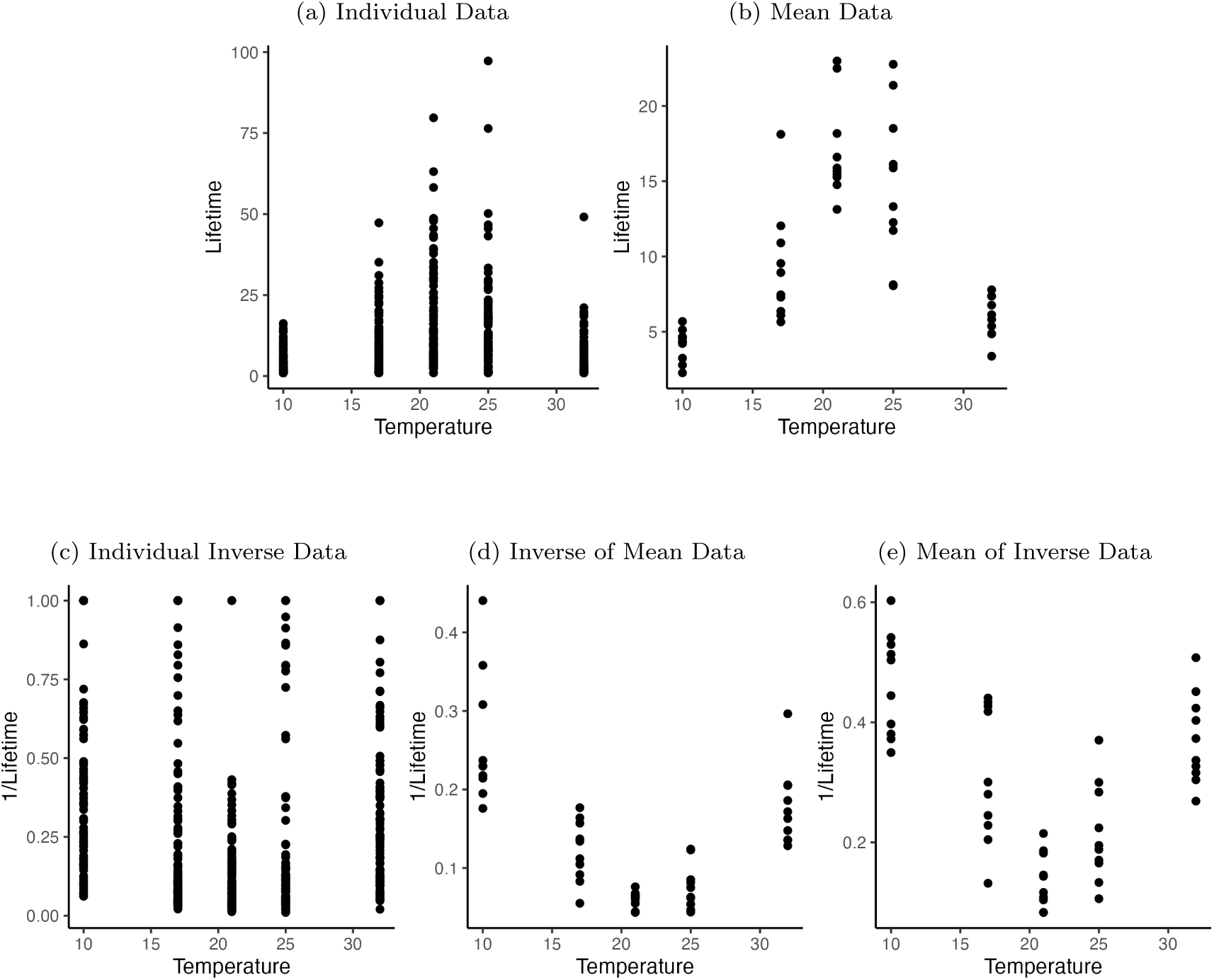
Scatterplots of the 100*^th^* dataset generated from an exponential distribution under summarization/transformation. 3a Individual data points (raw, untransformed data); 3b mean lifetime (subsets of the raw data are averaged at each temperature); 3c individual inverse data (each raw data point is inverted); 3d inverse of the mean data (i.e, values from 3b are inverted); and 3e is mean of the inverse data (i.e, values from 3c are averaged). Transformations shown are created using the same raw dataset for comparability. The raw lifetimes were truncated at 1 before any transformations.

**Figure 4:**
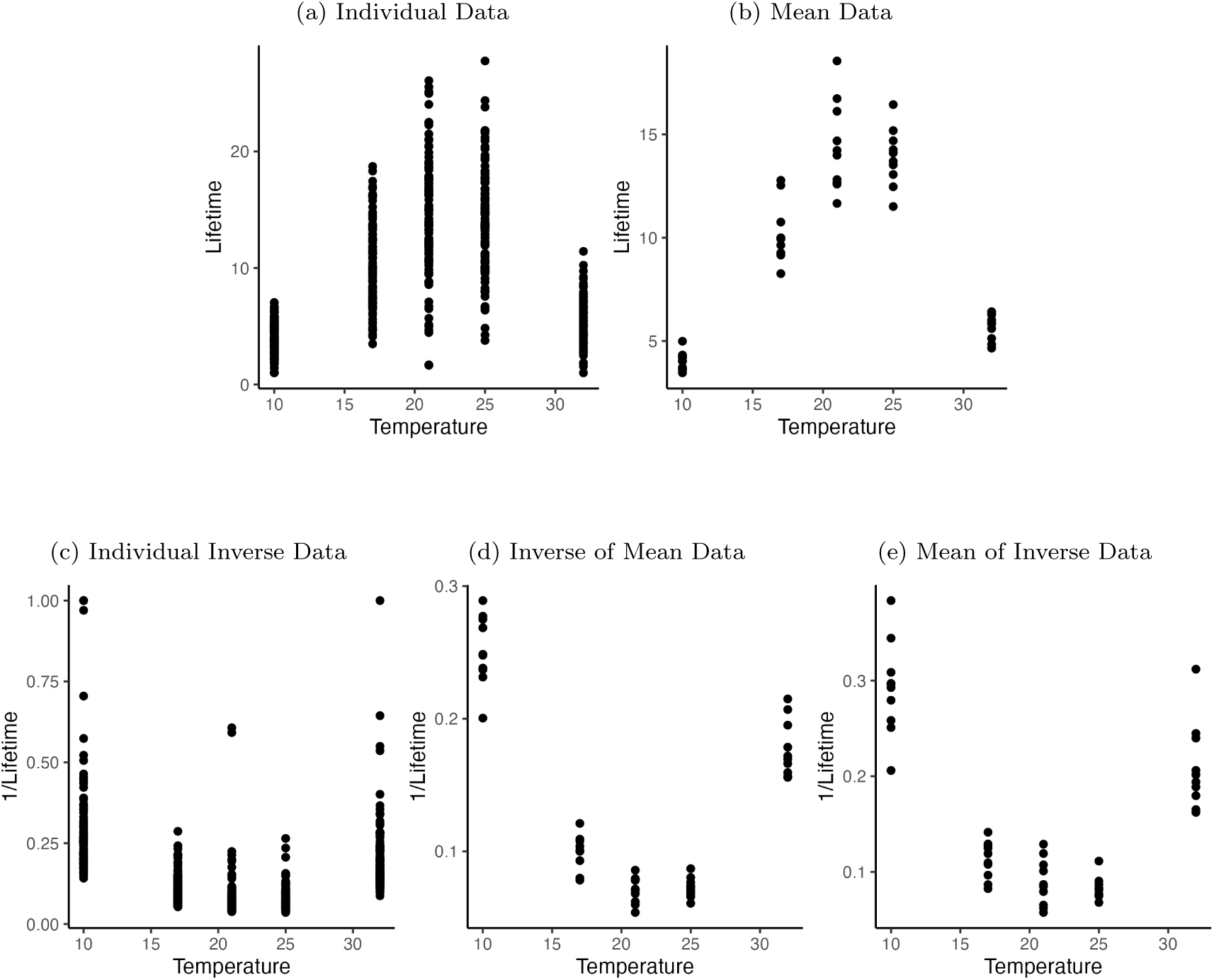
Scatter plots of the 100*^th^* dataset generated from a Weibull distribution. 4a Individual data points (raw, untransformed data); 4b mean lifetime (subsets of the raw data are averaged at each temperature); 4c individual inverse data (each raw data point is inverted); 4d inverse of the mean data (i.e, values from 4b are inverted); and 4e is mean of the inverse data (i.e, values from 4c are averaged). Transformations shown are created using the same raw dataset for comparability. The raw lifetimes are truncated at 1 before any transformations.

### 3.3. Data Modeling

After 100 datasets were generated for each of the 4 simulation experiments (wide vs. narrow intervals, Exponential and Weibull distributions), models were fit to the simulated data (including transformed data) using a Bayesian approach. We first describe the modeling for the raw individual-level data followed by assumptions for all transformed data.

Because we are interested in understanding the impact of a mismatch between the “true” distribution of the data around the TPC, for raw data from both distributions, we fit a TPC under the assumption that the underlying data-generating mechanism at each temperature was exponential,

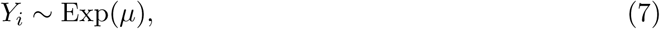

where *Y_i_* represents the response (adult lifetime in days) for observation *i*, and the rate parameter, *µ*, is the quadratic TPC in Equation 6. This approach enables us to examine the consequences of assuming an exponential data-generating process when the true underlying distribution is Weibull. For the raw datasets generated from a Weibull distribution, we also fit a TPC under the correct distribution, that is, we assumed,

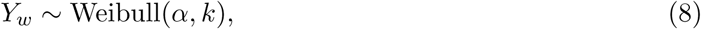

where *Y_w_* is the response at observation *w*, *k* is the shape parameter, and *α* is the scale parameter, given in Equation 5. These two approaches allow us to compare the performance of correctly specified and mis-specified data-generating mechanisms in a known TPC analysis.

For all models involving individual data, priors are shown in Table 1. All priors were chosen to be relatively noninformative; further explanations are available in the Supplementary Material. Note that because parameter *a* is small, we model the log of *a* to improve stability and convergence. Prior settings were the same for simulations using both a wide and a narrow temperature range.

**Table 1:** Priors for TPC parameters *a*, *T*_opt_, *c*, and the for the shape, *k*, when modeling the individual data. Priors are shown for data generated from both an exponential distribution, and a Weibull distribution. ∼ 𝒩 (*µ, σ*^2^) indicates a normal distribution with mean *µ* and variance *σ*^2^, while ∼ Exp(*λ*) represents an exponential distribution with a rate parameter *λ*.

| Exponential Data |  |  |  |  |  |
| --- | --- | --- | --- | --- | --- |
| Data | Model | $\log(a)$ Prior | $T_{\text{opt}}$ Prior | $c$ Prior | Shape Prior |
| Individual | Exponential | $\sim \mathcal{N}(0, 10)$ | $\sim \mathcal{N}(22, 10)$ | $\sim \text{Exp}(0.5)$ | NA |
| Weibull Data |  |  |  |  |  |
| Data | Model | $\log(a)$ Prior | $T_{\text{opt}}$ Prior | $c$ Prior | Shape Prior |
| Individual | Exponential | $\sim \mathcal{N}(0, 10)$ | $\sim \mathcal{N}(22, 10)$ | $\sim \text{Exp}(0.5)$ | NA |
| Individual | Weibull | $\sim \mathcal{N}(0, 10)$ | $\sim \mathcal{N}(22, 10)$ | $\sim \text{Exp}(0.5)$ | $\sim \text{Exp}(0.5)$ |

As discussed in Section 2, data that are transformed will not have the same underlying distribution as the raw data. Further, it is common to assume that trait data are approximately normal (or truncated normal to ensure positive traits), especially as this approximately corresponds to the assumptions of NLLS. Thus, for both the experiments using exponential data and the experiments using Weibull data, when using any transformations (i.e. the mean data, the individual inverse data, the IoM data, or the MoI data), the distribution of the data is assumed to be a normal distribution truncated at 0 (to avoid negative lifetimes or rates). That is, we assume that

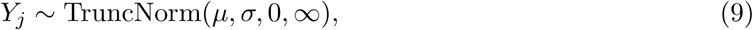

where *Y_j_* represents the response for observation *j* and *σ* is the estimated standard deviation. When using the mean data, we are modeling the lifetimes directly, and *µ* is the inverse of the quadratic function in Equation 6 (i.e. the lifetime curve). When modeling the “rate”, or any summary of the inverse of the lifetimes, *µ* is given by Equation 6. Although the mean of exponential random variables follows a Gamma distribution (see Supplementary Material), in our work, we use a truncated normal model to recreate common models and analyses that most researchers are currently doing (Shi and Ge, 2010; Rezende and Bozinovic, 2019). Using a normal model is the closest equivalent to using nonlinear least squares, which is a very popular technique when fitting TPCs. We then truncate this model so that predicted lifetimes cannot be less than 0.

When modeling the transformed data, the priors on parameters log(*a*) and *T*_opt_ remained the same as the individual-level data. The prior on parameter *c* differs between models, but was chosen to ensure that the values of *c* were positive and to reach convergence with biologically realistic values. Further explanation on the choice of priors can be found in the Supplementary Material. These priors are shown in Table 2. Settings were the same for the simulations using both wide and narrow temperature ranges.

**Table 2:** Priors for parameters *a*, *T*_opt_, *c*, and *σ* when modeling the transformed data. Priors are shown for data generated from both an exponential distribution, and a Weibull distribution. ∼ 𝒩 (*µ, σ*^2^) indicates a normal distribution with mean *µ* and variance *σ*^2^, ∼ Exp(*λ*) represents an exponential distribution with a rate parameter *λ*, and ∼ Gamma(*ã*,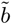) represents a gamma distribution with shape parameter *ã* and rate parameter 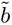 respectively.

| **Exponential Data** |  |  |  |  |
| --- | --- | --- | --- | --- |
| **Data** | **$\log(a)$ Prior** | **$T_{\text{opt}}$ Prior** | **$c$ Prior** | **$\sigma$ Prior** |
| Mean | $\sim \mathcal{N}(0, 10)$ | $\sim \mathcal{N}(22, 10)$ | $\sim \text{Exp}(0.5)$ | $\sim \text{Exp}(0.5)$ |
| Individual Inverse | $\sim \mathcal{N}(0, 10)$ | $\sim \mathcal{N}(22, 10)$ | $\sim \text{Gamma}(10, 1)$ | $\sim \text{Exp}(0.5)$ |
| Inverse of Mean | $\sim \mathcal{N}(0, 10)$ | $\sim \mathcal{N}(22, 10)$ | $\sim \text{Exp}(0.5)$ | $\sim \text{Exp}(0.5)$ |
| Mean of Inverse | $\sim \mathcal{N}(0, 10)$ | $\sim \mathcal{N}(22, 10)$ | $\sim \text{Gamma}(10, 1)$ | $\sim \text{Exp}(0.5)$ |
| **Weibull Data** |  |  |  |  |
| **Data** | **$\log(a)$ Prior** | **$T_{\text{opt}}$ Prior** | **$c$ Prior** | **$\sigma$ Prior** |
| Mean | $\sim \mathcal{N}(0, 10)$ | $\sim \mathcal{N}(22, 10)$ | $\sim \text{Exp}(0.5)$ | $\sim \text{Exp}(0.5)$ |
| Individual Inverse | $\sim \mathcal{N}(0, 10)$ | $\sim \mathcal{N}(22, 10)$ | $\sim \text{Exp}(0.5)$ | $\sim \text{Exp}(0.5)$ |
| Inverse of Mean | $\sim \mathcal{N}(0, 10)$ | $\sim \mathcal{N}(22, 10)$ | $\sim \text{Exp}(0.5)$ | $\sim \text{Exp}(0.5)$ |
| Mean of Inverse | $\sim \mathcal{N}(0, 10)$ | $\sim \mathcal{N}(22, 10)$ | $\sim \text{Exp}(0.5)$ | $\sim \text{Exp}(0.5)$ |

### 3.4. Common model specifications

#### Algorithm 1

Data Generation and Model Fitting

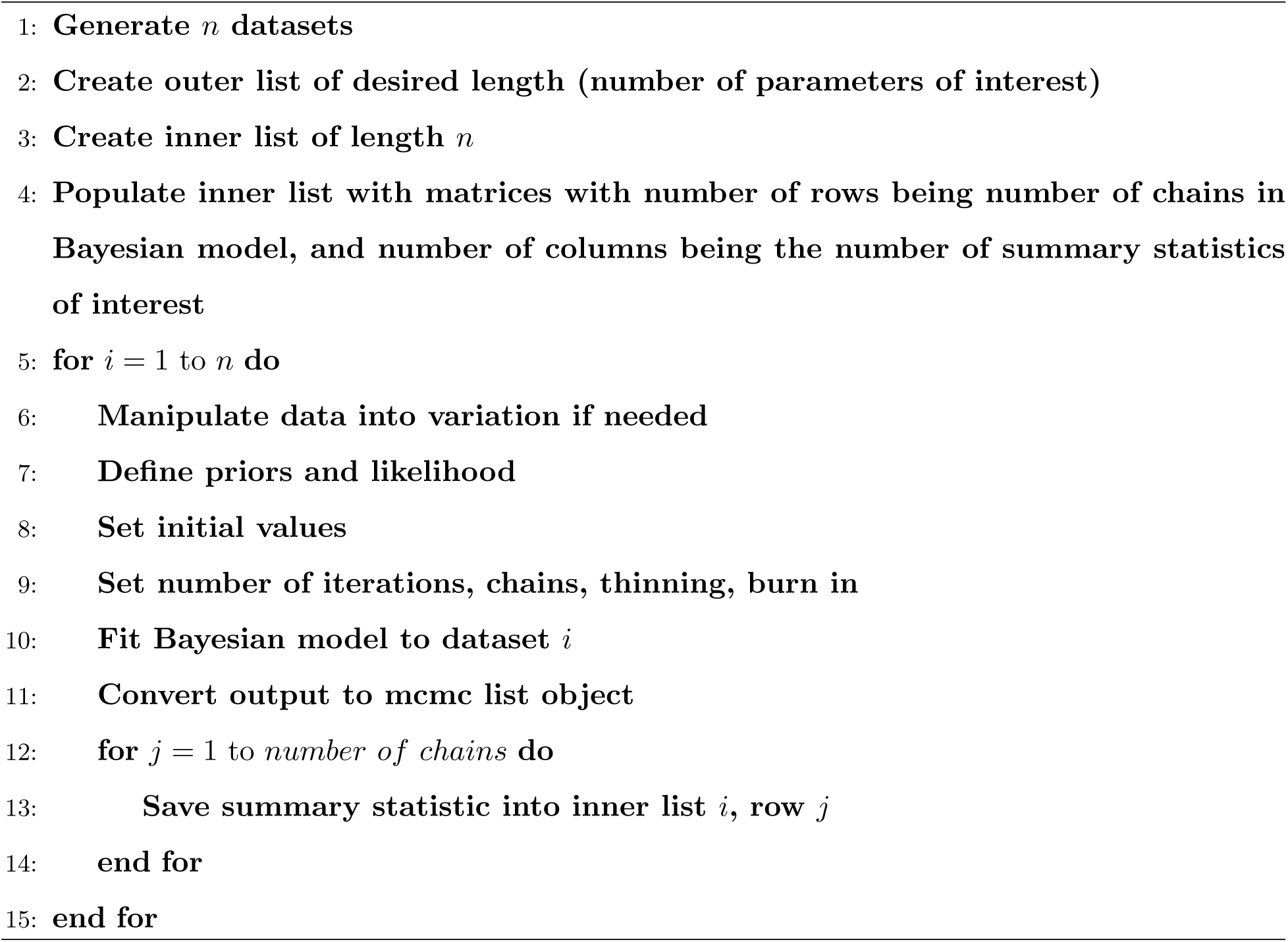

In our simulation experiments, the general algorithm as well as many of the settings remained consistent between all models, regardless of transformation or distribution. The general procedure for modeling using these generated datasets is shown below in Algorithm 1. The number of generated data sets, *n*, was 100, the parameters of interest included *a*, *T*_opt_, *c*, and the deviance, as well as sigma (*σ*) when using a truncated normal model or shape (*k*) when using a Weibull model. The summary statistics of interest were the posterior mean, posterior median, and the lower and upper highest density interval (HDI) bounds of the posterior samples. For each experiment, we ran 5 Markov chains, each with 60,000 iterations, dropping 30,000 burn-in, and using a thinning number of 8. This resulted in 18,750 posterior samples for each fitted dataset and transformation combination. These settings were the same for all simulation experiments. For each chain in each model, the initial values for all parameters were generated randomly from their respective prior distributions to help check for convergence among chains. Bayesian models were fit to the datasets using `JAGS/rjags` (Su and Yajima, 2024; Plummer, 2025) in R (R Core Team, 2025). Convergence was checked visually using trace plots for a randomly chosen subset of runs (∼ 5 % of total chains) for each experiment. Examples of these trace plots are given in the Supplementary Material.

### 3.5. Comparison methods

Within each experiment, we take multiple approaches to understand and compare the results of the different models. We first make comparisons between models based on a single dataset, including visualizations of the TPC using joint posterior samples of the three parameters, estimation of the peak of the curve (i.e. the optimal temperature (*T*_opt_) and the corresponding maximum lifetime (*L*_Max_)), and estimation of the tails of the curve (i.e. the upper (*T*_Upper_) and lower (*T*_Lower_) critical temperatures at which lifetime is predicted to be less than one day). The joint posterior samples of parameters *a*, *T*_opt_, and *c* came from a single chain in the 100*^th^* model fit (corresponding to the 100*^th^* simulated dataset). Each joint posterior sample, consisting of one value for each of the three parameters, was then used to evaluate the quadratic function at the relevant values of interest. For visualizing the joint posterior samples, the quadratic function was evaluated across a temperature range from 5^◦^C to 37^◦^C, while the peak and tails were evaluated according to their definitions described above. When estimating *T*_Upper_ and *T*_Lower_ for the individual inverse, IoM, and MoI data, we inverted the estimated mortality rate curves (as we modeled mortality rate directly) to obtain the corresponding lifetime curves needed for these calculations (visuals provided in the Supplementary Material).

We also performed three sets of comparisons across all 100 datasets within each simulation experiment as obtained from fitting to the individual or transformed data. The first comparison is between the uncertainty in estimating *T*_opt_, *L*_Max_, *T*_Upper_, and *T*_Lower_. For the values of *T*_opt_ and *L*_Max_, we define uncertainty as the length of the 95% HDI interval of the posterior samples. For *T*_Upper_ and *T*_Lower_, we define uncertainty as the length of the boxplot whiskers derived from their estimated values. For each modeling method, the median length is calculated and then compared. Although we visualize the spread of these estimations for a single dataset, we calculate the uncertainty across all 100 datasets using a similar sampling and evaluation method as described previously, but including all chains from all models. A full comparison of uncertainties is available in the Supplementary Material.

The second comparison is between the proportion of individual parameter coverage. For the three parameters (*a*, *T*_opt_, and *c*), we calculated the 95% HDI bounds of the posterior samples across each of the five chains in each of the 100 models. We then determined the proportion of these intervals that contained the true parameter value.

The third comparison is the range of RMSEs across the 100 models, calculated by comparing the true curve with the curves generated from joint posterior samples of the three parameters. For each of the 100 models within each method, we used the same joint posterior sampling approach described previously for single-model comparisons. We then calculate the RMSE between the true curve from Equation 6 and the estimated curve from each individual posterior sample, evaluated over a temperature range from 5^◦^C to 37^◦^C.

## 4. Results

Going forward, we focus on the models from the experiments using a wide temperature interval. Any differences in results or patterns between the wide temperature interval and the narrow temperature interval are noted in Section 4.3, and additional results are shown in the Supplementary Materials.

### 4.1. Exponential data

First we explore the results of the simulations based on data that is exponentially distributed around the TPC. The curves resulting from the joint posterior evaluations are visualized in Figure There is a panel for each of the five data variations. Each panel includes a median curve based on the posterior samples, with the true data-generating curve shown as a dashed line for comparison. The uncertainty in the estimates at each temperature is represented as the ribbon bounded by the 95% HDI bounds. When modeling with the individual or mean data, we are modeling the lifetimes, so the resulting curve is the lifetime curve. When using a form of inverted data, we are attempting to model the mortality rate directly, so the curve presented reflects this. The corresponding lifetime or inverse lifetime values from the raw or transformed dataset used to fit the model are included in each plot for reference. For the selected example dataset, using the individual data, the mean data, and the IoM data result in median curves similar to the true curve, with the smallest uncertainty. Although the curve created by using the MoI data is quite far from the true curve, we can see that it fits the data used to fit the model. That is, the bias in the parameter inference is caused by the bias introduced by this transformation.

To visualize the capacity of the model to capture the peak of the TPC, the 50%, 75%, and 95% HDI bounds of the posterior samples from the marginal distribution of *T*_opt_ are given in Figure 6a. For this particular dataset, the true value of *T*_opt_ is within the 50% or 75% bands for each method. However, differences arise in the uncertainty (calculated from all models). When using the individual inverse data, the uncertainty was almost 2.5 times that of the raw individual-level data. Figure 6b shows the visualization of the same HDI bounds for the estimated maximum lifetime at the optimal temperature. Using both the mean and the MoI data failed to capture the true maximum lifetime in the 95% bounds. Across all datasets, the uncertainty in estimates using the individual inverse data was more than six times larger than that of the individual data, while the uncertainty when using the IoM data was around 3.7 times larger.

**Figure 5:**
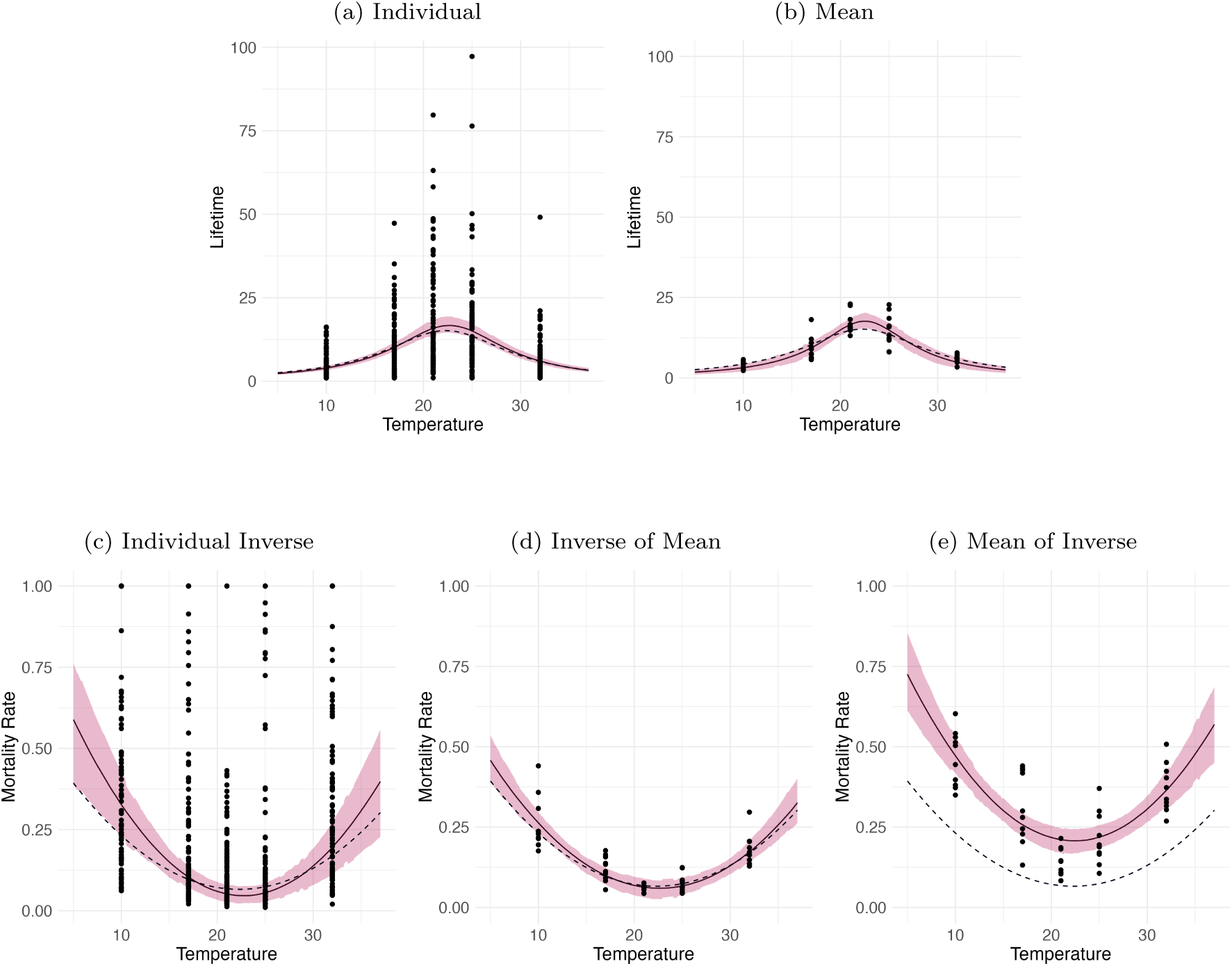
Comparison of the fitted TPC (median lifetime or mortality rate) when using data generated from an exponential distribution with a wide temperature interval across data transformations. 5a Individual data points; 5b mean data; 5c individual inverse data; 5d IoM data; and 5e MoI data. Curves were made by evaluating samples of the posterior function across a temperature range from 5^◦^C to 37^◦^C. The median response at each temperature is shown as the solid lines, while the true curve is a dashed line. The 95% HDI bounds of the response at each temperature (TPC) were also plotted as the pink ribbon. The corresponding transformed data used to fit the TPC is included.

**Figure 6:**
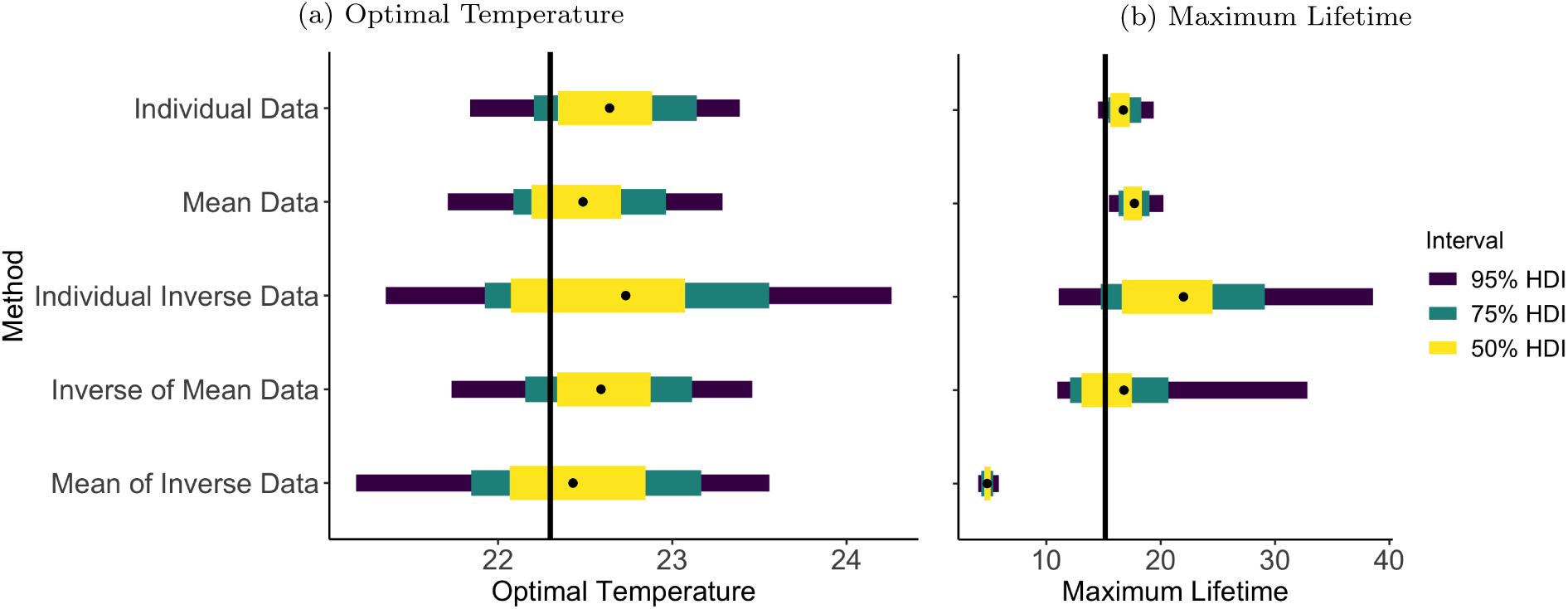
Posterior estimates and uncertainty of ^(^6a^)^ the estimated optimal temperature, *T_opt_* and ^(^6^b^) of the maximum lifetime, *L_Max_*, across data transformations for the single exponential dataset visualized in Figure 5. The vertical black lines represent the true values, the black dot represents the median of the posterior samples of each method, and the yellow, teal, and purple bars represent the 50%, 75%, and 95% HDI bounds, respectively.

The boxplots of the estimated values of *T*_Upper_ vs *T*_Lower_ are shown in Figure 7. Using the MoI data was the only method that did not contain either of the true values within the whiskers of its boxplot. Using the individual inverted data increased the width of the whisker-based uncertainty bands of both the upper and lower critical temperatures by ∼2.5 times the width of the bands when individual-level raw data were used to estimate the TPC. Additionally, transforming the data introduced directional bias. The lower critical temperature was overestimated, while the upper was underestimated, magnifying the size of the error in the same direction as the error when using the individual level data (that is, error due to sampling). Among the transformations, the MoI data introduced the most bias, while models using individual inverse and IoM data showed less bias. Note that the values and uncertainties from estimating the peak and tails of the curve apply only to the single dataset used to create the figures. The direction and magnitude of bias will reflect the particular sample for any fitted dataset.

**Figure 7:**
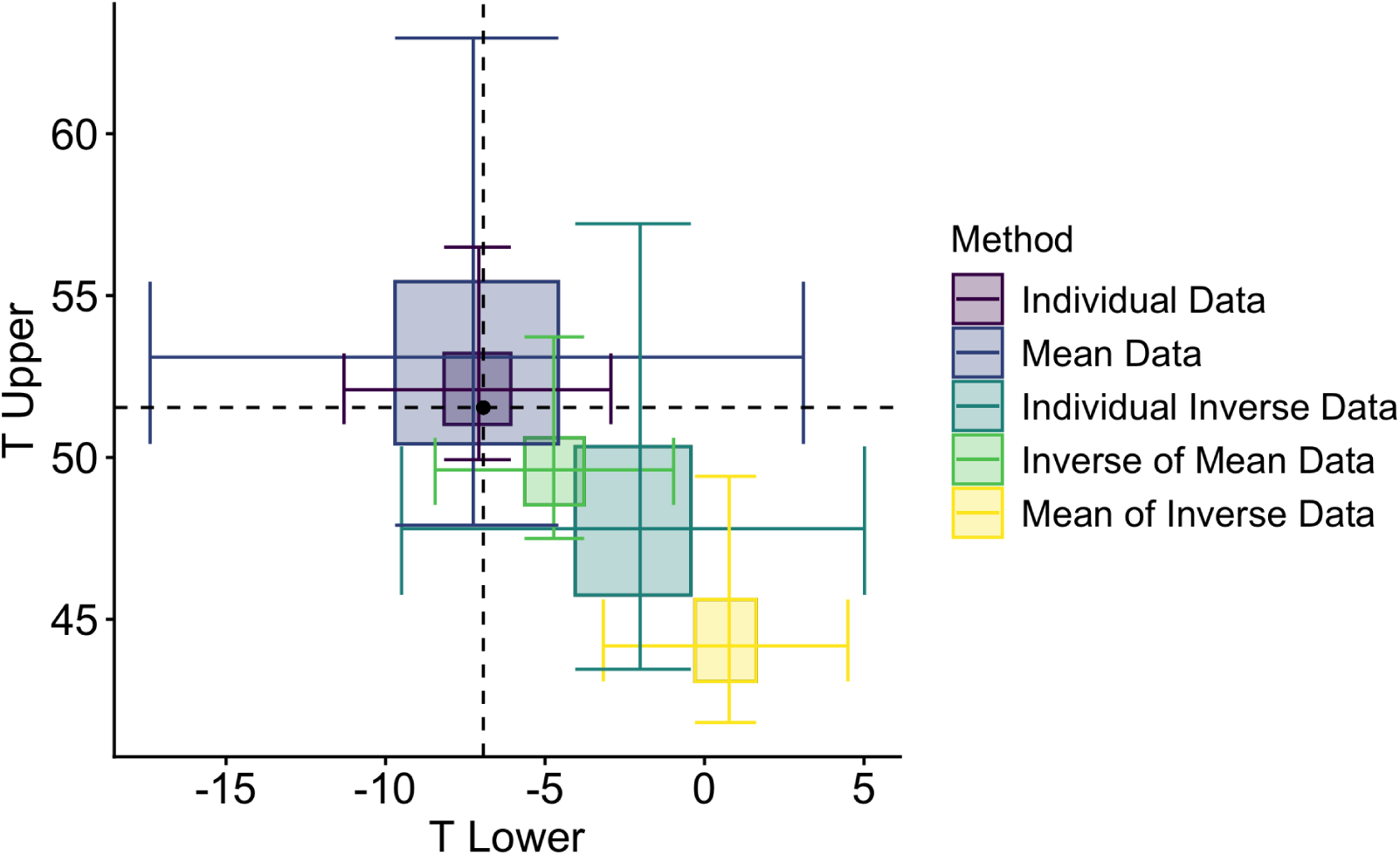
Boxplot summarizing the joint posterior distribution of the estimated lower (*T*_Lower_) and upper (*T*_Upper_) critical temperatures (where lifetime is less than one day) for the different data transformations when using exponential data with a wide temperature interval. The black point at the intersection of the dashed lines represents the true values, derived from Equation 6. Outliers not shown.

Next, we compare the fits using summaries across the 100 simulated datasets. First we explore the coverage of the marginal posterior distributions for each TPC parameters for the five different data transformations/fits (Table 3). Ideally, we would expect the proportions to be close to 0.95, indicating that the 95% HDIs accurately reflect the uncertainty in the parameter estimates without being too conservative. Notable results include a lower proportion of coverage of parameter *a* for all methods using a form of inverted data, as well as the fact that modeling using the MoI data was not able to capture the true value of parameter *c* in any chain in any of the 100 simulated data sets. As expected given the results for the single dataset explored above, using the individual data gave good coverage for all parameters. The mean data performed nearly as well, except for lower coverage of the minimum mortality rate *c* (and thus the maximum lifetime).

**Table 3:**
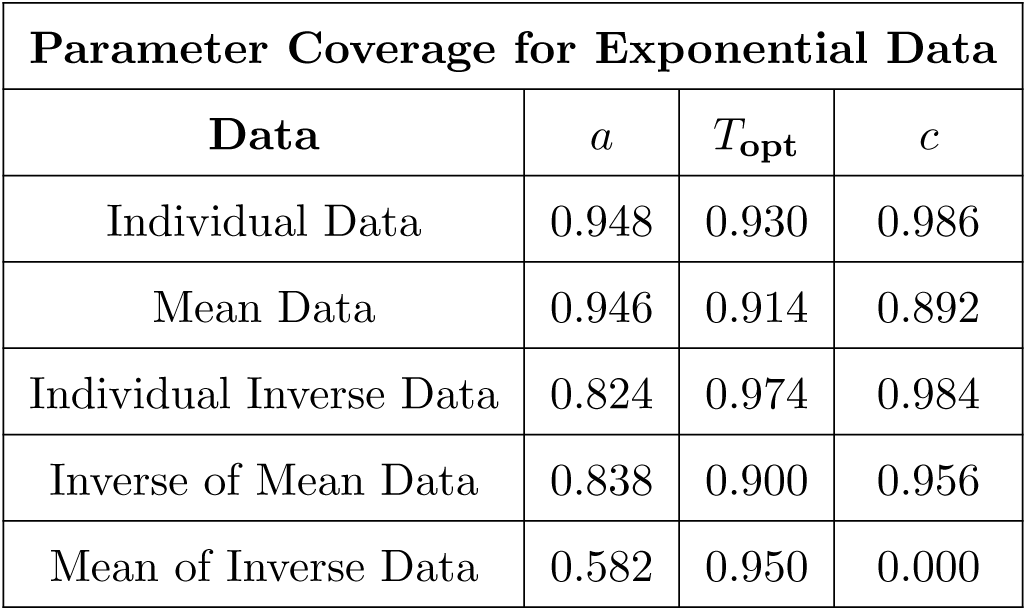
Proportion of coverage for parameters *a*, *T*_opt_, and *c*. The coverage is calculated as the proportion of chains (5 chains in each model, 100 models for each data variation, 500 chains total) in which the true value of the parameter lay within the 95% HDI bounds of the posterior samples.

Finally we examined the RMSE across all 100 simulated datasets. Recall that the RMSE is calculated as the distance between the posterior mean prediction and the mean of the data at each temperature. The resulting distribution of RMSE values from each method are shown in Figure 8. The average RMSE across all 100 models, as well as the 95% upper and lower HDI bounds are given. Since all methods had a lower bound close to zero, a logarithmic scale was used in order to better differentiate between the results. Among the methods, the individual data produced the lowest average RMSE, followed by the IoM data. The method using the mean data had the lowest individual RMSE values, but surprisingly also the highest (except for the MoI). MoI performed the most poorly, congruent with the other comparisons above.

**Figure 8:**
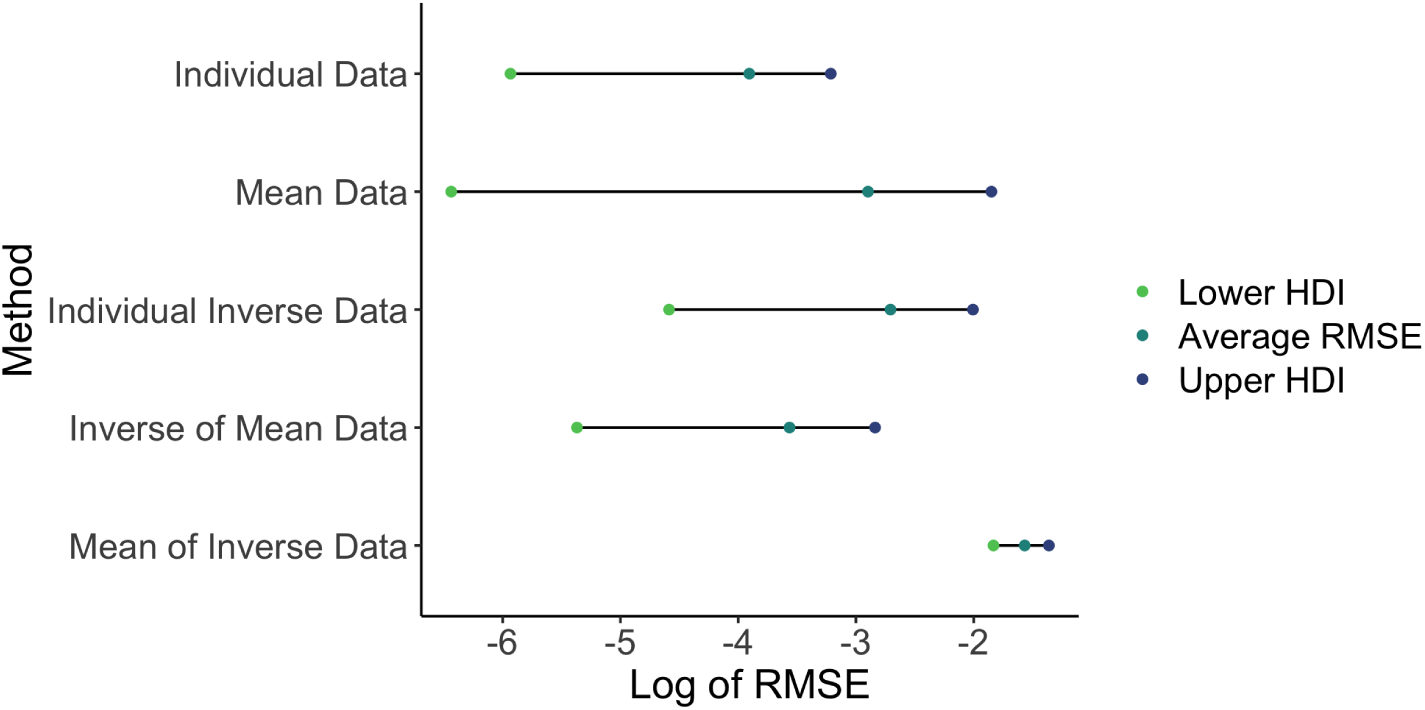
Posterior summaries of log(RMSE) between posterior estimates of the TPC and the true TPC across data transformations for all simulated datasets from the wide temperature experiment, exponentially distributed data. For each modeling method, the average RMSE and the upper and lower 95% HDI bounds of the 100 simulated datasets are shown. RMSE is calculated using all thinned posterior samples taken from chain one of each fit, calculating the posterior TPC at 250 temperatures from 5 to 37^◦^C, calculating the error between that curve to the true TPC, and then calculating RMSE. The plot uses a logarithmic scale to make the lower bounds easily distinguishable.

### 4.2. Weibull Data

Next, we evaluate the performance of the methods when the data come from a Weibull distribution around the TPC. Note that we have one additional model/data combination - the individual-level data gets modeled with both an exponential and a Weibull distribution here. The posterior distributions of the TPC for the single selected simulated data set are shown in Figure 9. Each panel again displays the median curve, the true curve, and the uncertainty in estimates for one of the methods. We see that using the individual data with a Weibull model, the mean data, and the IoM data produce curves close to the true curve with the least uncertainty for this dataset. However, the true curve is not uniformly contained in the uncertainty bounds for the IoM method. This is in contrast to the individual data with an exponential model where the estimated curve is close to the true model but the mound of uncertainty is high.

**Figure 9:**
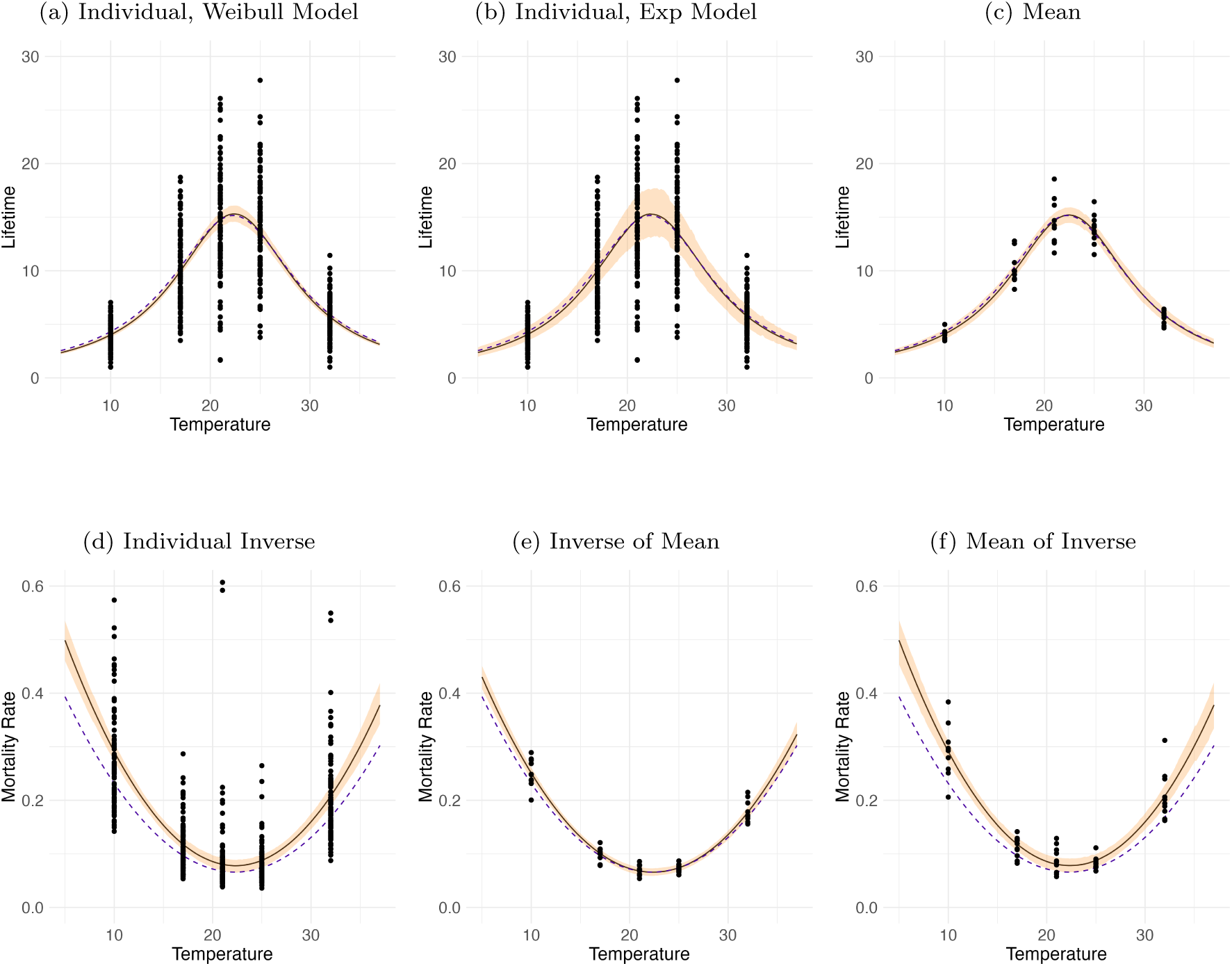
Comparison of the fitted TPC (median lifetime or mortality rate) when using data generated from a Weibull distribution with a wide temperature interval across data transformations. 9a Individual data points with a Weibull model; 9b individual data points with an exponential model; 9c mean data; 9d individual inverse data; 9e IoM data; and 9f MoI data. Curves were made by evaluating samples of the posterior function across a temperature range from 5^◦^C to 37^◦^C. The median response at each temperature is shown as the solid lines, while the true curve is a dashed line. The 95% HDI bounds of the response at each temperature (TPC) were also plotted as the orange ribbon. The corresponding transformed data used to fit the TPC is included.

In Figures 10a and 10b, we show marginal posterior plots of *T*_opt_ and *L*_Max_, respectively, to compare the estimated uncertainty in the peak of the curve across modeling approaches for the single dataset. In this dataset, the true values of both *T*_opt_ (Figure 10a) and *L*_Max_ (Figure 10b) are within at least the 95% bands for each method. When comparing the uncertainty across datasets, using the individual data with an exponential model resulted in approximately three times the uncertainty than using the individual data with a Weibull model for both values, and the uncertainty in parameters was greater for this method for both parameters than almost all other approaches. As we saw in the exponential data example, here the maximum lifetime estimate is least reliable for MoI approach.

**Figure 10:**
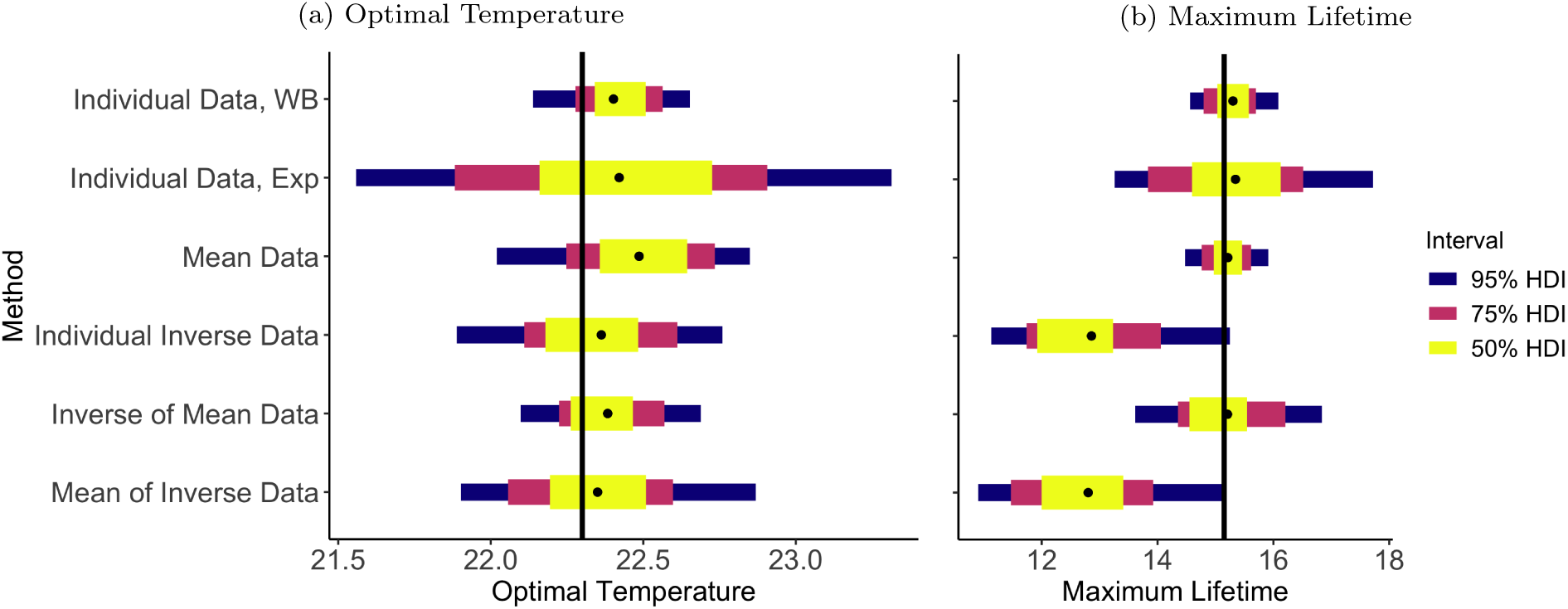
Posterior estimates and uncertainty of ^(^10a^)^ the estimated optimal temperature, *T_opt_* and ^(^10b^)^ of the maximum lifetime, *L_Max_*, across data transformations for the single Weibull dataset visualized in Figure 9. The vertical black lines represent the true values, the black dot represents the median of the posterior samples of each method, and the yellow, teal, and purple bars represent th 50%, 75%, and 95% HDI bounds, respectively.

The posterior estimates of *T*_Upper_ and *T*_Lower_ are visualized using boxplots in Figure 11. The models using the individual inverse data, IoM data, and MoI data did not capture the true values of the lower critical temperature. Of the three, only the model using the IoM data was able to capture the true value of the upper critical temperature within its whiskers. Among the methods that successfully captured the true value of *T*_Lower_, the smallest uncertainty comes from using the individual-level data with a Weibull model. Similar to using exponential data, we can see a directional bias with the lower critical temperature being overestimated and the upper being underestimated when using some of the data transformations. The individual inverse and MoI data exhibit the most bias.

**Figure 11:**
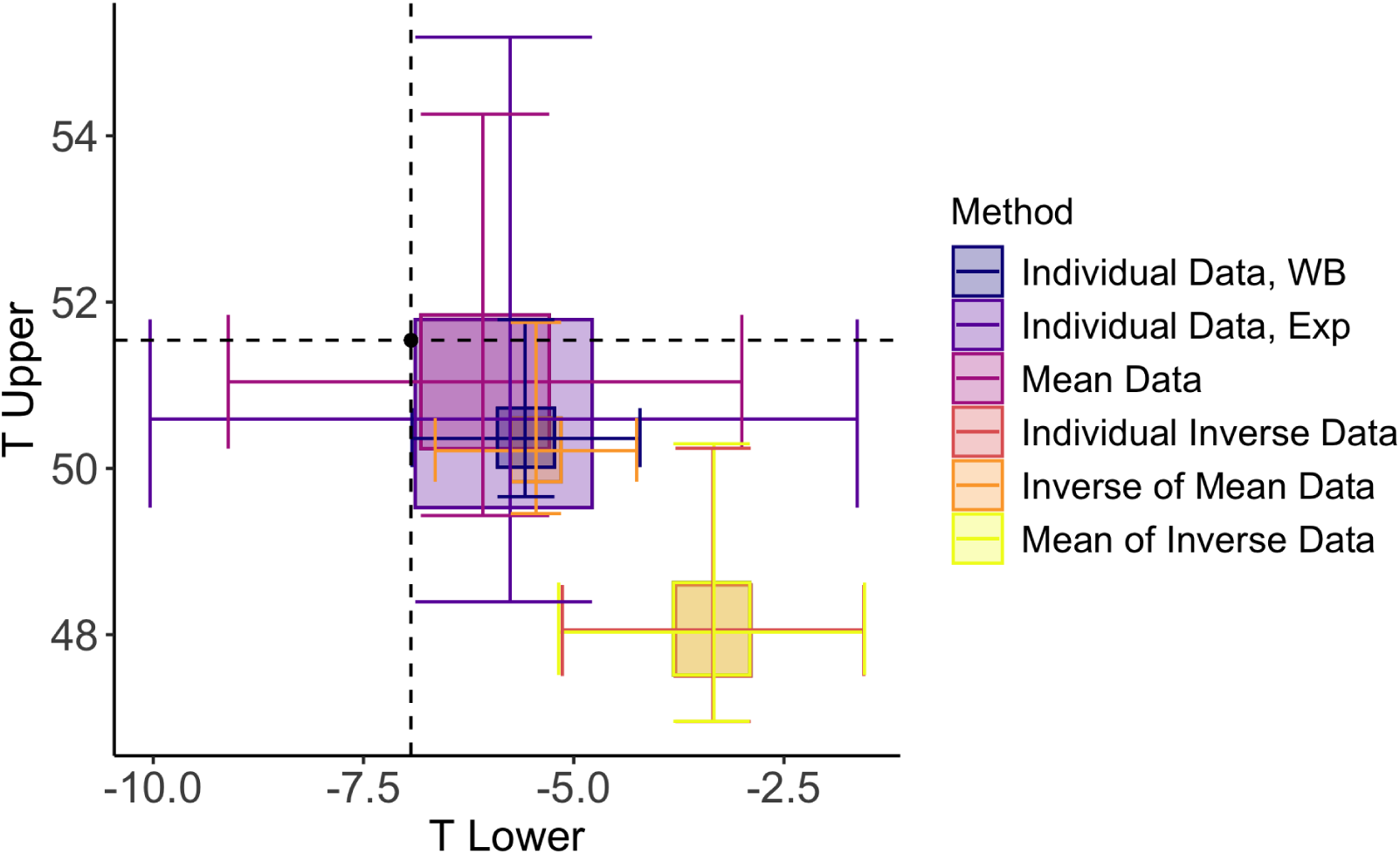
Boxplot summarizing the joint posterior distribution of the estimated lower (*T*_Lower_) and upper (*T*_Upper_) critical temperatures (where lifetime is less than one day) for the different data transformations when using Weibull data with a wide temperature interval. The black point at the intersection of the dashed lines represents the true values, derived from Equation 6. Outliers not shown.

We find similar patterns across the 100 simulated datasets as we did for the exponentially distributed data. In Table 4, we show the parameter coverage, with the inclusion of the shape parameter (*k*) when modeling with a Weibull distribution. Again, we would expect the proportion to be approximately 0.95. Notably, coverage is low for parameters *a* and *c* when using the individual inverse and the MoI data. The temperature at the optimum is more reliably within the 95% HDI bounds. In a few cases, the coverage is equal to 1, indicating conservative intervals; these intervals may be wider than necessary, but they consistently contain the true value.

**Table 4:** Proportion of coverage for parameters *a*, *T*_opt_, *c*, and *k* (shape). The coverage is calculated as the proportion of chains (5 chains in each model, 100 models for each data variation, 500 chains total) in which the true value of the parameter lay within the 95% HDI bounds of the posterior samples. “Individual Data*_WB_* ” represents the individual data with an Weibull model while “Individual Data*_Exp_*” represents the individual data with an exponential model.

| Parameter Coverage for Weibull Data |  |  |  |  |
| --- | --- | --- | --- | --- |
| Data | $a$ | $T_{\text{opt}}$ | $c$ | $k$ |
| Individual Data<sub>WB</sub> | 0.914 | 0.950 | 0.928 | 0.950 |
| Individual Data<sub>Exp</sub> | 1.000 | 1.000 | 1.000 | NA |
| Mean Data | 0.994 | 0.926 | 0.880 | NA |
| Individual Inverse Data | 0.374 | 0.938 | 0.258 | NA |
| Inverse of Mean Data | 0.858 | 0.950 | 1.000 | NA |
| Mean of Inverse Data | 0.406 | 0.932 | 0.294 | NA |

We also compare the (log) RMSE across all of the simulated datasets (Figure 12) The individual data modeled with a Weibull distribution produced the lowest average RMSE, followed by the IoM, the mean, and the individual data modeled with an exponential distribution. These three had relatively similar performance by this metric. This partly reflects the metric itself (which is calculated strictly based on the mean data and the estimated median posterior TPC). As with the exponential data, the other two inverse methods perform poorly by this metric.

**Figure 12:**
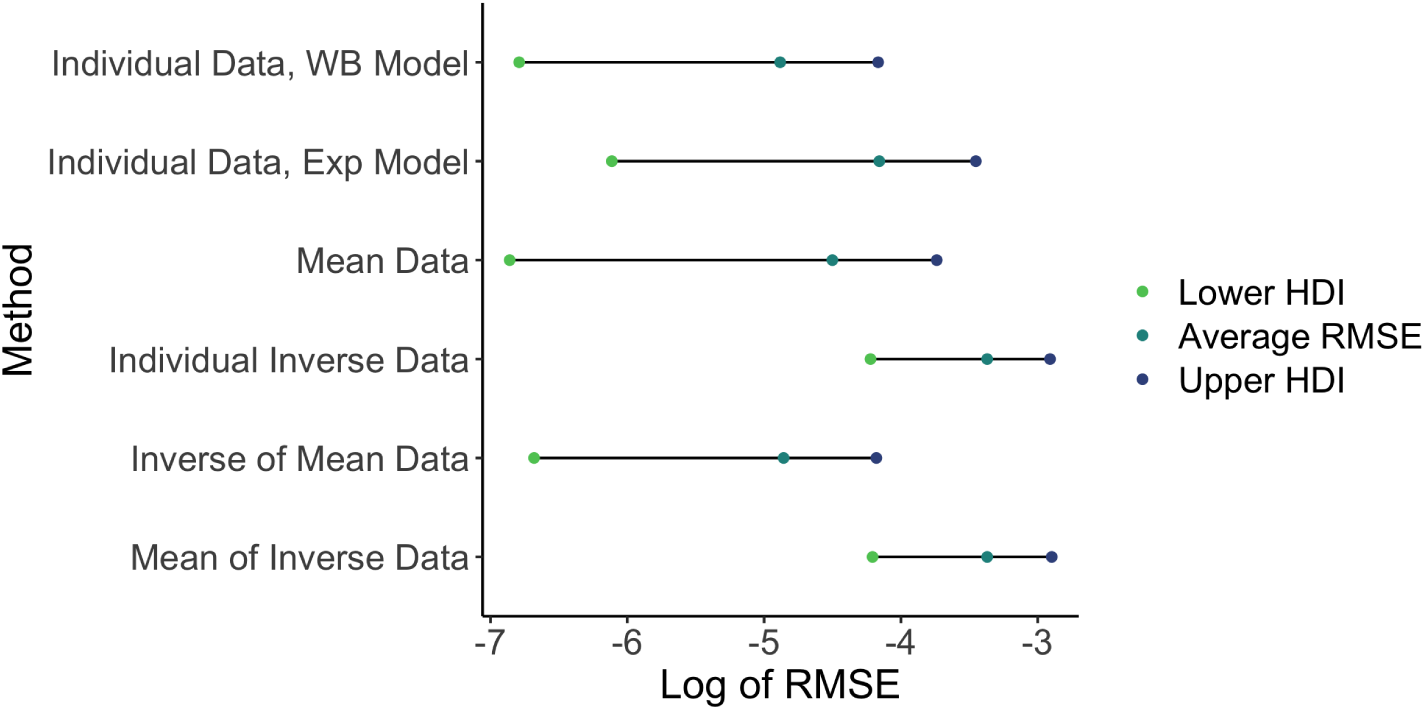
Posterior summaries of log(RMSE) between posterior estimates of the TPC and the true TPC across data transformations for all simulated datasets from the wide temperature experiment, data distributed according to a Weibull distribution. For each modeling method, the average RMSE and the upper and lower 95% HDI bounds of the 100 simulated datasets are shown. RMSE is calculated using all thinned posterior samples taken from chain one of each fit, calculating the posterior TPC at 250 temperatures from 5 to 37^◦^C, calculating the error between that curve to the true TPC, and then calculating RMSE. The plot uses a logarithmic scale to make the lower bounds easily distinguishable.

### 4.3. Wide vs. Narrow Temperature Interval

When comparing across all models fit to simulated exponential data, estimating *T*_opt_, *L*_Max_, *T*_Upper_, and *T*_Lower_ using the narrow temperature range generally resulted in greater uncertainty in estimates of critical temperatures, particularly when estimating *T*_Upper_ and *T*_Lower_, or the tails of the curve. Coverage comparisons show that the narrow temperature interval resulted in more parameters reaching coverage close to one, suggesting that the intervals may be wider than necessary due to increased uncertainty in all parameters.

Comparing visuals and estimates from a *single* simulated Weibull-generated dataset, models using a narrow temperature interval struggled to capture the true value of *L*_Max_ within the inner HDI bounds; the individual inverse data failed to capture the true value entirely. However, when estimating the tails of the curve, narrow interval models actually outperformed those using the wider interval. Both the individual inverse and MoI methods captured the true *T*_Upper_, whereas these methods using a wide temperature interval did not. The IoM data was able to capture the true values of both *T*_Upper_ and *T*_Lower_ when using a narrow temperature interval, compared to only the upper critical temperature when using a wide temperature interval. However, across the 100 models, the uncertainty of these values was typically larger in models made from a narrow temperature interval, particularly at the tails of the curve. This indicates that the results from the single dataset are non-representative. Most of the coverage proportions were similar between the two temperature intervals. One large difference is that when using a form of inverted data, parameter *a* had better coverage with a narrow interval, while parameter *c* had worse coverage.

In Figures 13a and 13b, we show the range of log RMSEs for each model under both the narrow and wide temperature intervals using exponential and Weibull data, respectively. We can see that generally, models using the wider temperature interval tended to have lower average RMSEs and HDI bounds, as well more narrow HDI bounds. Thus with a wider temperature range, not only did almost all of the models perform better, but they were less likely to perform poorly on a particular dataset. This is consistent with statistical theory about the role of the range of predictors in determining uncertainty and predictive accuracy in regression contexts.

**Figure 13:**
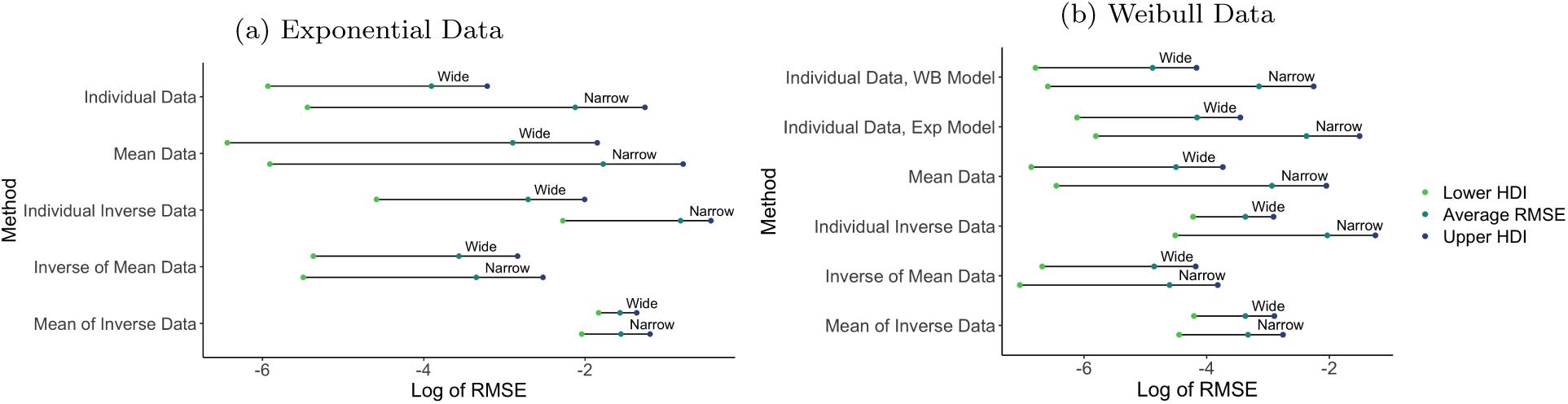
RMSE comparisons between a narrow and a wide temperature interval for data simulated from an exponential distribution (13a) and a Weibull distribution (13b). See caption of Figure 8 or 12 for description of RMSE calculation.

## 5. Discussion

In this paper we explored the coupled effects of the type of distribution of data around a TPC together with different approaches to summarizing and modeling the data on the ability to reconstruct a known TPC. More specifically, we included simulated data from two possible generating distributions around the TPC (exponential vs. Weibull) with four data summaries (no summary, mean of the data, inverse of the individual data, inverse of the mean, and mean of the inverted). In general, we found that less summarization/transformation (i.e., working with the untransformed, individual level data) with the appropriate matching distribution uniformly performs best in all circumstances by multiple metrics. Conversely, inverting the data or inverting and then taking an average performs poorly.

However, the adequacy of the other approaches is more subtle. From the simulation results of the experiment using data generated from an exponential distribution with a wide temperature interval, modeling with individual-level data was able to capture the true values of *T*_opt_, *L*_Max_, *T*_Upper_, and *T*_Lower_ in the HDI intervals or boxplot whiskers of the chosen dataset, as well as having one of the smallest uncertainties across all datasets. It also achieved the coverage proportions closest to 0.95 for all parameters. Modeling using the mean data was able to capture most true values in the chosen dataset, but had higher uncertainty across all datasets, approximately 1.8 times as large as the individual data when estimating *T*_Upper_ and *T*_Lower_. Modeling with the inverse of the mean data captured all true values, performing adequately but again with greater uncertainty than individual-level data.

Results were similar when modeling data generated from a Weibull distribution and a wide temperature interval. As for the exponentially distributed data, individual-level data with a Weibull model had the smallest uncertainty of the estimated values of *T*_opt_ across all models, and the second smallest uncertainty in all other values. This method also had the best coverage proportions, which were near (or equal to) 0.95 for all parameters. Using the mean data performed similarly, with the second-best coverage proportion for parameters *a* and *c*, as well as all true values (*T*_opt_, *L*_Max_, *T*_Upper_, and *T*_Lower_) within the 75% HDI bounds or whiskers of the boxplots for the visualized dataset. Using the IoM data with a truncated normal model managed to capture most true values as well, but again, with higher uncertainty. Both the individual inverse data and MoI data methods were unable to capture the tails of the curve and had extremely poor coverage proportions.

Interestingly, using the individual data but assuming the wrong model structure (i.e. assuming that the data are exponential when they are Weibull) produced worse results than using the mean of the lifetime and modeling with a truncated normal. Assuming the exponential distribution when the data were Weibull produced reasonable parameter estimates, but substantially larger uncertainty around them, and thus around the TPC (Figure 14). When estimating *T*_opt_, *L*_Max_, *T*_Upper_, and *T*_Lower_, the HDI bounds (e.g. Figure 10) and the length of the whiskers of the boxplots (e.g. Figure 11) were approximately 3 times that of using a Weibull model. In Figure 14, we overlay the curves made from evaluating the joint posterior samples of the models fit to a single dataset of individual data using both a Weibull and an exponential distribution (previously shown in Figures 9a and 9b). While both models do a good job of curve estimation, the Weibull model produces smaller bounds and less uncertainty. This suggests that even when the true data-generating process is not correctly specified, reasonable estimates can still be achieved. However, using the correct model reduces uncertainty and leads to more precise results.

**Figure 14:**
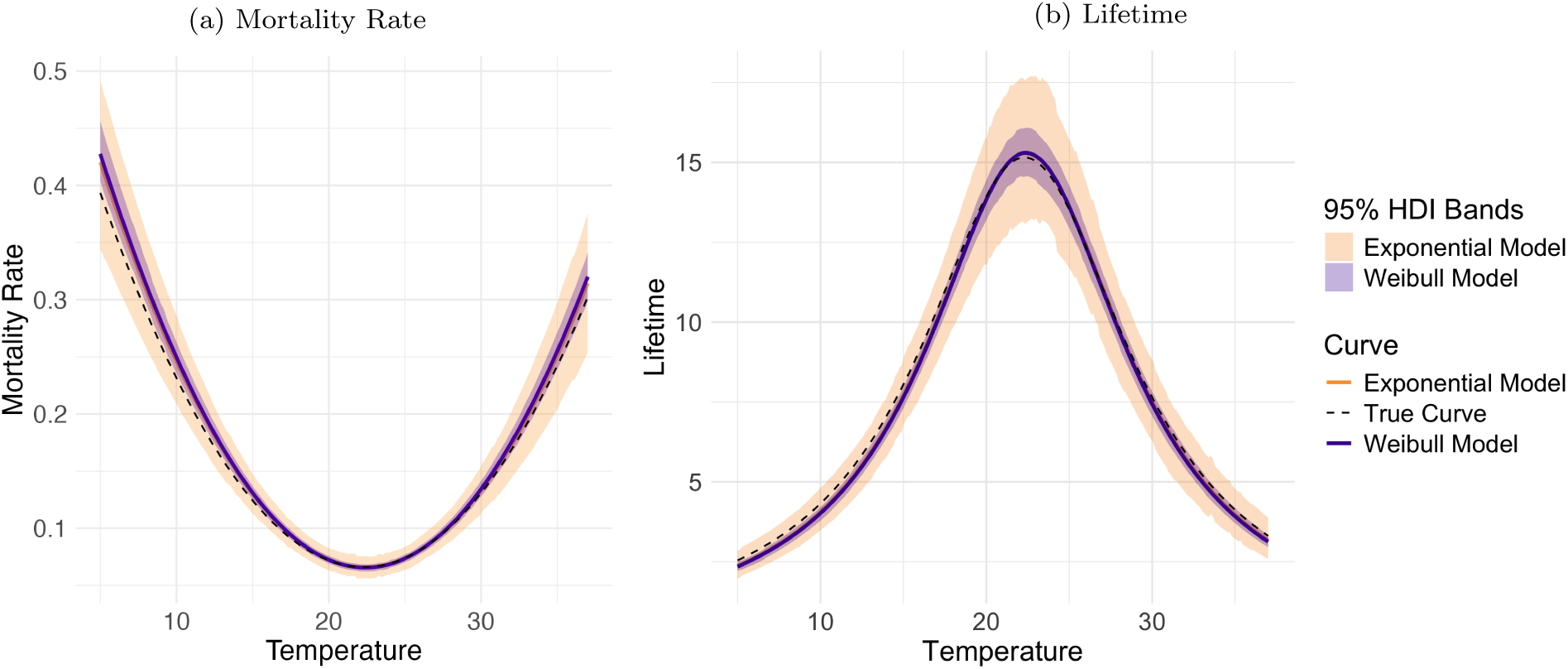
Comparison of the median mortality rate (14a) and lifetime curves (14b) from an exponential model and a Weibull model when using individual data generated from a Weibull distribution. Curves were made by evaluating samples of the posterior function across a temperature range from 5^◦^C to 37^◦^C. The median response at each temperature is shown as the solid lines, while the true curve is a dashed line. The 95% HDI bounds of the response at each temperature were also plotted as the orange (exponential) and purple (Weibull) ribbons.

Together, these results indicate approaches to obtain good estimates of the TPCs. First, raw, individual-level data are preferred when possible, and thus the raw data should also be published/provided when feasible. Then, before modeling, data exploration across temperatures should be used to ensure that a reasonable data distribution is chosen. For example, an exponential should only be used if the data clearly indicate an exponential pattern. If only summary data (usually averages) are available, modeling the mean data can recover parameter values and shape of the TPC. However, this results in greater uncertainty, particularly in the tails. This suggests that while mean data can still be used, we may be losing valuable information and that researchers should be cautious of their estimates if interested in something other than the peak. Further, the models will not be able to make predictions about the distribution of observed trait in that case – only the TPC itself.

If the goal is to model the mortality rate directly rather than the raw data, it is important to note that inverting individual observations before averaging them introduces bias. Because 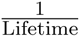 is a convex function, averaging per-individual rates 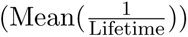 produces an upward bias relative to 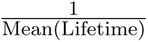 (Ruel and Ayres, 1999). In other words, inverting observations before averaging tends to overestimate the underlying rate, as seen in our simulation experiments (Figures 9f and 5e). Thus, if modeling the rate, one should take the mean-level data and then invert it (IoM data) and model these transformed observations with a truncated normal distribution. However, we suggest that the mortality rate framework should probably be avoided unless one is either fairly sure the data are exponentially distributed or if one is strictly interested in comparing results to earlier mortality rate calculations.

We also found that, as expected, the design of the experiment, specifically the range of temperatures considered, impacts the inference of TPC parameters. Based on the results of our simulation experiments using both wide and narrow temperature intervals, when a researcher is trying to model the rate directly, a wide temperature interval is imperative. Methods based on inverted lifetime data had more difficulty covering peak, tail, and true parameter values and typically resulted in larger uncertainty in estimates, as well as exhibiting higher RMSEs, indicating that wider temperature ranges improve overall model performance. When using individual-level or mean data, the experimental design had a smaller influence on the results. Although a wider interval is always preferred to decrease uncertainty, particularly in the tails, models fit to data generated under a narrow temperature interval were still able to capture the same true values as their wide-interval counterparts when lifetime was modeled directly. If a narrow interval is the only data available, it can serve as a practical alternative, but researchers should be aware that this approach results in increased uncertainty in estimates. This is in part because large uncertainties can limit inter-pretability; even when estimates contain the true value, the resulting intervals may be too large to support meaningful conclusions. For example, while the model using individual inverse data under a narrow temperature interval captured the true value of *T*_Upper_ within the boxplot whiskers, the estimated range spanned more than 150^◦^C, limiting the practical conclusions that can be drawn.

There are a few drawbacks and limitations to our simulation experiment and analysis. In our simulation, we used the exact simulated lifetimes, only truncating the data so that no lifetime would be below one day, corresponding to a maximum mortality rate of one. In physical experiments, researchers often record mortality once each day rather than constantly monitoring individuals. This results in lifetimes that are rounded up to the next whole day, even if the individual died much earlier. This systematic rounding will bias results towards longer lifetimes. This should not impact inference of the temperature at the optimum, but is likely to impact the level at the optimum. Further, other factors will influence the specific results, including the relative width of uncertainty bands between methods. For example, the calculated differences between methods and uncertainties in estimates will depend on things such as the trait being modeled and the sample size, and the underlying TPC shape.

Although substantial work has quantified the relationship between temperature and physiological traits or attempted to find the “best” fitting TPC, few studies have explored the effects of the data-generating mechanism itself on the analysis. As we have seen here, the assumed trait distribution (here exponential or Weibull for lifetimes) will affect the rest of the analysis, including the parameter estimates and uncertainty in the thermal performance curve. The current literature often assumes exponential waiting times for estimated rates, such as for dynamical systems, yet studies have shown that the mortality rate decelerates with age (Carey et al., 1992; Curtsinger et al., 1992; Wilson, 1994). Unlike the exponential distribution, which assumes a constant rate, the Weibull distribution allows the mortality rate to change with increasing age, suggesting that it may be a better fit for lifetime data in arthropods (Ricklefs and Scheuerlein, 2002). Because lifetime influences many physiological traits that are paramount in modeling populations, modeling it poorly could negatively effect related analyses.

Our findings indicate a set of potentially best practices for researchers to follow when fitting TPCs. Researchers should prioritize obtaining individual-level data collected across as many temperature levels as possible over the widest feasible interval, as this approach yields the most accurate models with the least uncertainty. When individual-level data are available, the distribution of the data at each temperature will inform which model should be used. If researchers wish to model the rate from available trait data, our simulations show that taking the mean of the data first and then inverting it to obtain the rate (i.e. using the IoM data) provides a much better fit and less uncertainty than using individual-level inverted data. While demonstrated for lifetime data, these guidelines should be applicable to other thermal traits of arthropods, especially those that are based on waiting times, such as development time. This should enable a better use of valuable and difficult to collect data on thermal traits, and support a variety of modeling approaches in the future.

## Acknowledgements

POZ and LRJ were partially funded by NSF DEB #2017463 and NSF DBI #2016264.

## Notes

### Competing Interest Statement

The authors have declared no competing interest.

