## Supplemental Materials for "Best Practices for Modeling Arthropod Lifetimes in a Bayesian Framework"

### 1 Transformation of Variables

This section details the transformation of variables used when summarizing data to use means, inverting it to obtain a "rate", taking the mean and then inverting, and inverting and then taking a mean. It is important to note that in some cases, we rely on the Central Limit Theorem (CLT) to obtain distributions. There are several things to be wary of when using the CLT to determine the distributions of our data. The first issue is the sample size. While this approximation becomes more accurate as sample size increases, its reliability decreases when there is a small number of observations in each mean value. Unless the individual dataset is large enough to have each mean composed of a (generally accepted) size of at least 30 individuals, this approximation may not be valid. Secondly, the mean must be sufficiently far from zero so that the normal approximation will not assign probability to negative lifetimes, which are biologically unrealistic.

#### 1.1 Exponential Distribution

This subsection includes the derivations for the transformation of variables for the different variations of data generated from an exponential distribution, including the transformation to the mean data, the inverse data, the inverse of the mean data, and the mean of the inverse data. For reference, if $x$ is a exponentially distributed random variable with rate parameter $\mu$, the pdf of $x$ is $f(x) = \mu e^{-\mu x}$. Table S1 shows a summary of each transformation's pdf, mean, and variance.

| Transformation | pdf | Mean | Variance |
|---|---|---|---|
| Mean | $f_Y(y) = \frac{(\mu n)^n}{\Gamma(n)} y^{n-1} e^{-\mu n y}$ | $\frac{1}{\mu}$ | $\frac{1}{\mu^2 n}$ |
| Inverse | $f(y) = \frac{\mu}{y^2} e^{-\frac{\mu}{y}}$ | NCFS | NCFS |
| Inverse of Mean | $f_Y(y) = \frac{(\mu n)^n}{\Gamma(n)} \left(\frac{1}{y}\right)^{n+1} e^{-\frac{\mu n}{y}}$ | NCFS | NCFS |
| Mean of Inverse | $f(y) = \frac{1}{\sqrt{2\pi\tilde{\sigma}^2}} e^{-\frac{1}{2\tilde{\sigma}^2}\left(y-\frac{1}{\mu}\right)^2}$ | NCFS | NCFS |

Table S1: The pdf, mean, and variance for each transformation of the exponential data. NCFS stands for no closed form solution. $\tilde{\sigma}^2$ in the pdf represents the calculated variance for that transformation, and can be found in the corresponding section.

##### 1.1.1 Mean Transformation

Below is the derivation showing the transformation of variables used to get the distribution of a mean transformation, $\bar{x}$, if $x$ is an exponential random variable. The sum of $n$ exponential variables with rate parameter $\mu$ is distributed as a Gamma distribution with parameters $n$ and $\mu$. First, we will prove this showing the MGF of both an exponential distribution and a Gamma distribution. Then, we will scale this by $\frac{1}{n}$ to get the distribution for the mean of exponential random variables.

MGF of exponential:

$$M_{exp}(t) = \int_0^\infty e^{tx} \mu e^{-\mu x} dx$$

$$M_{exp}(t) = \mu \int_0^\infty e^{x(t-\mu)} dx$$

$$M_{exp}(t) = \frac{\mu}{t-\mu} e^{x(t-\mu)} \Big|_0^\infty$$

$$M_{exp}(t) = \lim_{x \to \infty} \left( \frac{\mu}{t-\mu} e^{x(t-\mu)} - \frac{\mu}{t-\mu} \right)$$

$$M_{exp}(t) = \frac{\mu}{t-\mu} \lim_{x \to \infty} (e^{x(t-\mu)} - 1)$$

With the stipulation that $t < \mu$:

$$M_{exp}(t) = \frac{\mu}{t-\mu}(0-1)$$

$$M_{exp}(t) = \frac{\mu}{\mu - t}$$

MGF of Gamma:

$$M_{gam}(t) = \frac{b^a}{\Gamma(a)} \int_0^\infty x^{a-1} e^{-(a-t)x} dx$$

Let $u = x(b-t)$:

$$M_{gam}(t) = \frac{b^a}{\Gamma(a)} \int_0^\infty \left( \frac{u}{b-t} \right)^{a-1} e^{-u} \frac{dx}{b-t}$$

$$M_{gam}(t) = \frac{b^a}{\Gamma(a)(b-t)^a} \int_0^\infty u^{a-1} e^{-u} du$$

$$M_{gam}(t) = \frac{b^a \Gamma(a)}{\Gamma(a)(b-t)^a}$$

$$M_{gam}(t) = \frac{b^a}{(b-t)^a}$$

$$M_{gam}(t) = \left(\frac{b}{b-t}\right)^a$$

Using the property that for $W = W_1 + W_2 + ... + W_n$, the MGF is $M_W(t) = M_{W_1}(t) \times M_{W_2}(t) \times ... \times M_{W_n}(t)$, we have the MGF of the sum of $n$ exponential variables to be $\left(\frac{\mu}{\mu-t}\right)^n$, which is the MGF of a Gamma distribution with parameters $n$ and $\mu$. Scaling this distribution by $\frac{1}{n}$ affects the rate parameter, which becomes $\mu n$. Thus, the mean of $n$ exponential random variables is a Gamma distribution with parameters $n$ and $\mu n$,

$$f_Y(y) = \frac{(\mu n)^n}{\Gamma(n)} y^{n-1} e^{-\mu n y}.$$

The properties of a Gamma distribution tell us that the expected value is $\frac{1}{\mu}$, and the variance is $\frac{1}{\mu^2 n}$.

##### 1.1.2 Inverse Transformation

Below is the derivation showing the transformation of variables used to get the distribution of an inverse transformation, $\frac{1}{x}$, if $x$ is an exponential random variable. Let $y = \frac{1}{x}$, so that $x = \frac{1}{y}$ and $\left|\frac{\partial x}{\partial y}\right| = \frac{1}{y^2}$. Following the form $f_Y(y) = f_X(g^{-1}(y)) \left|\frac{\partial}{\partial y} g^{-1}(y)\right|$:

$$f_X(x) = \mu e^{-\mu x}$$

$$f_Y(y) = f_X\left(\frac{1}{y}\right)\left|\frac{\partial x}{\partial y}\right|$$

$$f\left(\frac{1}{x}\right) = f(y) = \frac{\mu}{y^2} e^{-\frac{\mu}{y}}$$

The pdf of an inverse transformation on an exponential random variable is $f(y) = \frac{\mu}{y^2} e^{-\frac{\mu}{y}}$. The calculation to find the expected value is as follows:

$$E(y) = \int_0^\infty \frac{\mu}{y} e^{-\frac{\mu}{y}} dy$$

Substituting $u = \frac{\mu}{y}$ and $du = -\frac{\mu}{y^2} dy$ gives us:

$$E(y) = \mu \int_\infty^0 -\frac{e^{-u}}{u} du$$

$\int_\infty^0 -\frac{e^{-u}}{u} du$ is an exponential integral and can be rewritten as $E_1(u)\Big|_0^\infty$. Substituting this in and reverting to $y$ gives:

$$E(y) = \mu E_1(u)\Big|_0^\infty$$

$$E(y) = \mu E_1\left(\frac{\mu}{y}\right)\Big|_0^\infty$$

This does not have a closed form solution, so we will need to rely on the simulation for results.

Using $V(y) = E(y^2) - (E(y))^2$, the calculation to find the variance is as follows:

$$E(y^2) = \int_0^\infty \mu e^{-\frac{\mu}{y}} dy$$

$$E(y^2) = \mu \int_0^\infty e^{-\frac{\mu}{y}} dy$$

Let $u = e^{-\frac{\mu}{y}}$, $du = \frac{\mu e^{-\frac{\mu}{y}}}{y^2}$, $dv = 1$, and $v = y$. Integrating by parts, $uv - \int duv$ gives us:

$$\mu\left(ye^{-\frac{\mu}{y}}\right) - \int_0^\infty \frac{\mu e^{-\frac{\mu}{y}}}{y} dy$$

Substituting $z = \frac{\mu}{y}$ and $dz = -\frac{\mu}{y^2} dy$ gives us:

$$\mu\left(ye^{-\frac{\mu}{y}} - \mu \int_0^\infty -\frac{e^{-z}}{z} dz\right)$$

$\int_0^\infty -\frac{e^{-z}}{z}dz$ can be written as $E_1(z)\Big|_0^\infty$, which represents the exponential integral.

$$E\left(y^2\right) = \mu y e^{-\frac{\mu}{y}} - \mu^2 E_1\left(\frac{\mu}{y}\right)\Big|_0^\infty$$

$$V(y) = \mu y e^{-\frac{\mu}{y}} - \mu^2 E_1\left(\frac{\mu}{y}\right)\Big|_0^\infty - \left(\mu E_1\left(\frac{\mu}{y}\right)\Big|_0^\infty\right)^2$$

This does not have a closed form solution, so we will need to rely on the simulation for results.

###### 1.1.3 Inverse of Mean Transformation

Below is the derivation showing the transformation of variables used to get the distribution of an inverse of the mean transformation, $\frac{1}{\bar{x}}$, if $x$ is an exponential random variable. From calculating the distribution of a mean variable, we know that $\bar{x}$ is following a Gamma distribution with parameters $n$ and $\mu n$. Let $y = \frac{1}{z}$, where $z$ represents the mean of exponential random variables, so that $z = \frac{1}{y}$ and $\left|\frac{\partial z}{\partial y}\right| = \frac{1}{y^2}$. Following the form $f_Y(y) = f_Z(g^{-1}(y))\left|\frac{\partial}{\partial y}g^{-1}(y)\right|$:

$$f_Z(z) = \frac{(\mu n)^n}{\Gamma(n)} z^{n-1} e^{-\mu n z}$$

$$f_Y(y) = f_Z\left(\frac{1}{y}\right)\left|\frac{\partial z}{\partial y}\right|$$

$$f(y) = f\left(\frac{1}{z}\right) = \frac{(\mu n)^n}{\Gamma(n)}\left(\frac{1}{y}\right)^{n-1} e^{-\frac{\mu n}{y}}\left(\frac{1}{y^2}\right)$$

$$f\left(\frac{1}{\bar{x}}\right) = f\left(\frac{1}{z}\right) = f(y) = \frac{(\mu n)^n}{\Gamma(n)}\left(\frac{1}{y}\right)^{n+1} e^{-\frac{\mu n}{y}}$$

The pdf of a mean transformation, then an inverse transformation on an exponential random variable is $f(y) = \frac{(\mu n)^n}{\Gamma(n)}\left(\frac{1}{y}\right)^{n+1} e^{-\frac{\mu n}{y}}$.

The calculation to find the expected value is as follows:

$$E(y) = \int_0^\infty \frac{(\mu n)^n}{\Gamma(n)} \left(\frac{1}{y}\right)^n e^{-\frac{\mu n}{y}} dy$$

$$E(y) = \frac{(\mu n)^n}{\Gamma(n)} \int_0^\infty \frac{e^{-\frac{\mu n}{y}}}{y^n} dy$$

Substituting $u = n^{n-1}\mu^{n-1}y^{1-n}$ and $du = \frac{(1-n)n^{n-1}\mu^{n-1}}{y^n}$ gives us:

$$E(y) = \frac{(\mu n)^n}{\Gamma(n)} \left(-\frac{n^{1-n}\mu^{1-n}}{n-1} \int_0^\infty e^{\frac{1}{u^{\frac{1}{1-n}}}} du\right)$$

$\int_0^\infty e^{\frac{1}{u^{\frac{1}{1-n}}}} du$ is an incomplete gamma function and can be rewritten as $-(n-1)(-1)^{n-1}\Gamma\left(n-1, \frac{1}{u^{\frac{1}{1-n}}}\right)\Big|_0^\infty$. After plugging this back in and simplifying, the expected value comes to

$$E(y) = \frac{(n\mu)\Gamma\left(n-1, -\frac{n\mu}{y}\right)}{\Gamma(n)(-1)^n}\Bigg|_0^\infty,$$

which does not have a closed form solution, and we will need to rely on the simulation. Using $V(y) = E(y^2) - (E(y))^2$, the calculation to find the variance is as follows:

$$E(y^2) = \int_0^\infty \frac{(\mu n)^n}{\Gamma(n)} y^{1-n} e^{-\frac{\mu n}{y}} dy$$

$$E(y^2) = \frac{(\mu n)^n}{\Gamma(n)} \int_0^\infty y^{1-n} e^{-\frac{\mu n}{y}} dy$$

Substituting $u = n^{n-2}\mu^{n-2}y^{2-n}$ and $du = (2-n)n^{n-2}\mu^{n-2}y^{1-n}$ gives us:

$$E(y^2) = \frac{(\mu n)^n}{\Gamma(n)} \left(-\frac{n^{2-n}\mu^{2-n}}{n-2} \int_0^\infty e^{\frac{1}{u^{\frac{1}{2-n}}}} du\right)$$

$\int_0^\infty e^{\frac{1}{u^{\frac{1}{2-n}}}} du$ is an incomplete gamma function and can be rewritten as

$$(n-2)(-1)^{2-n}\Gamma\left(n-2, \frac{1}{u^{\frac{1}{2-n}}}\right)\Big|_0^\infty$$

. After plugging this back in and simplifying, the integral comes to

$$E(y^2) = \frac{(n\mu)^2\Gamma\left(n-2, -\frac{n\mu}{y}\right)}{\Gamma(n)(-1)^n}\Big|_0^\infty,$$

which does not have a closed form solution, and we will need to rely on the simulation. The variance then comes to $\frac{(n\mu)^2\Gamma\left(n-2,-\frac{n\mu}{y}\right)}{\Gamma(n)(-1)^n}\Big|_0^\infty - \left(\frac{(n\mu)\Gamma\left(n-1,-\frac{n\mu}{y}\right)}{\Gamma(n)(-1)^n}\Big|_0^\infty\right)^2$.

##### 1.1.4 Mean of Inverse Transformation

When attempting to transform the random variable to the mean of the inverse data, it would require a mean transformation to the previously derived inverse distribution, $f(y) = \frac{\mu}{y^2}e^{-\frac{\mu}{y}}$. Let $z = \frac{\sum_{i=1}^n y}{n}$. We would need the Jacobian and the pdf of the transformation. While this is not available in closed form solution, the Central Limit Theorem tells us that it will be approximately normal with mean of the inverse distribution, and variance of the inverse distribution divided by the sample size. As calculated previously, the mean is $\tilde{\mu} = \mu E_1\left(\frac{\mu}{y}\right)\Big|_0^\infty$, and the variance will be $\tilde{\sigma}^2 = \mu y e^{-\frac{\mu}{y}} - \mu^2 E_1\left(\frac{\mu}{y}\right)\Big|_0^\infty - \left(\mu E_1(\frac{\mu}{y}\Big|_0^\infty\right)^2$. Since our $n$ is relatively small, it is not a perfect approximation, but as the CLT states, it should be approaching this normal distribution as $n$ approaches infinity. Therefore, the pdf is $f(y) = \frac{1}{\sqrt{2\pi\tilde{\sigma}^2}}e^{-\frac{1}{2\tilde{\sigma}^2}(y-\tilde{\mu})^2}$, where $\tilde{\mu}$ and $\tilde{\sigma}^2$ are given above.

#### 1.2 Weibull Distribution

This subsection includes the derivations for the transformation of variables for the different variations of data generated from a Weibull distribution, including transformations to the mean data, the inverse data, the inverse of the mean data, and the mean of the inverse data. For reference, if $x$ is distributed as a Weibull random variable with shape parameter $k$ and scale parameter $\alpha$, the pdf of $x$ is $f(x) = \left(\frac{k}{\alpha}\right)\left(\frac{x}{\alpha}\right)^{k-1} e^{-\left(\frac{x}{\alpha}\right)^k}$. Table S2 shows a summary of each transformation's pdf, mean, and variance.

| Transformation | pdf | Mean | Variance |
|---|---|---|---|
| Mean | $f(y) = \frac{1}{\sqrt{2\pi\tilde{\sigma}^2}} e^{-\frac{1}{2\tilde{\sigma}^2}(y-\tilde{\mu})^2}$ | $\alpha\Gamma\left(1+\frac{1}{k}\right)$ | $\frac{\alpha^2\left(\Gamma\left(1+\frac{2}{k}\right)-\Gamma^2\left(1+\frac{1}{k}\right)\right)}{n}$ |
| Inverse | $f(y) = \left(\frac{k}{\alpha}\right)\left(\frac{1}{\alpha y}\right)^{k-1}\left(\frac{1}{y^2}\right) e^{-\left(\frac{1}{\alpha y}\right)^k}$ | NCFS | NCFS |
| Inverse of Mean | $f(y) = \frac{1}{\sqrt{2\pi\tilde{\sigma}^2}} e^{-\frac{\left(\frac{1}{y}-\tilde{\mu}\right)^2}{2\tilde{\sigma}^2}} \frac{1}{y^2}$ | NCFS | NCFS |
| Mean of Inverse | $f(y) = \frac{1}{\sqrt{2\pi\tilde{\sigma}^2}} e^{-\frac{1}{2\tilde{\sigma}^2}(y-\tilde{\mu})^2}$ | NCFS | NCFS |

Table S2: The pdf, mean, and variance for each transformation of the Weibull data. NCFS stands for no closed form solution. $\tilde{\mu}$ and $\tilde{\sigma}^2$ in each pdf represent the calculated mean and variance for that transformation, and can be found in the corresponding section.

##### 1.2.1 Mean Transformation

Below is the derivation showing the transformation of variables used to get the distribution of a mean transformation, $\bar{x}$, if $x$ is a Weibull random variable. The CLT states that the mean of random variables that follow a Weibull distribution will be approximately normal, with mean and variance from the Weibull distribution. Using the moments of the Weibull distribution and the fact that $E(X^n) = \alpha^n\Gamma\left(1+\frac{n}{k}\right)$ from McCool (2012), we can say that it will follow an approximate normal distribution with mean $\tilde{\mu} = \alpha\Gamma\left(1+\frac{1}{k}\right)$ and variance $\tilde{\sigma}^2 = \frac{\alpha^2\left(\Gamma\left(1+\frac{2}{k}\right)-\Gamma^2\left(1+\frac{1}{k}\right)\right)}{n}$, where $k$ is the shape and $\alpha$ is the scale. Therefore, the pdf is $f(y) = \frac{1}{\sqrt{2\pi\tilde{\sigma}^2}} e^{-\frac{1}{2\tilde{\sigma}^2}(y-\tilde{\mu})^2}$, where $\tilde{\mu}$ and $\tilde{\sigma}^2$ are given above.

##### 1.2.2 Inverse Transformation

Below is the derivation showing the transformation of variables used to get the distribution of an inverse transformation, $\frac{1}{x}$, if $x$ is a Weibull random variable. Let $y = \frac{1}{x}$, so that $x = \frac{1}{y}$ and $\left|\frac{\partial x}{\partial y}\right| = \frac{1}{y^2}$. Following the form $f_Y(y) = f_X(g^{-1}(y))\left|\frac{\partial}{\partial y} g^{-1}(y)\right|$:

$$f(x) = \left(\frac{k}{\alpha}\right)\left(\frac{x}{\alpha}\right)^{k-1} e^{-\left(\frac{x}{\alpha}\right)^k}$$

$$f_Y(y) = f_X\left(\frac{1}{y}\right)\left|\frac{\partial x}{\partial y}\right|$$

$$f\left(\frac{1}{x}\right) = f(y) = \left(\frac{k}{\alpha}\right)\left(\frac{1}{\alpha y}\right)^{k-1}\left(\frac{1}{y^2}\right)e^{-\left(\frac{1}{\alpha y}\right)^k}$$

The pdf of an inverse transformation on a Weibull random variable is $f(y) = \left(\frac{k}{\alpha}\right)\left(\frac{1}{\alpha y}\right)^{k-1}\left(\frac{1}{y^2}\right)e^{-\left(\frac{1}{\alpha y}\right)^k}$.

The calculation to find the expected value is as follows:

$$E(y) = \left(\frac{k}{\alpha}\right)\left(\frac{1}{\alpha}\right)^{k-1}\int_0^\infty y^{-k}e^{-\left(\frac{1}{\alpha y}\right)^k}dy$$

$$E(y) = k\left(\frac{1}{\alpha}\right)^k\int_0^\infty \frac{e^{-\left(\frac{1}{\alpha y}\right)^k}}{y^k}dy$$

Substituting $u = \alpha^{1-k}y^{1-k}$ and $du = \frac{(1-k)\alpha^{1-k}}{y^k}dy$ gives us:

$$E(y) = k\alpha^{-k}\left(-\frac{\alpha^{k-1}}{k-1}\right)\int_0^\infty e^{-\frac{1}{u^{\frac{k}{1-k}}}}du$$

$\int_0^\infty e^{-\frac{1}{u^{\frac{k}{1-k}}}}du$ is an incomplete gamma function and can be rewritten as $\frac{(1-k)\Gamma\left(-\frac{1-k}{k}, \frac{1}{u^{\frac{k}{1-k}}}\right)}{k}\Big|_0^\infty$.

After plugging this back in and simplifying, the integral comes to

$$E(y) = k\alpha^{-k}\left(\frac{(1-k)\alpha^{k-1}\Gamma\left(-\frac{1-k}{k}, \left(\frac{1}{\alpha y}\right)^k\right)}{k(k-1)}\right)\Bigg|_0^\infty$$

$$E(y) = \frac{\Gamma\left(\frac{k-1}{k}, \left(\frac{1}{\alpha y}\right)^k\right)}{\alpha}\Bigg|_0^\infty$$

which does not have a closed form solution, and we will need to rely on the simulation.

Using $V(y) = E(y^2) - (E(y))^2$, the calculation to find the variance is as follows:

$$E(y^2) = \left(\frac{k}{\alpha}\right)\left(\frac{1}{\alpha}\right)^{k-1} \int_0^\infty y^{1-k} e^{-\left(\frac{1}{\alpha y}\right)^k} dy$$

$$E(y^2) = k\alpha^{-k} \int_0^\infty e^{-\left(\frac{1}{\alpha y}\right)^k} y^{1-k} dy$$

Substituting $u = \alpha^{2-k}y^{2-k}$ and $du = (2-k)\alpha^{2-k}y^{1-k}dy$ gives us:

$$E(y^2) = -\frac{k}{\alpha^2(k-2)} \int_0^\infty e^{-\frac{1}{u^{\frac{k}{2-k}}}} du$$

$\int_0^\infty e^{-\frac{1}{u^{\frac{k}{2-k}}}} du$ is an incomplete gamma function and can be rewritten as $\left.\frac{(2-k)\Gamma\left(-\frac{2-k}{k}, \frac{1}{u^{\frac{k}{2-k}}}\right)}{k}\right|_0^\infty$. After plugging this back in and simplifying, the integral comes to

$$E(y^2) = \left.\frac{(2-k)\Gamma\left(-\frac{2-k}{k}, \left(\frac{1}{\alpha y}\right)^k\right)}{\alpha^2(k-2)}\right|_0^\infty$$

$$E(y^2) = \left.\frac{\Gamma\left(\frac{k-2}{k}, \left(\frac{1}{\alpha y}\right)^k\right)}{\alpha^2}\right|_0^\infty$$

which does not have a closed form solution, and we will need to rely on the simulation. The variance then comes to $\left.\frac{\Gamma\left(\frac{k-2}{k},\left(\frac{1}{\alpha y}\right)^k\right)}{\alpha^2}\right|_0^\infty - \left(\left.\frac{\Gamma\left(\frac{k-1}{k},\left(\frac{1}{\alpha y}\right)^k\right)}{\alpha}\right|_0^\infty\right)^2$.

##### 1.2.3 Inverse of Mean Transformation

Below is the derivation showing the transformation of variables used to get the distribution of an inverse of the mean transformation, $\frac{1}{\bar{x}}$, if $x$ is an exponential random variable. From calculating the distribution of a mean variable, we know that $\bar{x}$ is approximately normally distributed with mean $\tilde{\mu} = \alpha\Gamma\left(1 + \frac{1}{k}\right)$ and variance $\tilde{\sigma}^2 = \frac{\alpha^2\left(\Gamma\left(1+\frac{2}{k}\right)-\Gamma^2\left(1+\frac{1}{k}\right)\right)}{n}$, where $k$ is the shape and $\alpha$ is the scale. Let $y = \frac{1}{z}$, where $z$ represents the mean of Weibull random variables, so that $z = \frac{1}{y}$ and $\left|\frac{\partial z}{\partial y}\right| = \frac{1}{y^2}$. Following the form $f_Y(y) = f_Z(g^{-1}(y))\left|\frac{\partial}{\partial y}g^{-1}(y)\right|$:

$$f_Z(z) = \frac{1}{\sqrt{2\pi\sigma^2}}e^{-\frac{(z-\mu)^2}{2\sigma^2}}$$

$$f_Y(y) = f_Z\left(\frac{1}{y}\right)\left|\frac{\partial z}{\partial y}\right|$$

$$f\left(\frac{1}{z}\right) = f(y) = \frac{1}{\sqrt{2\pi\tilde{\sigma}^2}}e^{-\frac{\left(\frac{1}{y}-\tilde{\mu}\right)^2}{2\tilde{\sigma}^2}}\frac{1}{y^2}$$

where $\tilde{\mu}$ and $\tilde{\sigma}^2$ are defined above. Since our distribution for the mean of variables following a Weibull distribution was approximate due to using the CLT, this distribution will also be approximate.

The pdf of a mean transformation then an inverse transformation on a Weibull random variable is approximately $f(y) = \frac{1}{\sqrt{2\pi\tilde{\sigma}^2}}e^{-\frac{\left(\frac{1}{y}-\tilde{\mu}\right)^2}{2\tilde{\sigma}^2}}\frac{1}{y^2}$, with $\tilde{\mu}$ and $\tilde{\sigma}^2$ defined above. The integrals to find the expected value and the variance have no closed form solutions, so we have to rely on the simulation experiment for results.

##### 1.2.4 Mean of Inverse Transformation

When attempting to transform the random variable to the mean of the inverse, it would require a mean transformation to the previously derived inverse distribution, $f(y) = \left(\frac{k}{\alpha}\right)\left(\frac{1}{\alpha y}\right)^{k-1}\left(\frac{1}{y^2}\right)e^{-\left(\frac{1}{\alpha y}\right)^k}$. Let $z = \frac{\sum_{i=1}^{n} y}{n}$. We would need the Jacobian and the pdf of the transformation. While this is not available in closed form solution, the Central Limit Theorem (CLT) tells us that it will be approximately normal with mean of the inverse distribution, and variance of the inverse distribution divided by the sample size. As calculated previously, the mean and variance are not in closed form solution. The mean is $\tilde{\mu} = \frac{\Gamma(\frac{k-1}{k}, \left(\frac{1}{\alpha y}\right)^k)}{\alpha}\Big|_0^\infty$ and the variance will be $\tilde{\sigma}^2 = \frac{\Gamma(\frac{k-2}{k}, \left(\frac{1}{\alpha y}\right)^k)}{\alpha^2}\Big|_0^\infty - \left(\frac{\Gamma(\frac{k-1}{k}, \left(\frac{1}{\alpha y}\right)^k)}{\alpha}\Big|_0^\infty\right)^2$, scaled by $\frac{1}{n}$. Since our $n$ is relatively small, it is not a perfect approximation, but as the CLT states, it should be approaching this normal distribution as $n$ approaches infinity. Therefore, the pdf of a mean of the inverse transformation is approximately $f(y) = \frac{1}{\sqrt{2\pi\tilde{\sigma}^2}}e^{-\frac{1}{2\tilde{\sigma}^2}(y-\tilde{\mu})^2}$, where $\tilde{\mu}$ and $\tilde{\sigma}^2$ are given above.

#### 2 Priors and Modeling

Although we refer to the model using individual-level data from a Weibull distribution with a Weibull likelihood as a Weibull model, the software used for model fitting does not have an explicit Weibull likelihood, so we use a generalized gamma distribution. However, the generalized gamma can be parameterized to be equivalent to a Weibull distribution, and we use that parameterization here.

When modeling using a truncated normal distribution, in the software used, the parameterization of a normal distribution uses the precision, tau ($\tau$), rather than the standard deviation, $\sigma$. In order to get the correct parameterization, we used the following relationship,

$$\tau = \frac{1}{\sigma^2}. \tag{1}$$

In all modeling scenarios, parameter $a$ was sampled in log space to improve convergence due to its small value. A normal prior with a mean of 0 and a precision of $\frac{1}{10}$ (variance of 10) was used. This was chosen to be a relatively non-informative prior. A normal prior with a mean of 22 and a precision of $\frac{1}{10}$ (variance of 10) was used on parameter $T_{\text{opt}}$. Again, this large variance was chosen so that the prior remained relatively noninformative. The prior was centered around 22 based on the idea that optimal temperatures often fall within a moderate range, frequently centered around approximately 22 (Pawar et al., 2024).

In most cases, an exponential prior with a relatively noninformative rate of 0.5 was placed on parameter $c$. This ensured that $c$ was positive, as well as let the parameter be able to take on many values. However, when modeling both the individual inverse and the mean of the inverse data from an exponential distribution, a gamma prior with a shape of 10 and rate of 1 was placed on parameter $c$. With these two types of data, many of the estimates of the mortality rate were quite higher due to extremely small lifetimes being inverted. A gamma prior with shape 10 and rate 1 pulls $c$ away from zero, but has a large variance which still allows for small values of $c$.

When using a truncated normal model, an exponential prior with a rate parameter of 0.5 was placed on sigma. This ensured values were positive, as well as tending towards smaller values of sigma. With an inverse-gamma prior, which is commonly used as it is the conjugate prior, when sigma is estimated to be near zero, the inference is sensitive to the choices of hyperparameters in the inverse gamma distribution (Gelman, 2006; McElreath, 2018). As we are modeling very small values of mortality rates when using any form of inverted data, we have very small response values, and in turn, small values of sigma. An exponential prior puts more emphasis on smaller values of sigma.

When modeling the individual data from a Weibull distribution with a Weibull model, an exponential prior with a noninformative rate of 0.5 was placed on the shape parameter. This ensured that the shape was positive while being relatively noninformative.

### 3 Data

This section shows examples of each transformation of data using data generated from a narrow temperature interval.

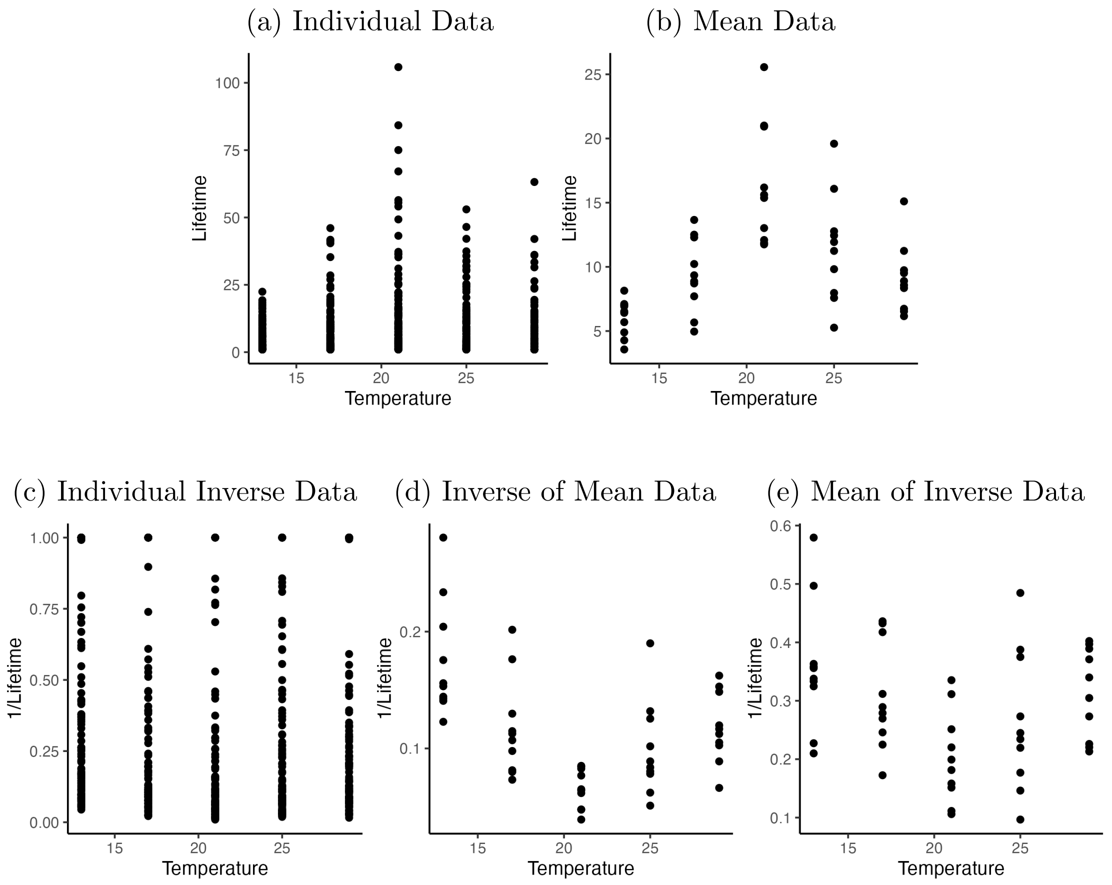


Figure S1: Scatterplots of the $100^{th}$ dataset generated from an exponential distribution under summarization/transformation. S1a Individual data points (raw, untransformed data); S1b mean lifetime (subsets of the raw data are averaged at each temperature); S1c individual inverse data (each raw data point is inverted); S1d inverse of the mean data (i.e, values from S1b are inverted); and S1e is mean of the inverse data (i.e, values from S1c are averaged). Transformations shown are created using the same raw dataset for comparability. The raw lifetimes were truncated at 1 before any transformations.

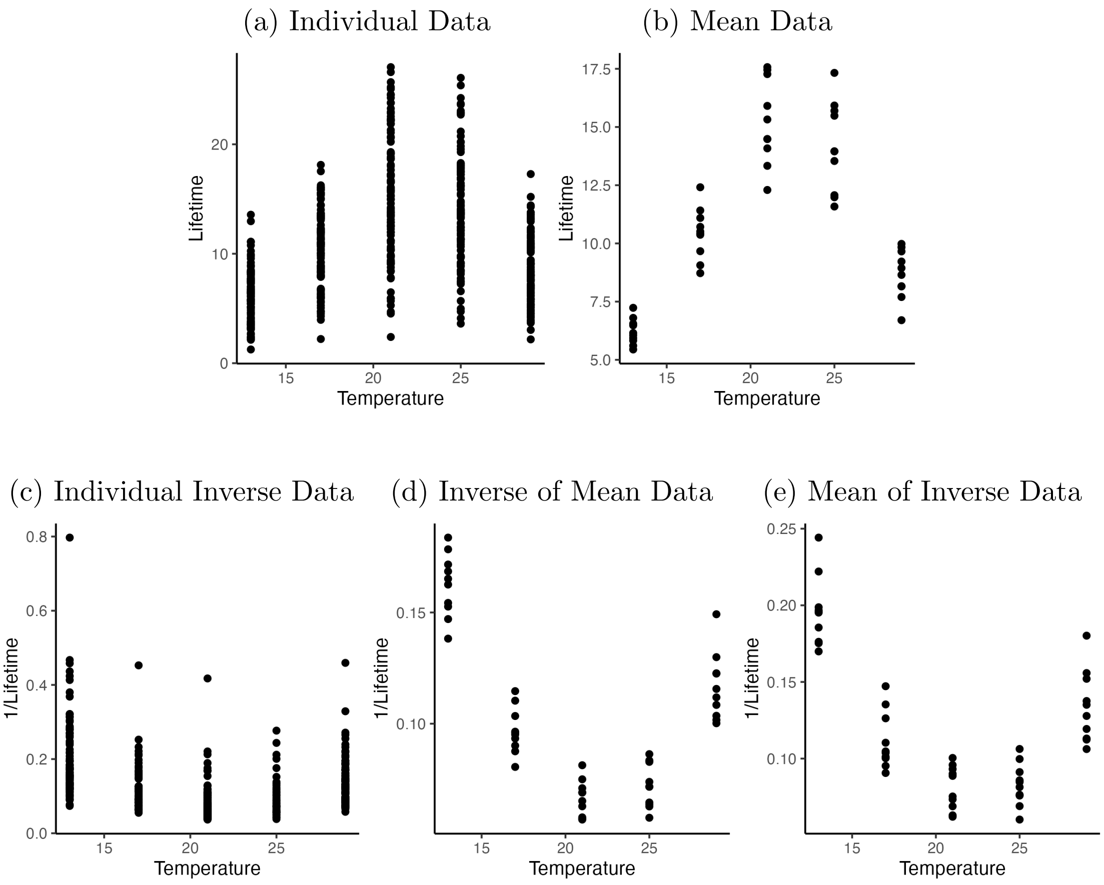


Figure S2: Scatter plots of the $100^{th}$ dataset generated from a Weibull distribution with a narrow temperature interval. S2a Individual data points (raw, untransformed data); S2b mean lifetime (subsets of the raw data are averaged at each temperature); S2c individual inverse data (each raw data point is inverted); S2d inverse of the mean data (i.e, values from S2b are inverted); and S2e is mean of the inverse data (i.e, values from S2c are averaged). Transformations shown are created using the same raw dataset for comparability. The raw lifetimes are truncated at 1 before any transformations.

### 4 Posterior Evaluation

#### 4.1 Parameter Coverage Comparisons

This subsection contains tables of the coverage proportions for each method when using data generated from a narrow temperature interval.

| Parameter Coverage for Narrow Exponential Data | | | |
|---|---|---|---|
| **Data** | **Parameter $a$** | **Parameter $T_{\text{opt}}$** | **Parameter $c$** |
| Individual Data | 0.974 | 0.972 | 0.970 |
| Mean Data | 0.970 | 0.966 | 0.926 |
| Individual Inverse Data | 0.794 | 0.994 | 0.998 |
| Inverse of Mean Data | 0.892 | 0.952 | 0.976 |
| Mean of Inverse Data | 0.876 | 0.950 | 0.000 |

Table S3: Proportion of coverage for parameters $a$, $T_{\text{opt}}$, and $c$. The coverage is calculated as the proportion of chains (5 chains in each model, 100 models for each data variation, 500 chains total) in which the true value of the parameter lay within the 95% HDI bounds of the posterior samples. Values are from models using exponential data generated from a narrow temperature interval.

| Parameter Coverage for Narrow Weibull Data | | | | |
|---|---|---|---|---|
| **Data** | **Parameter $a$** | **Parameter $T_{\text{opt}}$** | **Parameter $c$** | **Parameter $k$** |
| Individual $\text{Data}_{\text{WB}}$ | 0.912 | 0.942 | 0.946 | 0.998 |
| Individual $\text{Data}_{\text{Exp}}$ | 1.000 | 1.000 | 1.000 | NA |
| Mean Data | 0.998 | 0.964 | 0.896 | NA |
| Individual Inverse Data | 0.658 | 0.976 | 0.232 | NA |
| Inverse of Mean Data | 0.914 | 0.922 | 0.980 | NA |
| Mean of Inverse Data | 0.674 | 0.978 | 0.214 | NA |

Table S4: Proportion of coverage for parameters $a$, $T_{\text{opt}}$, $c$, and $k$ (shape). The coverage is calculated as the proportion of chains (5 chains in each model, 100 models for each data variation, 500 chains total) in which the true value of the parameter lay within the 95% HDI bounds of the posterior samples. "Individual $\text{Data}_{\text{WB}}$" represents the individual data with an Weibull model while "Individual $\text{Data}_{\text{Exp}}$" represents the individual data with an exponential model. Values are from models using Weibull data generated from a narrow temperature interval.

#### 4.2 HDI Plots

This subsection shows the HDI plots created when using data from a narrow temperature interval.

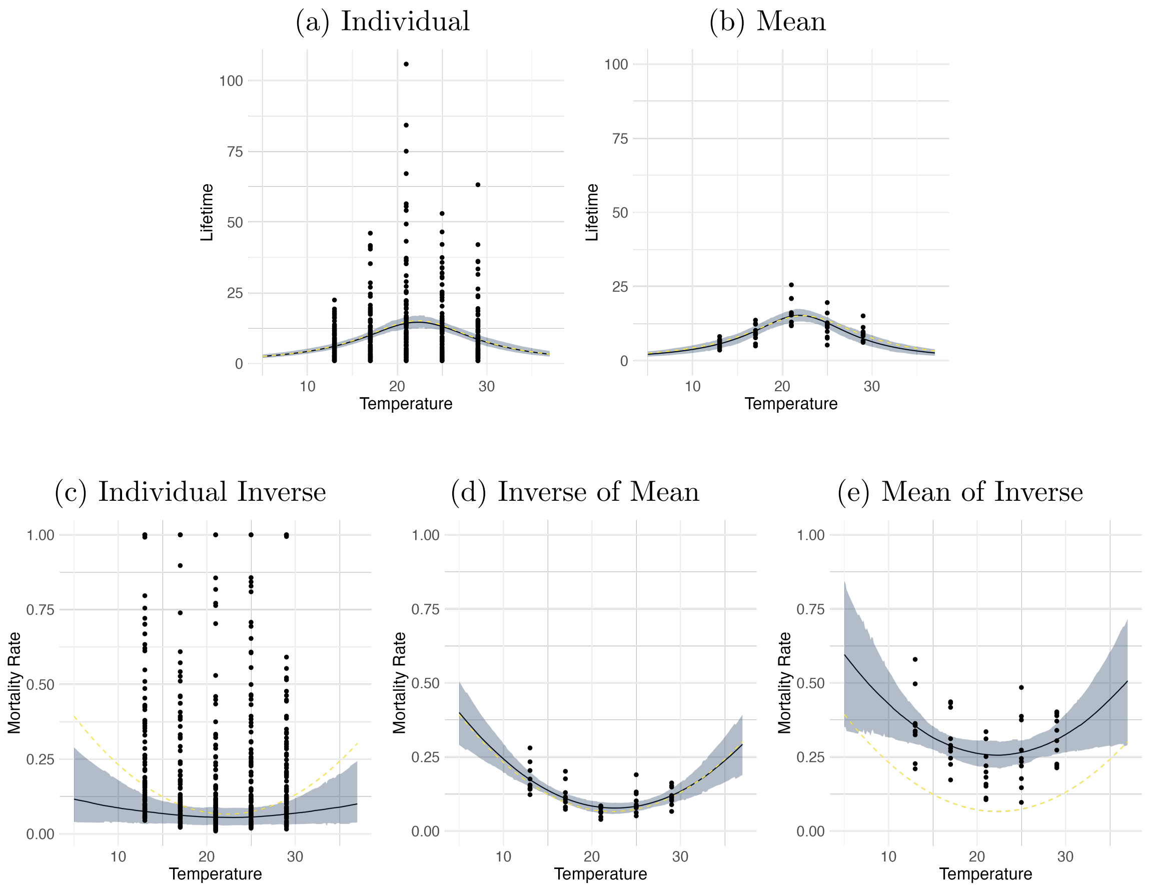


Figure S3: Comparison of the fitted TPC (median lifetime or mortality rate) when using data generated from an exponential distribution with a narrow temperature interval across data transformations. S3a Individual data points; S3b mean data; S3c individual inverse data; S3d IoM data; and S3e MoI data. Curves were made by evaluating samples of the posterior function across a temperature range from 5°C to 37°C. The median response at each temperature is shown as the solid lines, while the true curve is a dashed line. The 95% HDI bounds of the response at each temperature (TPC) were also plotted as the grey ribbon. The corresponding transformed data used to fit the TPC is included.

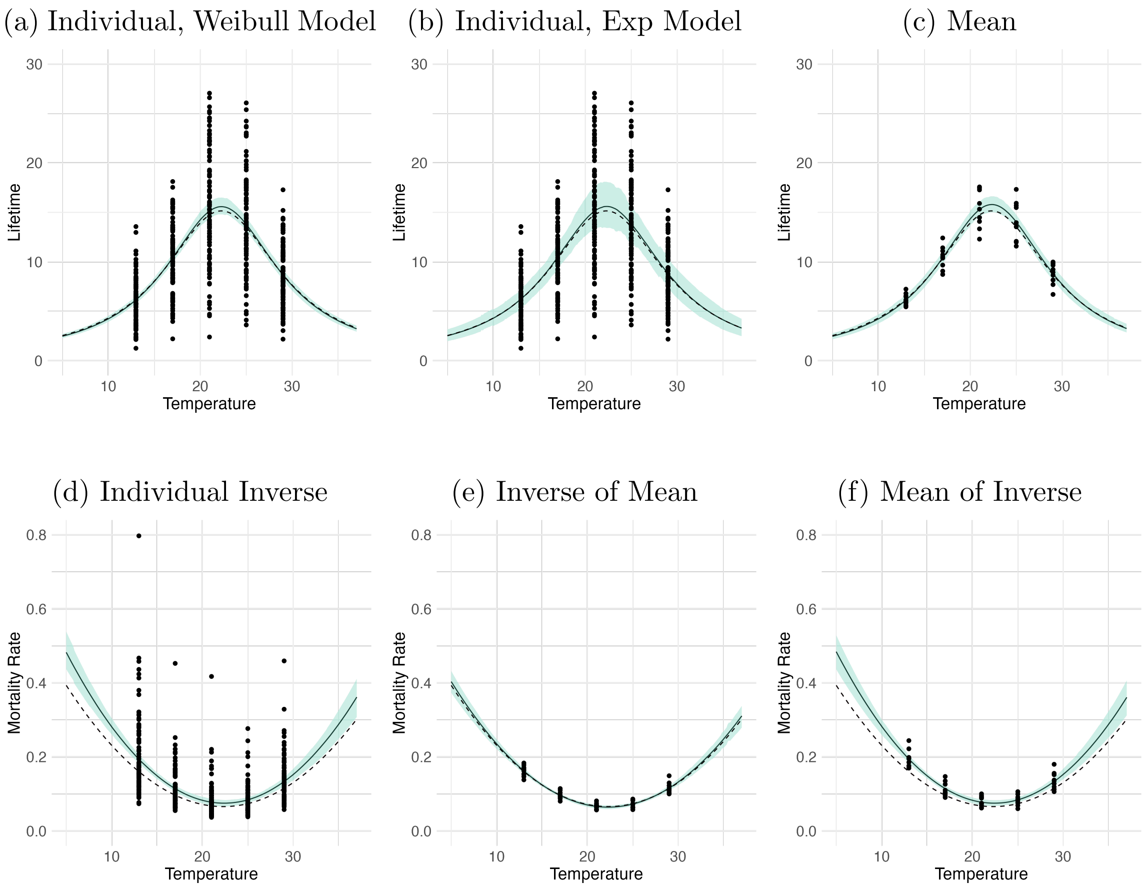


Figure S4: Comparison of the fitted TPC (median lifetime or mortality rate) when using data generated from a Weibull distribution with a narrow temperature interval across data transformations. S4a Individual data points with a Weibull model; S4b individual data points with an exponential model; S4c mean data; S4d individual inverse data; S4e IoM data; and S4f MoI data. Curves were made by evaluating samples of the posterior function across a temperature range from 5°C to 37°C. The median response at each temperature is shown as the solid lines, while the true curve is a dashed line. The 95% HDI bounds of the response at each temperature (TPC) were also plotted as the green ribbon. The corresponding transformed data used to fit the TPC is included.

#### 4.3 Extreme Temperature HDI Plots

This subsection gives a visual of how we obtained the upper and lower critical temperatures where the lifetime reaches one day. For all methods, estimated lifetime curves were created from posterior samples. When using a form of inverted data, we are modeling the mortality rate curve, so this was then inverted to obtain the lifetime curve. We then evaluated each function to find the estimated temperatures where the response was equal to one. One each plot, there is a horizontal green line at a lifetime of 1, meaning we are looking for the values where it intersects the true curve. In addition to this, we have circled where these intersections are. For each experiment, we also include a close up of the plot made using the mean data, so the difference in where the true curve and the estimated curve reach one day is clear.

##### 4.3.1 Wide

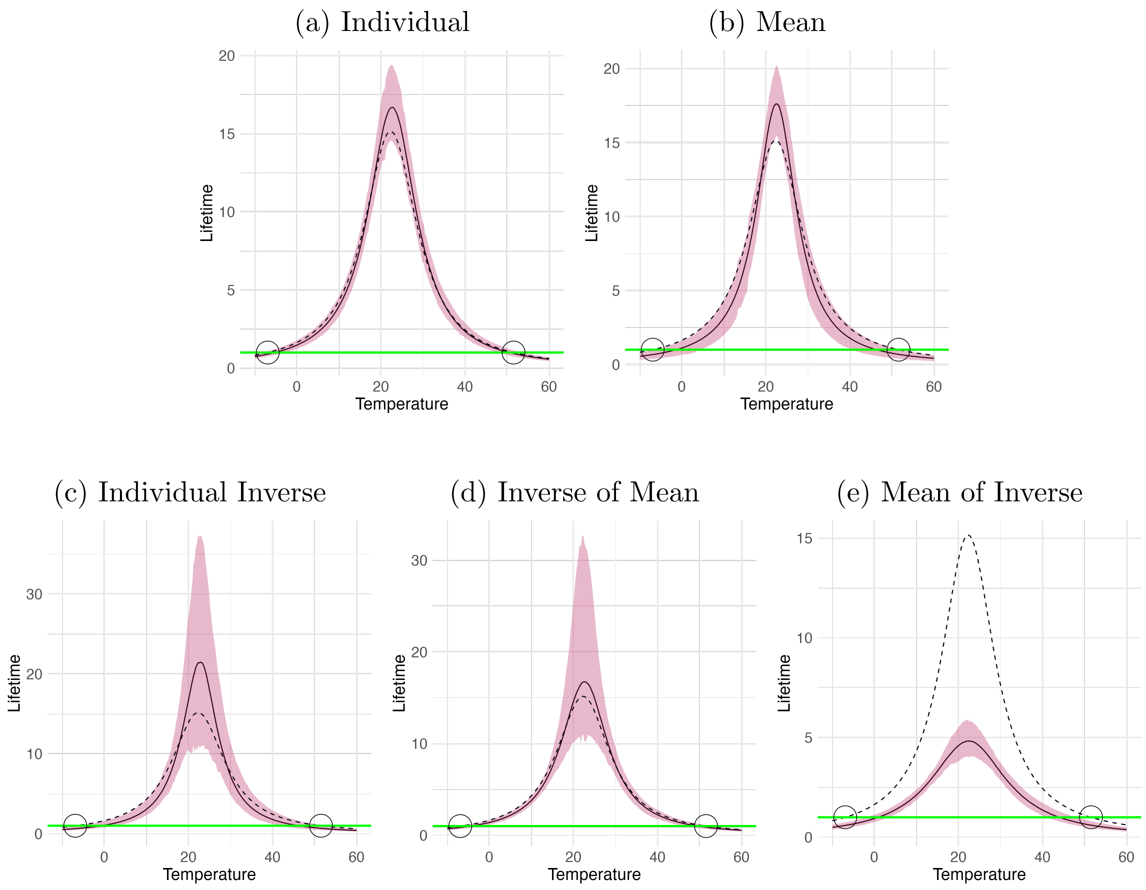


Figure S5: Median lifetime TPC when using data generated from an exponential distribution with a wide temperature interval across data transformations. S5a Individual data points; S5b mean data; S5c individual inverse data; S5d IoM data; and S5e MoI data. Curves were made by evaluating samples of the posterior function across a temperature range from -10°C to 60°C. The median response at each temperature is shown as the solid lines, while the true curve is a dashed line. The 95% HDI bounds of the response at each temperature (TPC) were also plotted as the pink ribbon. A horizontal green line is shown at a lifetime of 1 the indicate the critical temperatures. We circle these intersections where the lifetime value of the true curve is one day.

Figure S6: Inverse of Mean Zoom

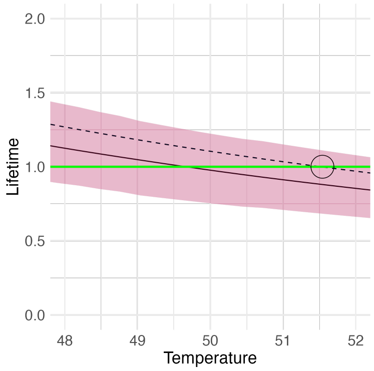


Figure S7: A zoomed in view of the upper critical temperature from Figure S5d. The circle highlights the intersection of the true curve (dashed line) and the horizontal line at 1. This intersection is the true upper critical temperature, or the temperature when lifetime reaches one day using the true curve. We can see that the estimated curve (solid line) intersects the horizontal line at a temperature slightly lower than the true value.

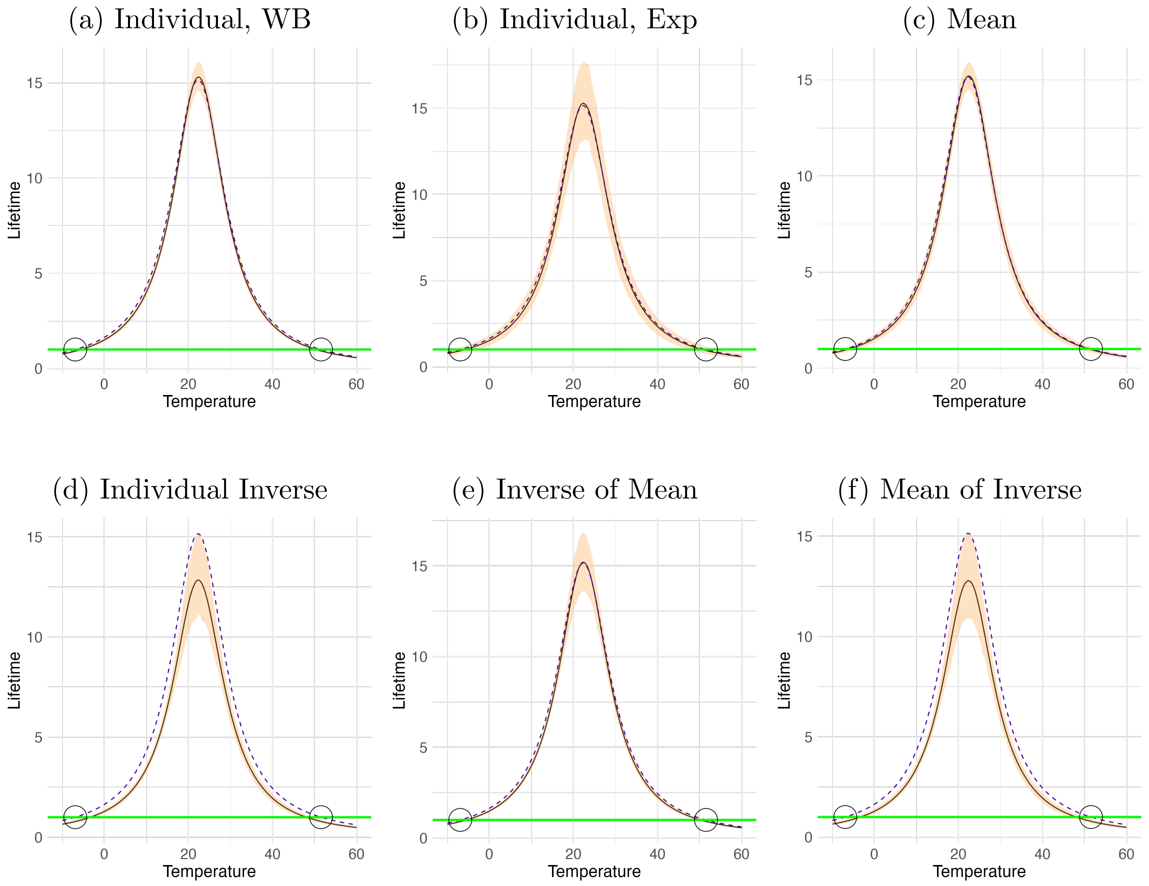


Figure S8: Median lifetime TPC when using data generated from a Weibull distribution with a wide temperature interval across data transformations. S8a Individual data points with a Weibull model; S8b individual data points with an exponential model; S8c mean data; S8d individual inverse data; S8e IoM data; and S8f MoI data. Curves were made by evaluating samples of the posterior function across a temperature range from -10°C to 60°C. The median response at each temperature is shown as the solid lines, while the true curve is a dashed line. The 95% HDI bounds of the response at each temperature (TPC) were also plotted as the orange ribbon. A horizontal green line is shown at a lifetime of 1 the indicate the critical temperatures. We circle these intersections where the lifetime value of the true curve is one day.

Figure S9: Inverse of Mean Zoom

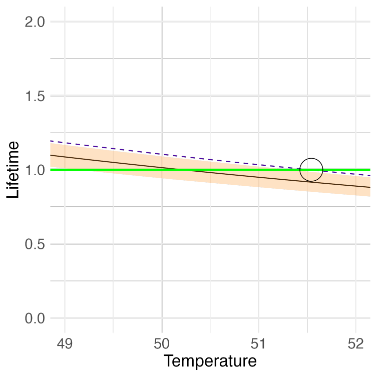


Figure S10: A zoomed in view of the upper critical temperature from Figure S8e. The circle highlights the intersection of the true curve (dashed line) and the horizontal line at 1. This intersection is the true upper critical temperature, or the temperature when lifetime reaches one day using the true curve. We can see that the estimated curve (solid line) intersects the horizontal line at a temperature slightly lower than the true value.

##### 4.3.2 Narrow

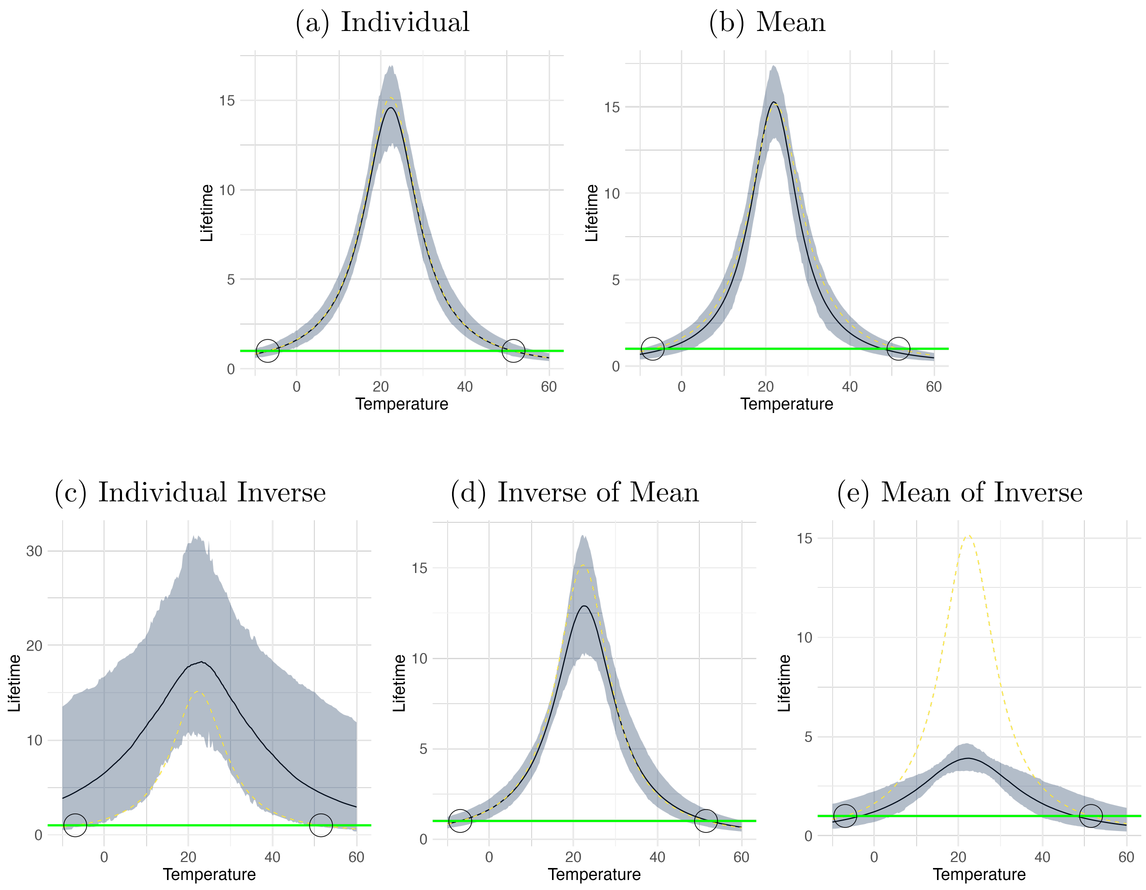


Figure S11: Median lifetime TPC when using data generated from an exponential distribution with a narrow temperature interval across data transformations. S11a Individual data points; S11b mean data; S11c individual inverse data; S11d IoM data; and S11e MoI data. Curves were made by evaluating samples of the posterior function across a temperature range from -10°C to 60°C. The median response at each temperature is shown as the solid lines, while the true curve is a dashed line. The 95% HDI bounds of the response at each temperature (TPC) were also plotted as the grey ribbon. A horizontal green line is shown at a lifetime of 1 the indicate the critical temperatures. We circle these intersections where the lifetime value of the true curve is one day.

Figure S12: Inverse of Mean Zoom

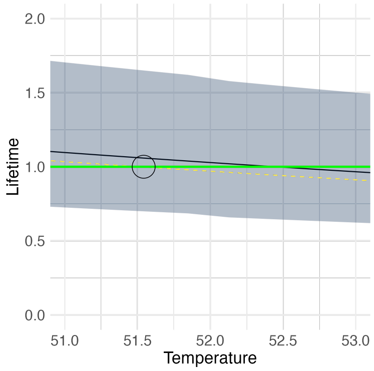


Figure S13: A zoomed in view of the upper critical temperature from Figure S11d. The circle highlights the intersection of the true curve (dashed line) and the horizontal line at 1. This intersection is the true upper critical temperature, or the temperature when lifetime reaches one day using the true curve. We can see that the estimated curve (solid line) intersects the horizontal line at a temperature slightly higher than the true value.

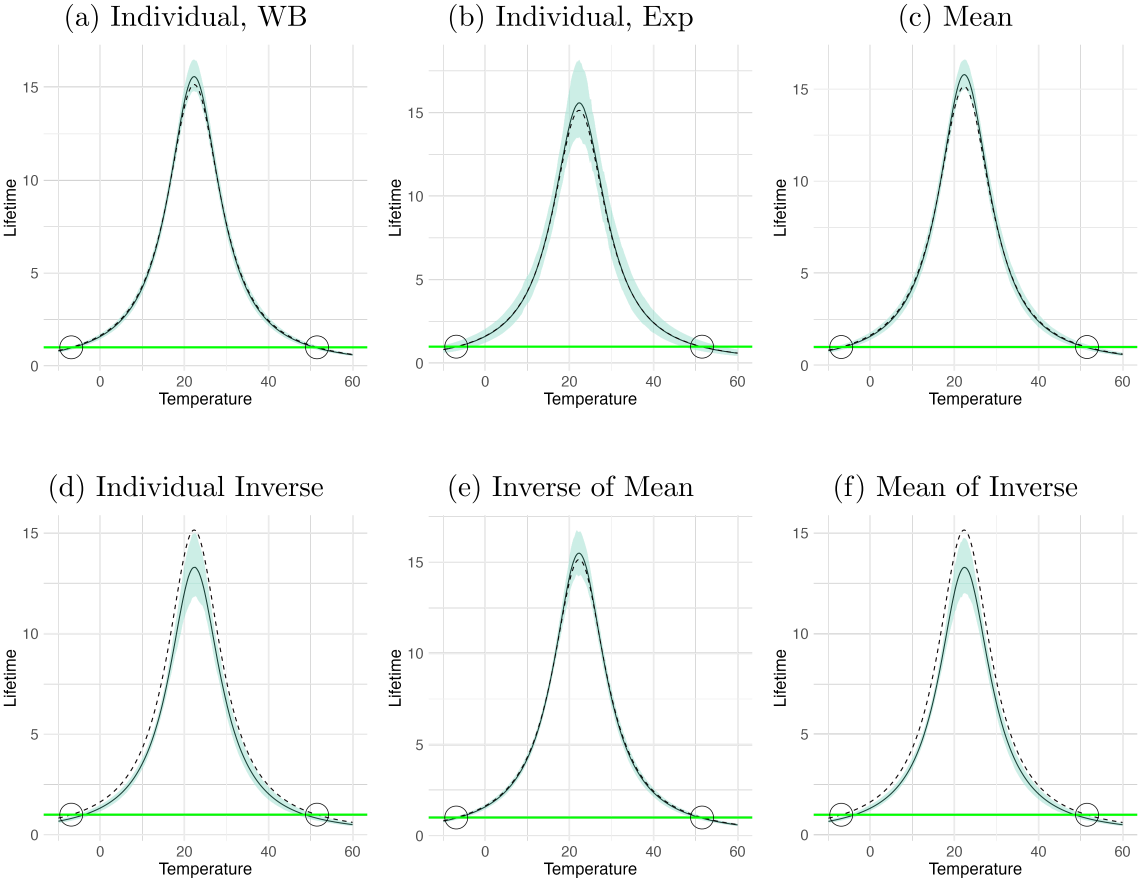


Figure S14: Median lifetime TPC when using data generated from a Weibull distribution with a narrow temperature interval across data transformations. S14a Individual data points with a Weibull model; S14b individual data points with an exponential model; S14c mean data; S14d individual inverse data; S14e IoM data; and S14f MoI data. Curves were made by evaluating samples of the posterior function across a temperature range from -10°C to 60°C. The median response at each temperature is shown as the solid lines, while the true curve is a dashed line. The 95% HDI bounds of the response at each temperature (TPC) were also plotted as the green ribbon. A horizontal green line is shown at a lifetime of 1 the indicate the critical temperatures. We circle these intersections where the lifetime value of the true curve is one day.

Figure S15: Inverse of Mean Zoom

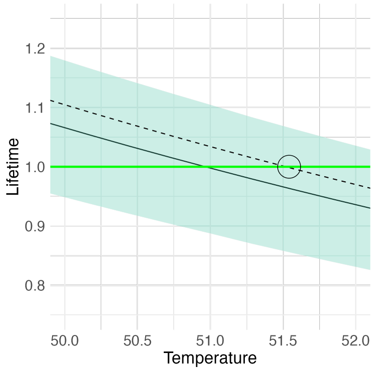


Figure S16: A zoomed in view of the upper critical temperature from Figure S14e. The circle highlights the intersection of the true curve (dashed line) and the horizontal line at 1. This intersection is the upper critical temperature, or the temperature when lifetime reaches one day. We can see that the estimated curve (solid line) intersects the horizontal line at a temperature slightly lower than the true value.

#### 4.4 Curve Shape Evaluation

##### 4.4.1 Wide

This subsubsection includes joint boxplots of the optimal temperature and maximum lifetime estimates.

(a) Exponential Data

(b) Weibull Data

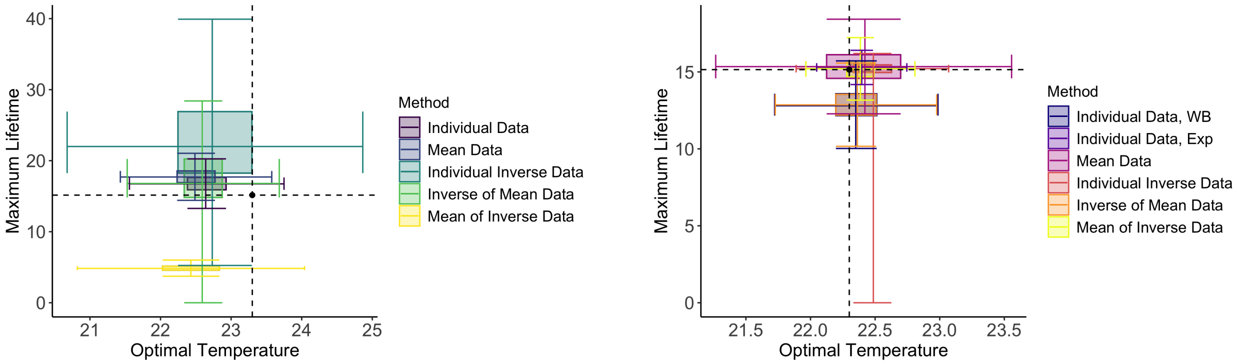


Figure S17: Boxplot summarizing the joint posterior distribution of the optimal temperature and maximum lifetime for the different data transformations when using exponential (S17a) and Weibull (S17b) data with a wide temperature interval. The black point at the intersection of the dashed lines represents the true values, derived from Equation ??. Outliers not shown.

###### 4.4.2 Narrow

This subsubsection includes both separate and joint boxplots of the optimal temperature and maximum lifetime estimates and joint boxplots for the estimates of the critical temperatures.

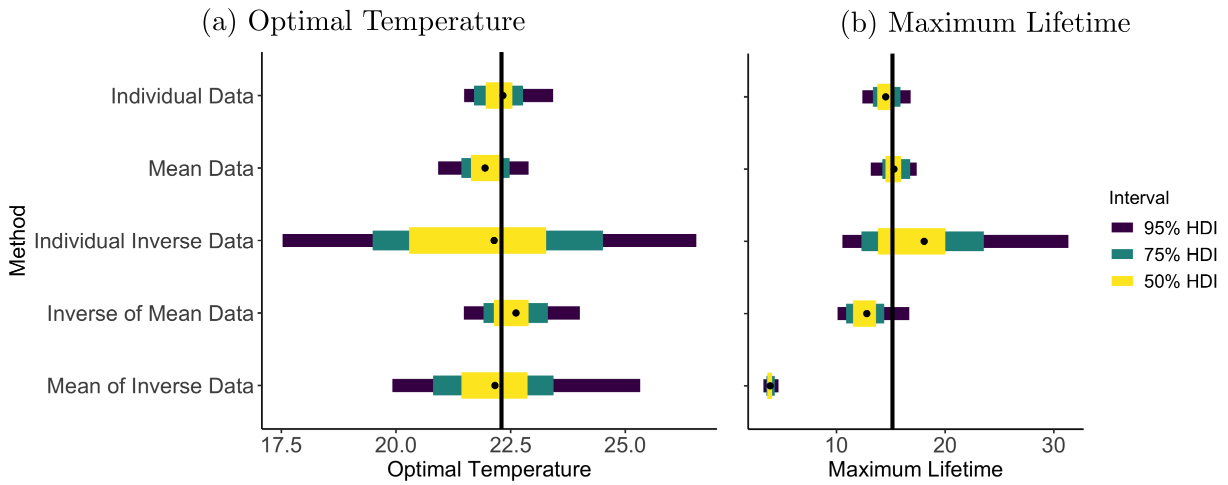


Figure S18: Posterior estimates and uncertainty of (S18a) the estimated optimal temperature, $T_{opt}$ and (S18b) of the maximum lifetime, $L_{Max}$, across data transformations for the single exponential dataset visualized in Figure S3. The vertical black lines represent the true values, the black dot represents the median of the posterior samples of each method, and the yellow, teal, and purple bars represent the 50%, 75%, and 95% HDI bounds, respectively.

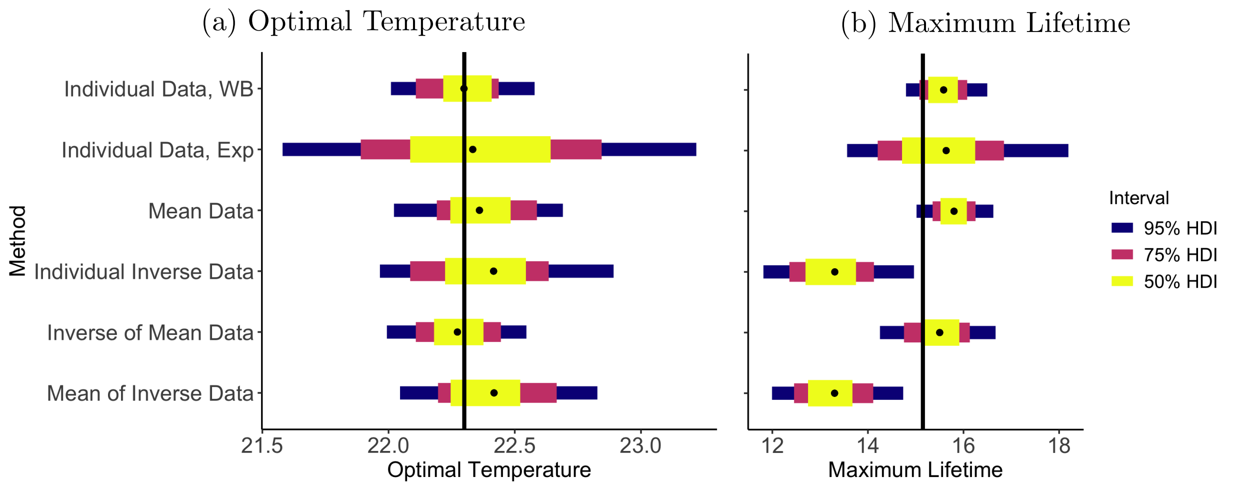


Figure S19: Posterior estimates and uncertainty of (S19a) the estimated optimal temperature, $T_{opt}$ and (S19b) of the maximum lifetime, $L_{Max}$, across data transformations for the single exponential dataset visualized in Figure S4. The vertical black lines represent the true values, the black dot represents the median of the posterior samples of each method, and the yellow, teal, and purple bars represent the 50%, 75%, and 95% HDI bounds, respectively.

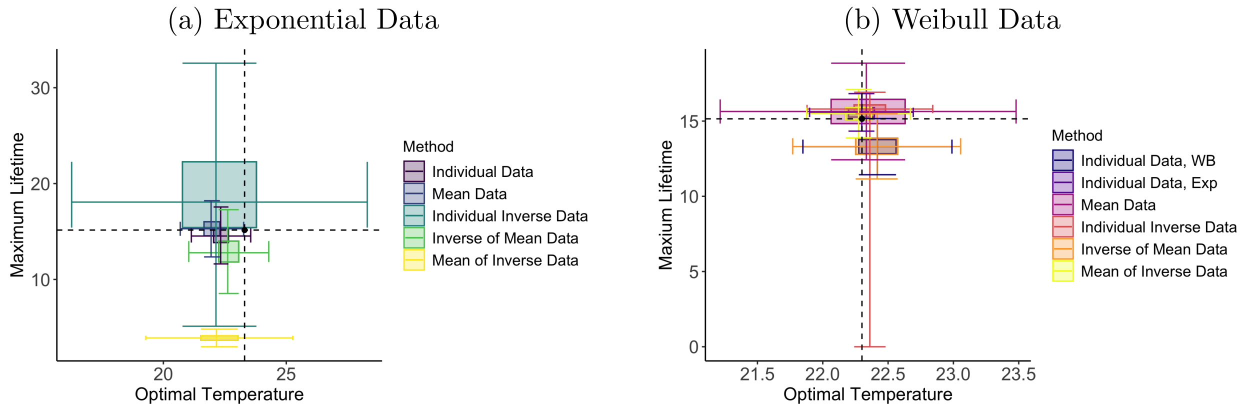


Figure S20: Boxplot summarizing the joint posterior distribution of the optimal temperature and maximum lifetime for the different data transformations when using exponential (S20a) and Weibull (S20b) data with a narrow temperature interval. The black point at the intersection of the dashed lines represents the true values, derived from Equation ??. Outliers not shown.

(a) Exponential Data (b) Weibull Data

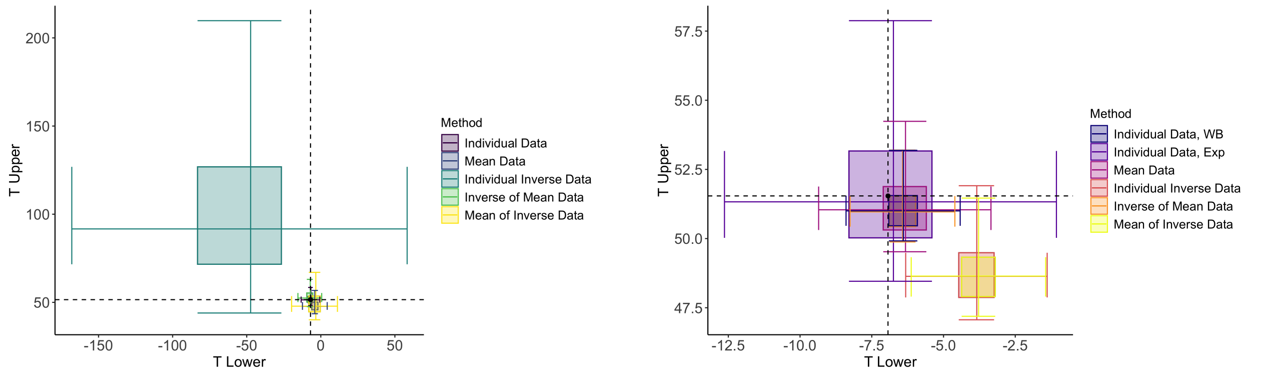


Figure S21: Boxplot summarizing the joint posterior distribution of the estimated lower ($T_{\text{Lower}}$) and upper ($T_{\text{Upper}}$) critical temperatures (where lifetime is less than one day) for the different data transformations when using exponential (S21a) and Weibull (S21b) data with a narrow temperature interval. The black point at the intersection of the dashed lines represents the true values, derived from Equation **??**. Outliers not shown.

#### 4.5 Uncertainty

##### 4.5.1 Wide

| Method | Median($T_{\text{opt}}$) | Times($T_{\text{opt}}$) | Median($L_{\text{Max}}$) | Times($L_{\text{Max}}$) |
|---|---|---|---|---|
| Individual Data | 1.75 | 1.00 | 4.46 | 1.00 |
| Mean Data | 2.04 | 1.17 | 4.11 | 0.921 |
| Individual Inverse Data | 4.18 | 2.39 | 28.4 | 6.37 |
| Inverse of Mean Data | 1.87 | 1.07 | 16.6 | 3.71 |
| Mean of Inverse Data | 2.72 | 1.56 | 1.53 | 0.343 |

Table S5: Median length of the 95% HDI intervals of posterior samples of parameters $T_{\text{opt}}$ and the inverse of parameter $c$ ($L_{\text{Max}}$) for the methods across all datasets. Values are also expressed as multiples of the uncertainty obtained when using individual-level data (e.g. mean data having a value of 1.17 in the "Times($T_{\text{opt}}$)" column indicates that the uncertainty is 1.17 times that of the individual data). Made using data generated from an exponential distribution with a wide temperature interval.

| Method | Median(Low) | Times(Low) | Median(High) | Times(High) |
|---|---|---|---|---|
| Individual Data | 9.19 | 1.00 | 9.85 | 1.00 |
| Mean Data | 16.7 | 1.82 | 17.4 | 1.76 |
| Individual Inverse Data | 22.1 | 2.40 | 25.2 | 2.56 |
| Inverse of Mean Data | 8.45 | 0.919 | 10.0 | 1.02 |
| Mean of Inverse Data | 9.08 | 0.987 | 11.2 | 1.14 |

Table S6: Median length of the uncertainty in $T_{\text{Upper}}$ and $T_{\text{Lower}}$, calculated as the length of the boxplot whiskers created from evaluating joint posterior samples for when lifetime reaches one day for the methods across all datasets. Values are also expressed as multiples of the uncertainty obtained when using individual-level data (e.g. mean data having a value of 1.82 in the "Times(Low)" column indicates that the uncertainty is 1.82 times that of the individual data). Made using data generated from an exponential distribution with a wide temperature interval.

| Method | Median($T_{\text{opt}}$) | Times($T_{\text{opt}}$) | Median($L_{\text{Max}}$) | Times($L_{\text{Max}}$) |
|---|---|---|---|---|
| Individual Data, WB | 0.580 | 1.00 | 1.55 | 1.00 |
| Individual Data, Exp | 1.76 | 3.03 | 4.42 | 2.85 |
| Mean Data | 0.788 | 1.36 | 1.31 | 0.846 |
| Individual Inverse Data | 0.918 | 1.58 | 3.70 | 2.38 |
| Inverse of Mean Data | 0.602 | 1.04 | 2.91 | 1.88 |
| Mean of Inverse Data | 0.928 | 1.60 | 3.64 | 2.35 |

Table S7: Median length of the 95% HDI intervals of posterior samples of parameters $T_{\text{opt}}$ and the inverse of parameter $c$ ($L_{\text{Max}}$) for the methods across all datasets. Values are also expressed as multiples of the uncertainty obtained when using individual-level data (e.g. mean data having a value of 1.36 in the "Times($T_{\text{opt}}$)" column indicates that the uncertainty is 1.36 times that of the individual data). Made using data generated from a Weibull distribution with a wide temperature interval.

| Method | Median(Low) | Times(Low) | Median(High) | Times(High) |
|---|---|---|---|---|
| Individual Data, WB | 3.12 | 1.00 | 3.33 | 1.00 |
| Individual Data, Exp | 9.28 | 2.98 | 9.90 | 2.98 |
| Mean Data | 5.75 | 1.85 | 5.98 | 1.80 |
| Individual Inverse Data | 3.85 | 1.23 | 4.68 | 1.41 |
| Inverse of Mean Data | 2.72 | 0.873 | 3.24 | 0.974 |
| Mean of Inverse Data | 3.79 | 1.22 | 4.64 | 1.39 |

Table S8: Median length of the uncertainty in $T_{\text{Upper}}$ and $T_{\text{Lower}}$, calculated as the length of the boxplot whiskers created from evaluating joint posterior samples for when lifetime reaches one day for the methods across all datasets. Values are also expressed as multiples of the uncertainty obtained when using individual-level data (e.g. mean data having a value of 1.85 in the "Times(Low)" column indicates that the uncertainty is 1.85 times that of the individual data). Made using data generated from a Weibull distribution with a wide temperature interval.

##### 4.5.2 Narrow

| Method | Median($T_{\text{opt}}$) | Times($T_{\text{opt}}$) | Median($L_{\text{Max}}$) | Times($L_{\text{Max}}$) |
|---|---|---|---|---|
| Individual Data | 1.87 | 1.00 | 4.63 | 1.00 |
| Mean Data | 2.08 | 1.11 | 4.07 | 0.879 |
| Individual Inverse Data | 9.11 | 4.86 | 29.6 | 6.40 |
| Inverse of Mean Data | 2.20 | 1.17 | 8.57 | 1.85 |
| Mean of Inverse Data | 4.51 | 2.41 | 1.62 | 0.349 |

Table S9: Median length of the 95% HDI intervals of posterior samples of parameters $T_{\text{opt}}$ and the inverse of parameter $c$ ($L_{\text{Max}}$) for the methods across all datasets. Values are also expressed as multiples of the uncertainty obtained when using individual-level data (e.g. mean data having a value of 1.11 in the "Times($T_{\text{opt}}$)" column indicates that the uncertainty is 1.11 times that of the individual data). Made using data generated from an exponential distribution with a narrow temperature interval.

| Method | Median(Low) | Times(Low) | Median(High) | Times(High) |
|---|---|---|---|---|
| Individual Data | 13.4 | 1.00 | 14.8 | 1.00 |
| Mean Data | 17.2 | 1.33 | 18.2 | 1.23 |
| Individual Inverse Data | 203 | 15.2 | 205 | 13.8 |
| Inverse of Mean Data | 13.7 | 1.02 | 16.2 | 1.09 |
| Mean of Inverse Data | 20.2 | 1.51 | 26.0 | 1.75 |

Table S10: Median length of the uncertainty in $T_{\text{Upper}}$ and $T_{\text{Lower}}$, calculated as the length of the boxplot whiskers created from evaluating joint posterior samples for when lifetime reaches one day for the methods across all datasets. Values are also expressed as multiples of the uncertainty obtained when using individual-level data (e.g. mean data having a value of 1.33 in the "Times(Low)" column indicates that the uncertainty is 1.33 times that of the individual data). Made using data generated from an exponential distribution with a narrow temperature interval.

| Method | Median($T_{\text{opt}}$) | Times($T_{\text{opt}}$) | Median($L_{\text{Max}}$) | Times($L_{\text{Max}}$) |
|---|---|---|---|---|
| Individual Data, WB | 0.580 | 1.00 | 1.65 | 1.00 |
| Individual Data, Exp | 1.83 | 3.15 | 4.72 | 2.86 |
| Mean Data | 0.730 | 0.88 | 1.45 | 1.26 |
| Individual Inverse Data | 1.04 | 1.80 | 3.21 | 1.95 |
| Inverse of Mean Data | 0.670 | 1.15 | 2.48 | 1.50 |
| Mean of Inverse Data | 1.08 | 1.86 | 3.26 | 1.97 |

Table S11: Median length of the 95% HDI intervals of posterior samples of parameters $T_{\text{opt}}$ and the inverse of parameter $c$ ($L_{\text{Max}}$) for the methods across all datasets. Values are also expressed as multiples of the uncertainty obtained when using individual-level data (e.g. mean data having a value of 0.88 in the "Times($T_{\text{opt}}$)" column indicates that the uncertainty is 0.88 times that of the individual data). Made using data generated from a Weibull distribution with a narrow temperature interval.

| Method | Median(Low) | Times(Low) | Median(High) | Times(High) |
|---|---|---|---|---|
| Individual Data, WB | 4.19 | 1.00 | 4.60 | 1.00 |
| Individual Data, Exp | 13.1 | 3.12 | 14.5 | 3.14 |
| Mean Data | 6.07 | 1.45 | 6.30 | 1.37 |
| Individual Inverse Data | 6.32 | 1.51 | 4.60 | 1.68 |
| Inverse of Mean Data | 4.32 | 1.03 | 5.21 | 1.13 |
| Mean of Inverse Data | 6.32 | 1.51 | 7.70 | 1.67 |

Table S12: Median length of the uncertainty in $T_{\text{Upper}}$ and $T_{\text{Lower}}$, calculated as the length of the boxplot whiskers created from evaluating joint posterior samples for when lifetime reaches one day for the methods across all datasets. Values are also expressed as multiples of the uncertainty obtained when using individual-level data (e.g. mean data having a value of 1.45 in the "Times(Low)" column indicates that the uncertainty is 1.45 times that of the individual data). Made using data generated from a Weibull distribution with a narrow temperature interval.

#### 4.6 Q-Q Plots

This subsection includes QQplots where the empirical distribution comes from the true points of the $100^{th}$ data set. For models using individual-level data and an exponential model, the theoretical distribution is an exponential distribution for both data-generating mechanisms. The rate parameter is the true quadratic mortality curve. When modeling individual-level data generated from a Weibull distribution with a Weibull model, the theoretical distribution is a Weibull distribution with the scale parameter including the true quadratic mortality rate curve and estimated shape parameter. For models using the mean, individual inverse, inverse of the mean, or mean of the inverse data, the theoretical distribution was a normal distribution truncated at 0, regardless of the data-generating mechanism. The mean of the truncated normal is the true quadratic mortality curve. Parameters are estimated using posterior medians.

##### 4.6.1 Wide

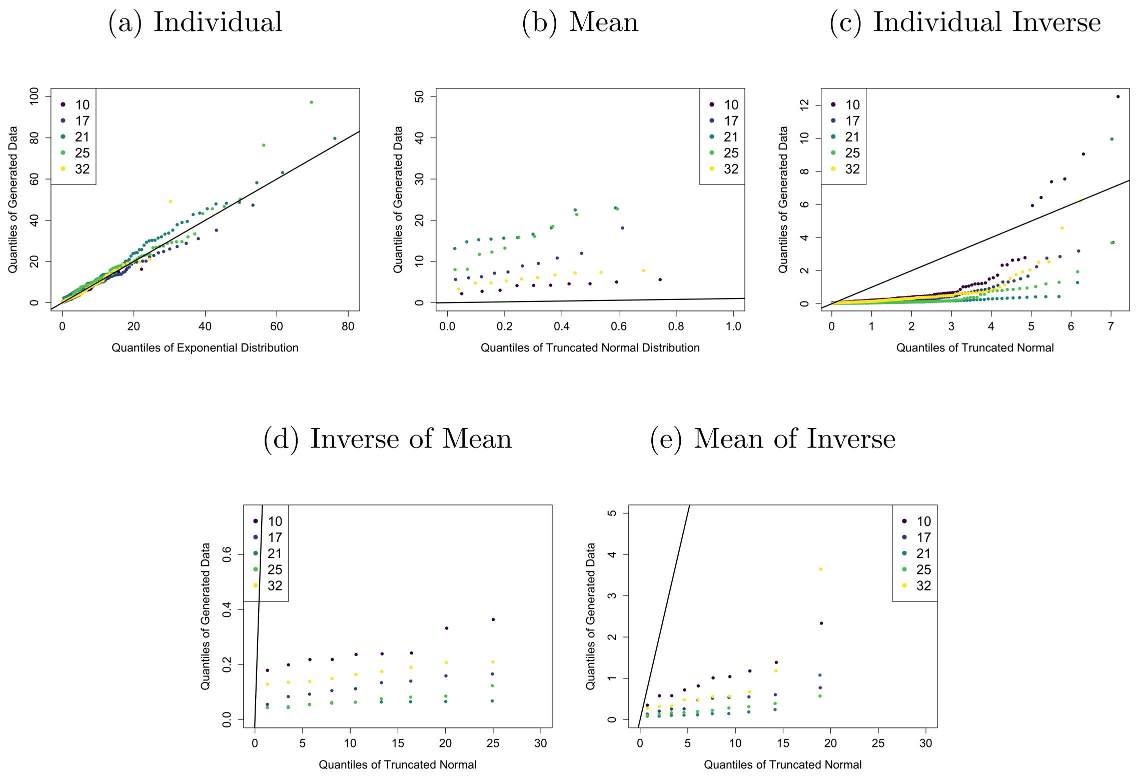


Figure S22: Quantile-quantile plots for individual-level data (S22a), mean data (S22b), individual inverse data (S22c), IoM data (S22d), and MoI data (S22e) using data generated from an exponential distribution with a wide temperature interval.

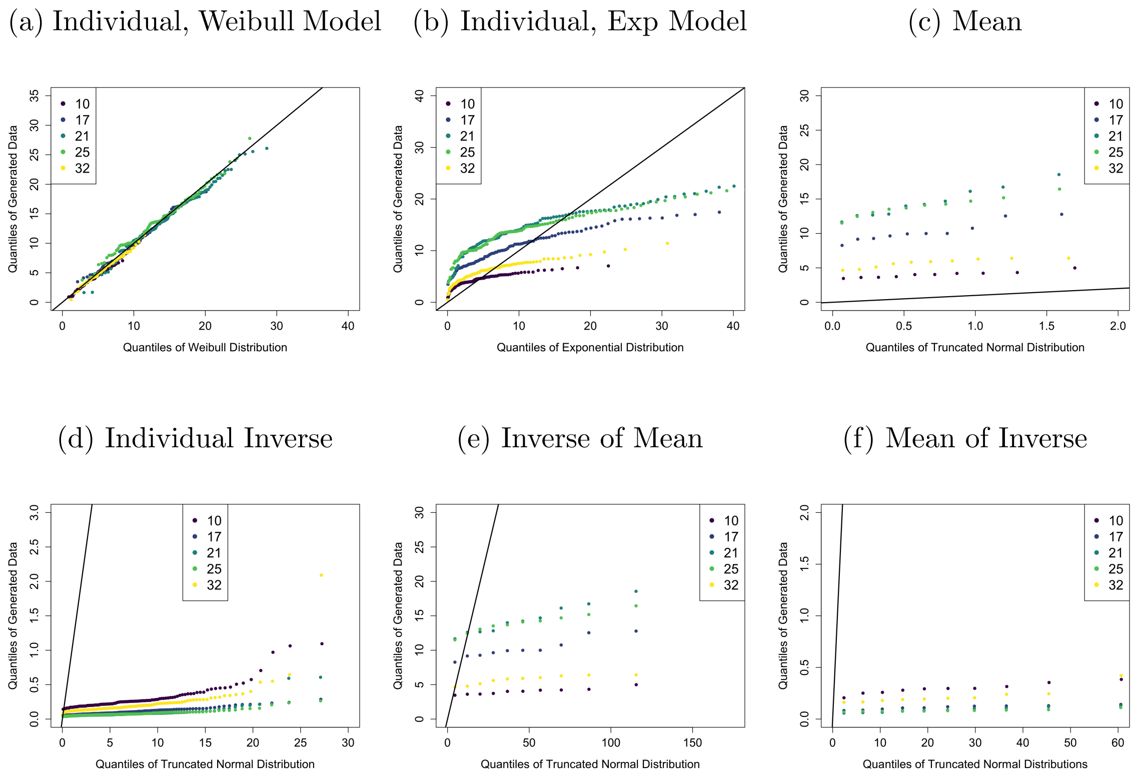


Figure S23: Quantile-quantile plots for individual-level data (S23a) modeled with a Weibull model, individual-level data (S23b) modeled with an exponential model, mean data (S23c), individual inverse data (S23d), IoM data (S23e), and MoI data (S23f) using data generated from a Weibull distribution with a wide temperature interval.

##### 4.6.2 Narrow

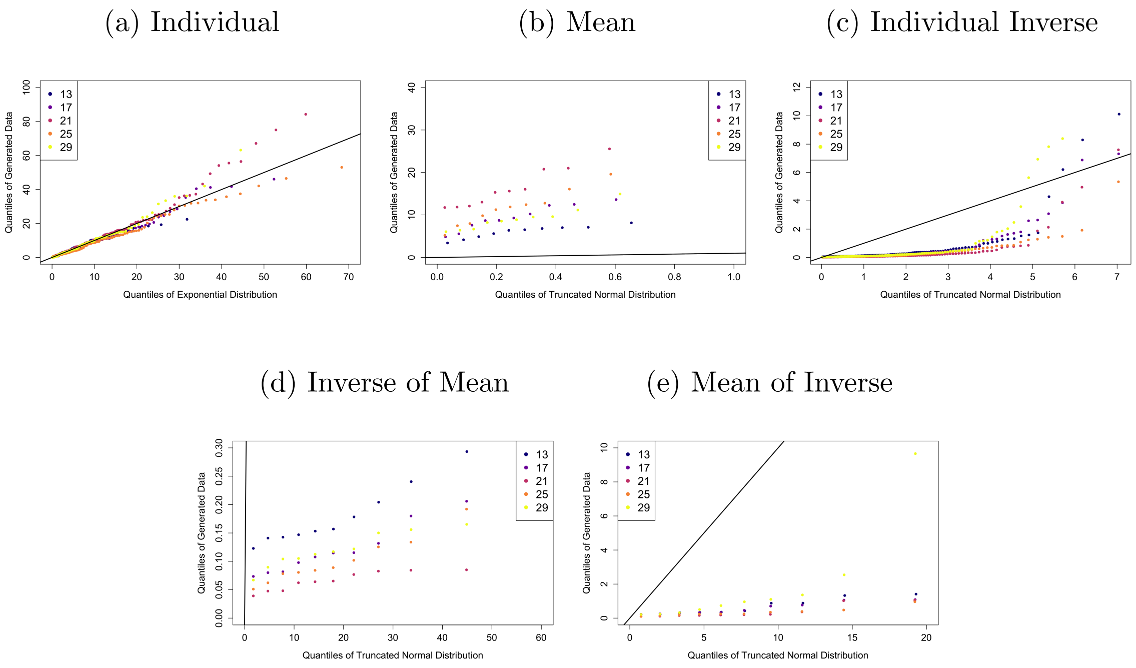


Figure S24: Quantile-quantile plots for individual-level data (S24a), mean data (S24b), individual inverse data (S24c), IoM data (S24d), and MoI data (S24e) using data generated from an exponential distribution with a wide temperature interval.

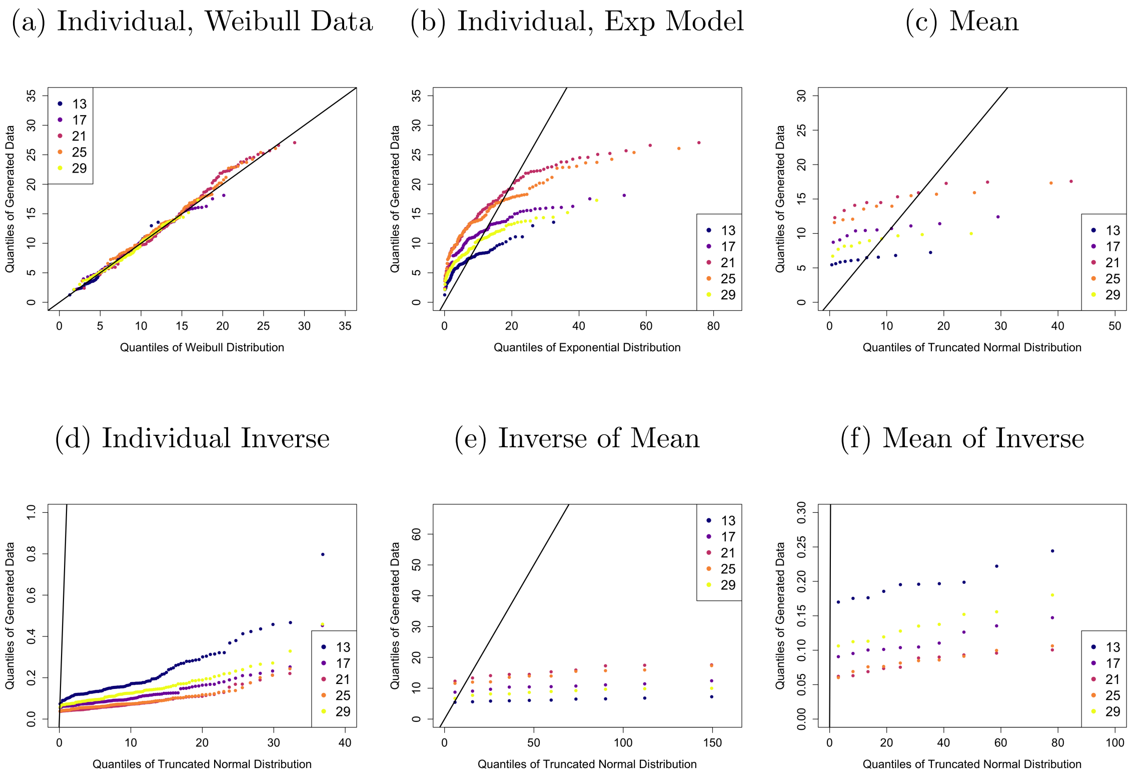


Figure S25: Quantile-quantile plots for individual-level data (S25a) modeled with a Weibull model, individual-level data (S25b) modeled with an exponential model, mean data (S25c), individual inverse data (S25d), IoM data (S25e), and MoI data (S25f) using data generated from a Weibull distribution with a narrow temperature interval.

#### 4.7 Narrow vs Wide Comparison

This subsection includes HDI plots of both mortality rate and lifetime curves for each method. The estimated curves from both a narrow and a wide temperature interval are included to compare the effects of experimental design.

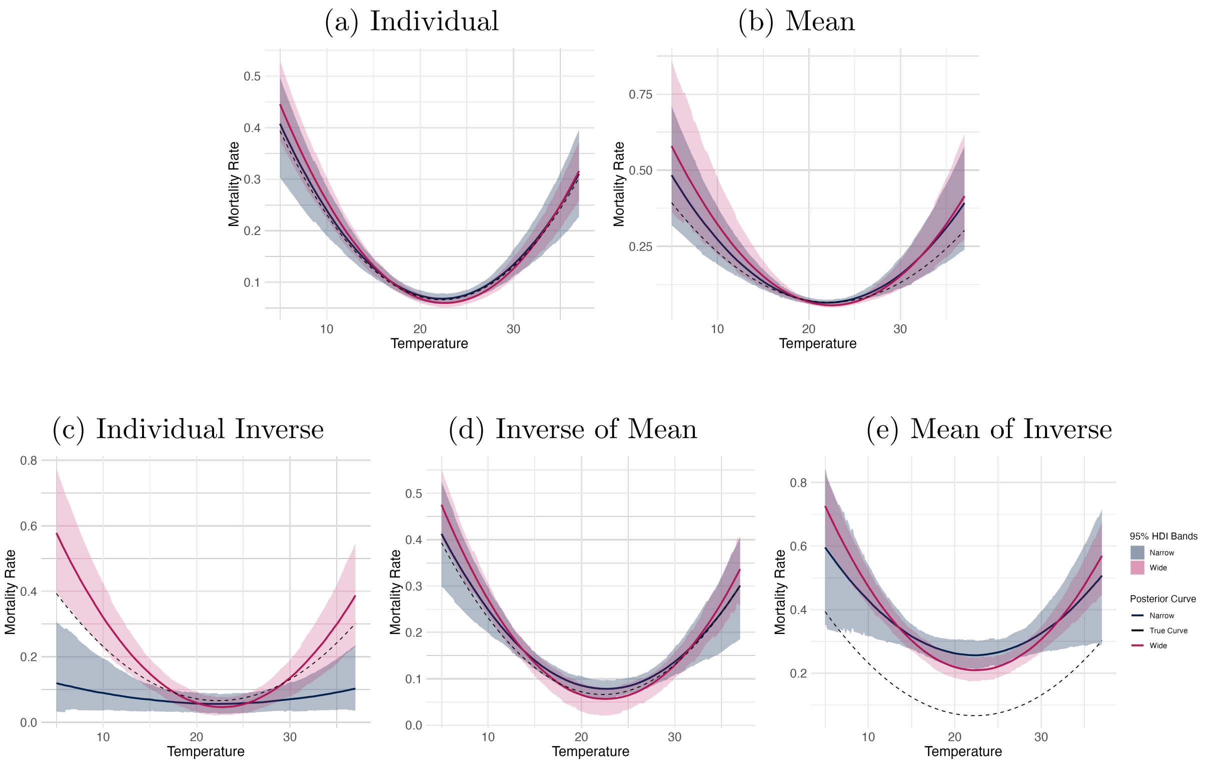


Figure S26: Comparison of the mortality rate TPC when using data generated from an exponential distribution with a wide (pink) and a narrow (grey) temperature interval across data transformations. S26a Individual data points; S26b mean data; S26c individual inverse data; S26d IoM data; and S26e MoI data. Curves were made by evaluating samples of the posterior function across a temperature range from 5°C to 37°C. The median response at each temperature is shown as the solid lines, while the true curve is a dashed line. The 95% HDI bounds of the response at each temperature (TPC) were also plotted as the pink (wide) and grey (narrow) ribbons.

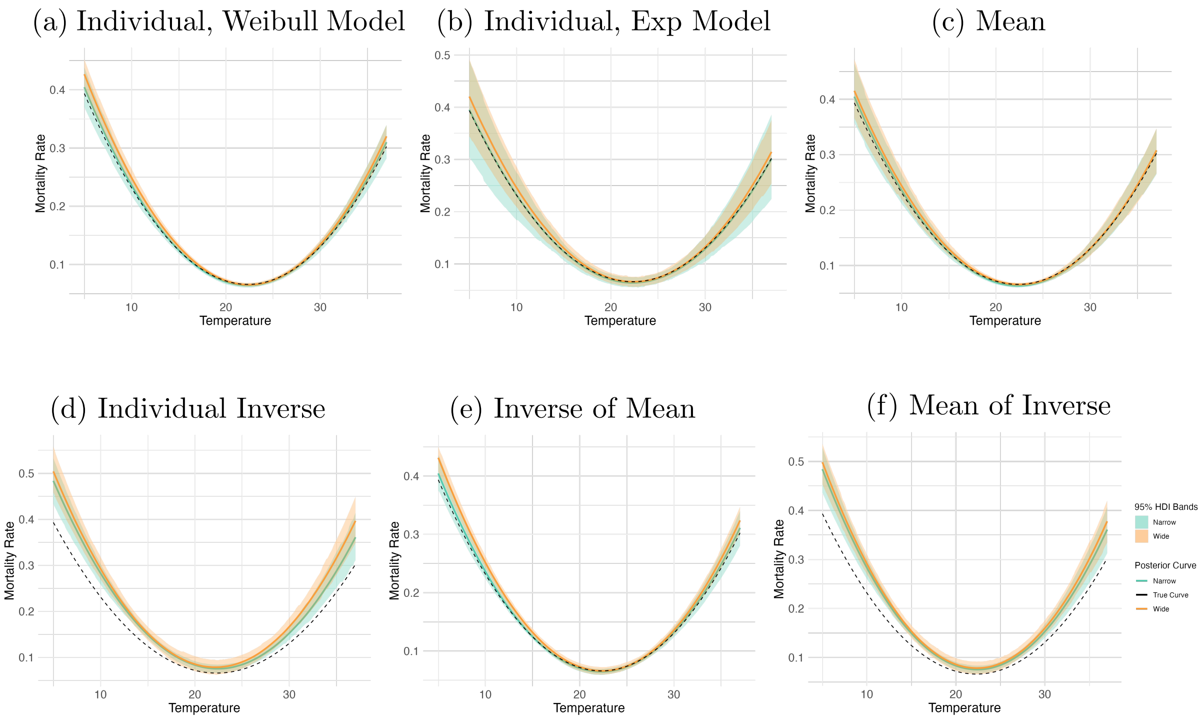


Figure S27: Comparison of the mortality rate TPC when using data generated from a Weibull distribution with a wide (orange) and a narrow (green) temperature interval across data transformations. S27a Individual data points with a Weibull model; S27b individual data points with an exponential model; S27c mean data; S27d individual inverse data; S27e IoM data; and S27f MoI data. Curves were made by evaluating samples of the posterior function across a temperature range from 5°C to 37°C. The median response at each temperature is shown as the solid lines, while the true curve is a dashed line. The 95% HDI bounds of the response at each temperature (TPC) were also plotted as the orange (wide) and green (narrow) ribbons.

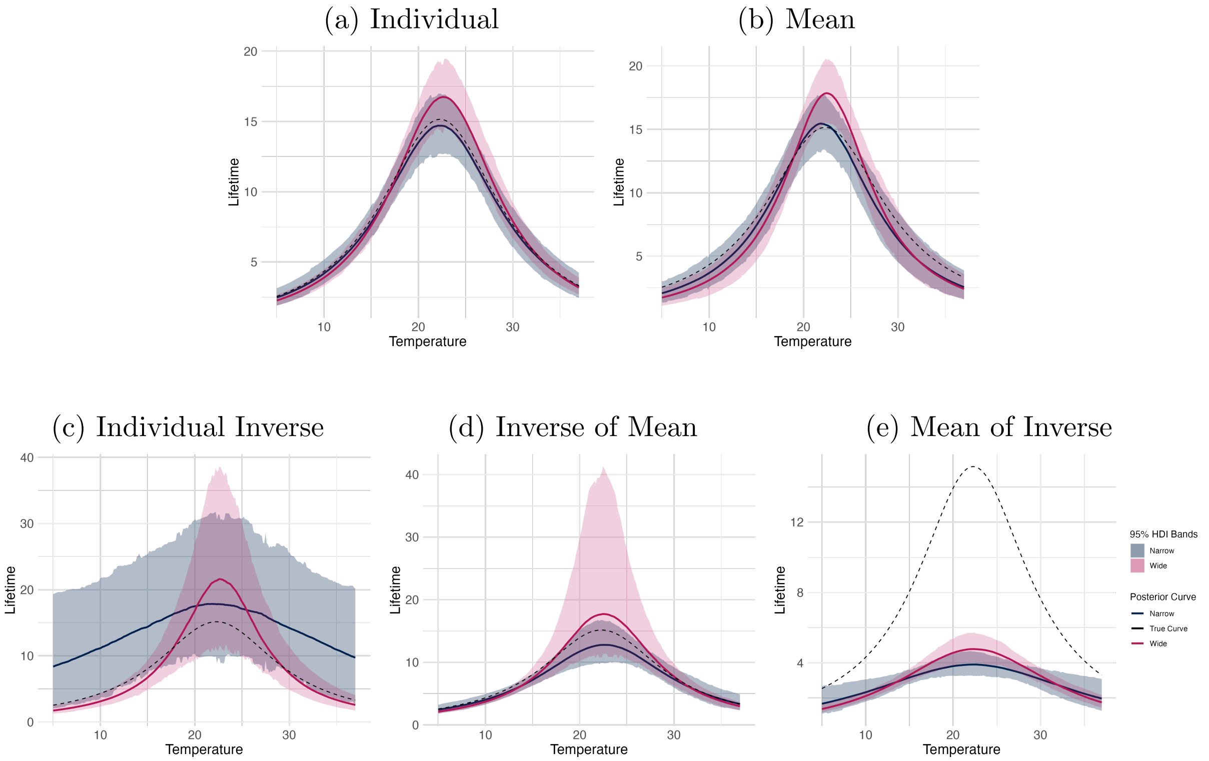


Figure S28: Comparison of the lifetime TPC when using data generated from an exponential distribution with a wide (pink) and a narrow (grey) temperature interval across data transformations. S28a Individual data points; S28b mean data; S28c individual inverse data; S28d IoM data; and S28e MoI data. Curves were made by evaluating samples of the posterior function across a temperature range from 5°C to 37°C. The median response at each temperature is shown as the solid lines, while the true curve is a dashed line. The 95% HDI bounds of the response at each temperature (TPC) were also plotted as the pink (wide) and grey (narrow) ribbons.

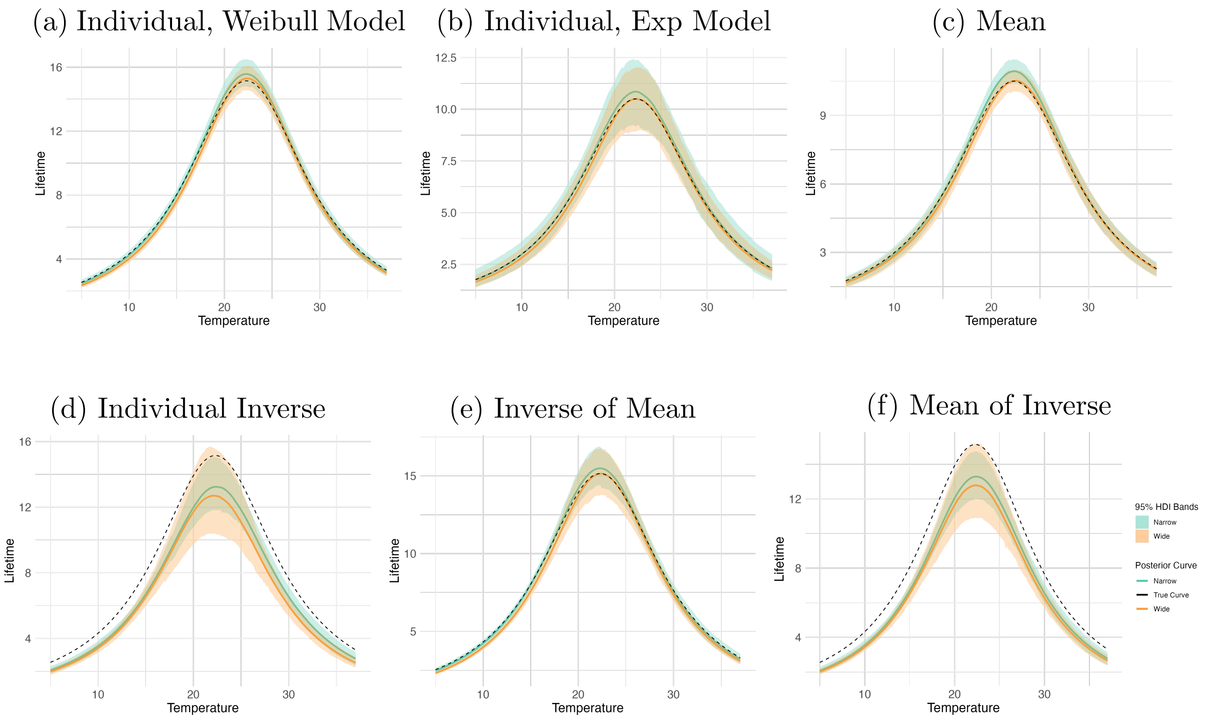


Figure S29: Comparison of the lifetime TPC when using data generated from a Weibull distribution with a wide (orange) and a narrow (green) temperature interval across data transformations. S29a Individual data points with a Weibull model; S29b individual data points with an exponential model; S29c mean data; S29d individual inverse data; S29e IoM data; and S29f MoI data. Curves were made by evaluating samples of the posterior function across a temperature range from 5°C to 37°C. The median response at each temperature is shown as the solid lines, while the true curve is a dashed line. The 95% HDI bounds of the response at each temperature (TPC) were also plotted as the orange (wide) and green (narrow) ribbons.

### 5 Model Logistics

This section includes the trace plots from one randomly selected dataset and model of each method for each temperature interval to show convergence.

#### 5.1 Exponential Data

##### 5.1.1 Individual Data

(a) Wide temperature interval

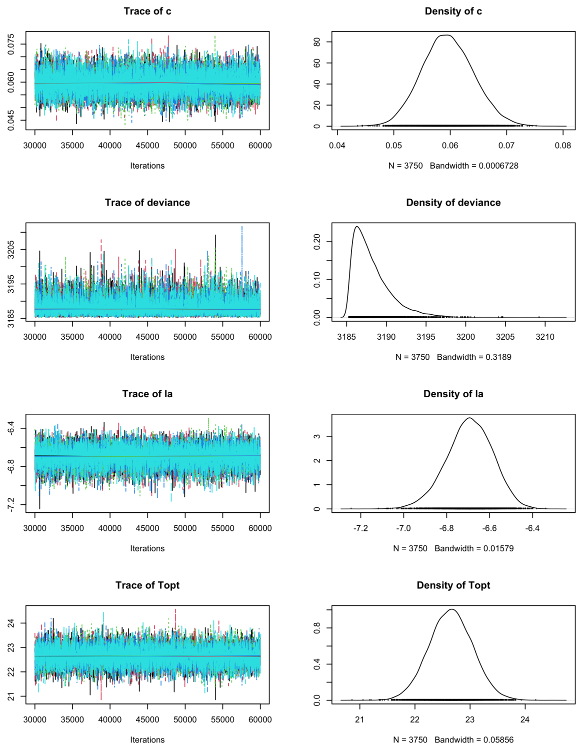


(b) Narrow temperature interval

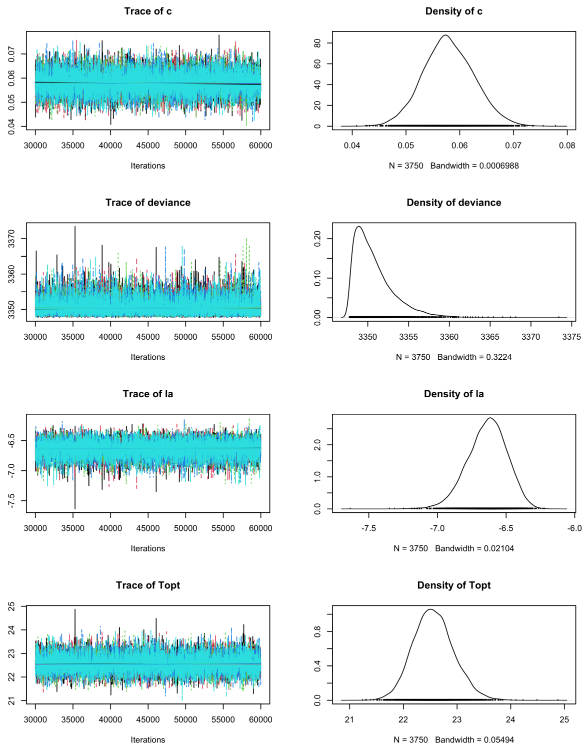


Figure S30: The trace plots for a randomly chosen dataset when modeling using individual-level data, generated from an exponential distribution with a wide temperature interval (S30a) and a narrow temperature interval (S30b).

##### 5.1.2 Mean Data

(a) Wide temperature interval (b) Narrow temperature interval

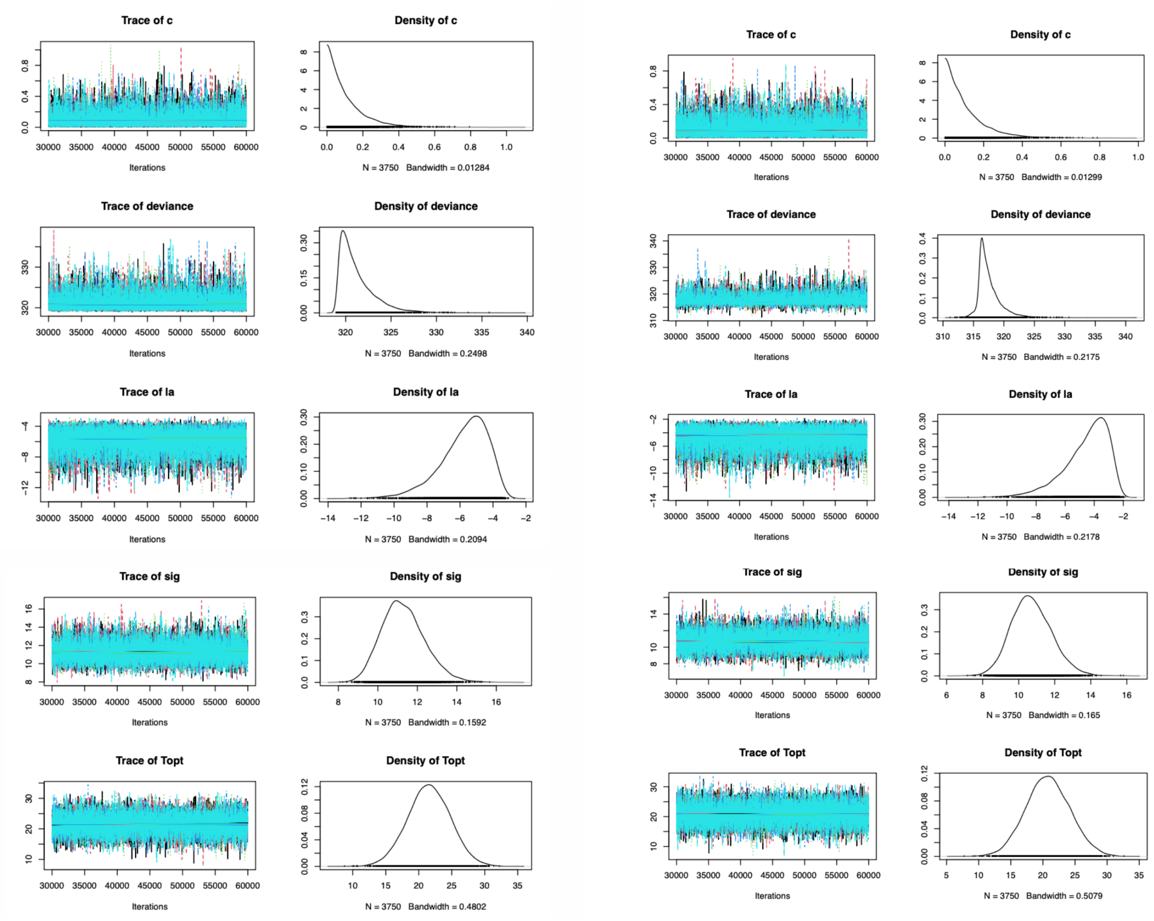


Figure S31: The trace plots for a randomly chosen dataset when modeling using mean data, generated from an exponential distribution with a wide temperature interval (S31a) and a narrow temperature interval (S31b).

##### 5.1.3 Inverse Data

(a) Wide temperature interval

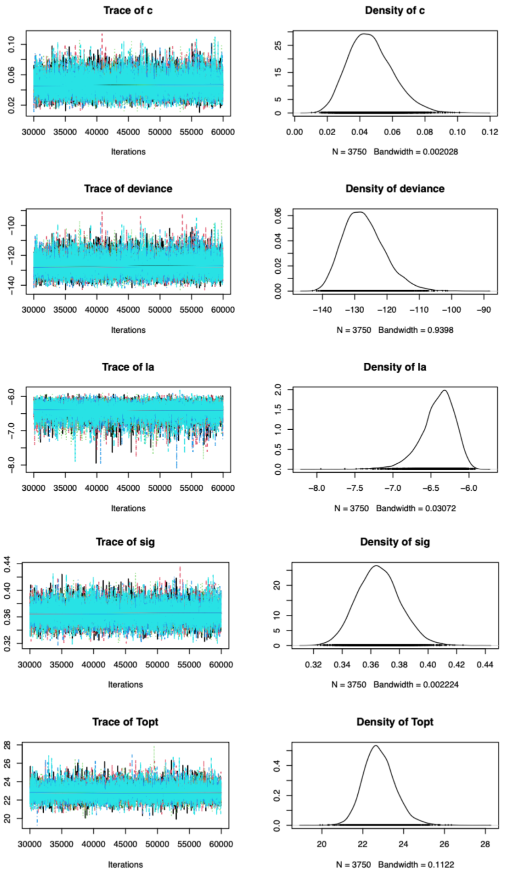


(b) Narrow temperature interval

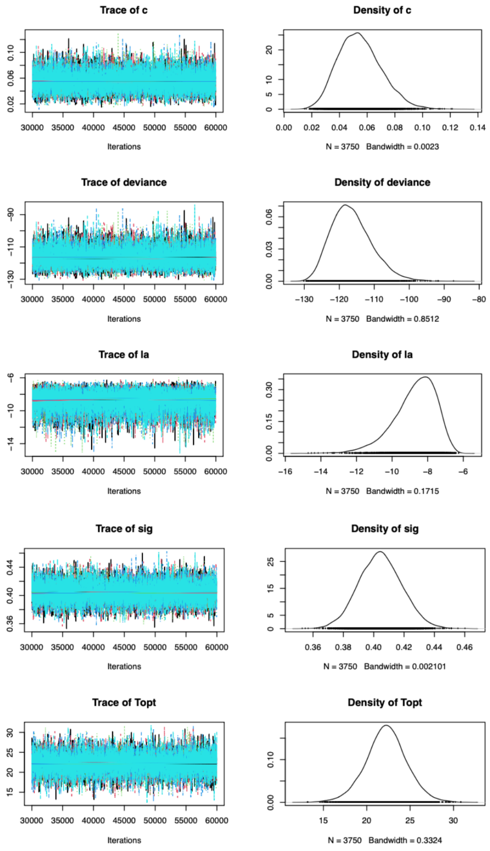


Figure S32: The trace plots for a randomly chosen dataset when modeling using individual inverse data, generated from an exponential distribution with a wide temperature interval (S32a) and a narrow temperature interval (S32b).

##### 5.1.4 Inverse of Mean Data

(a) Wide temperature interval (b) Narrow temperature interval

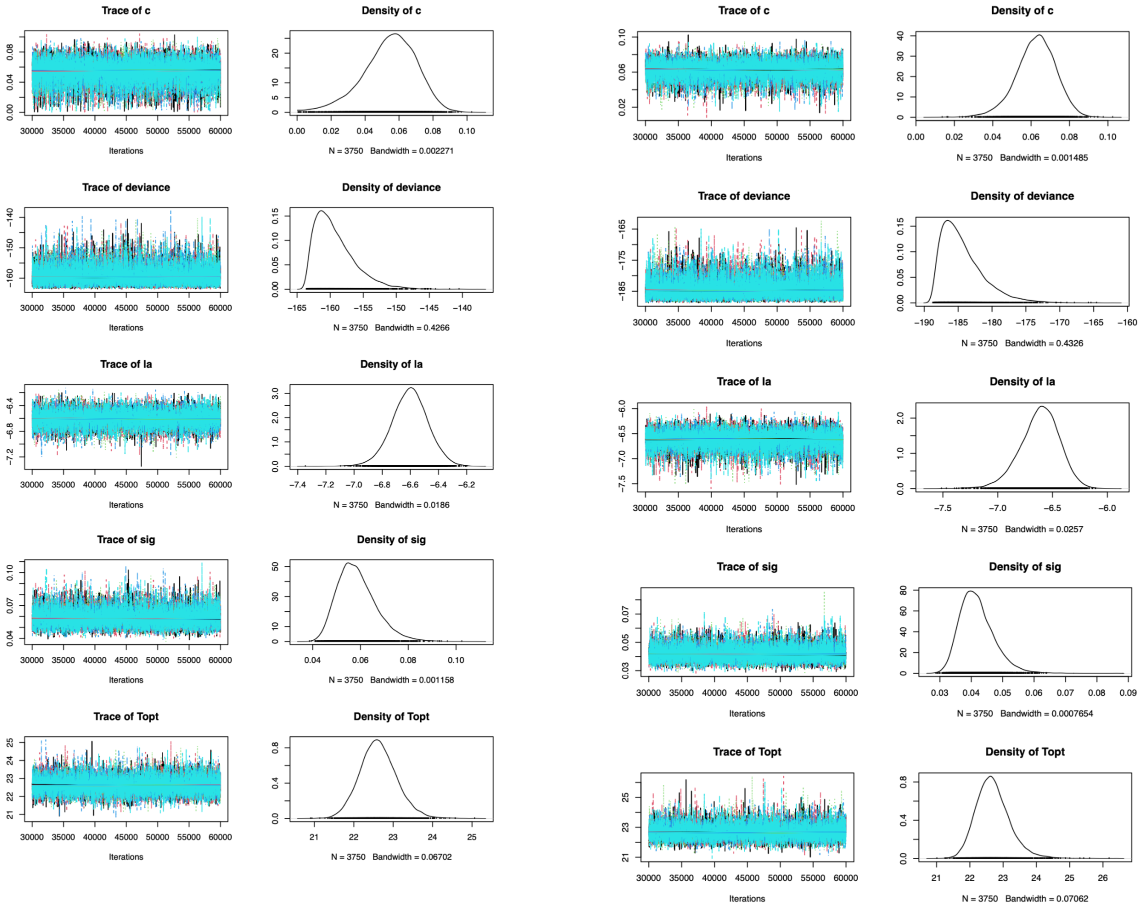


Figure S33: The trace plots for a randomly chosen dataset when modeling using the inverse of the mean data, generated from an exponential distribution with a wide temperature interval (S33a) and a narrow temperature interval (S33b).

##### 5.1.5 Mean of Inverse Data

(a) Wide temperature interval (b) Narrow temperature interval

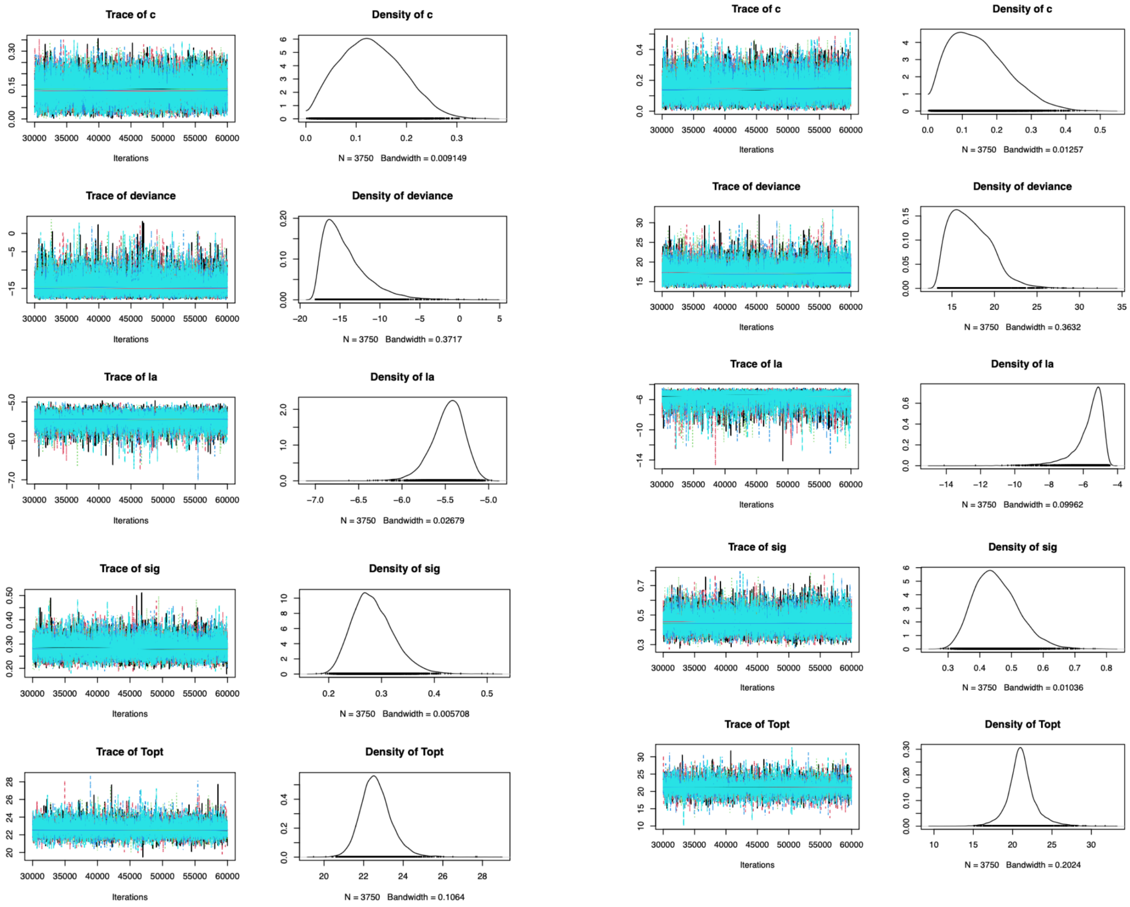


Figure S34: The trace plots for a randomly chosen dataset when modeling using the mean of the inverse data, generated from an exponential distribution with a wide temperature interval (S34a) and a narrow temperature interval (S34b).

#### 5.2 Weibull Data

##### 5.2.1 Individual Data, Weibull Model

(a) Wide temperature interval

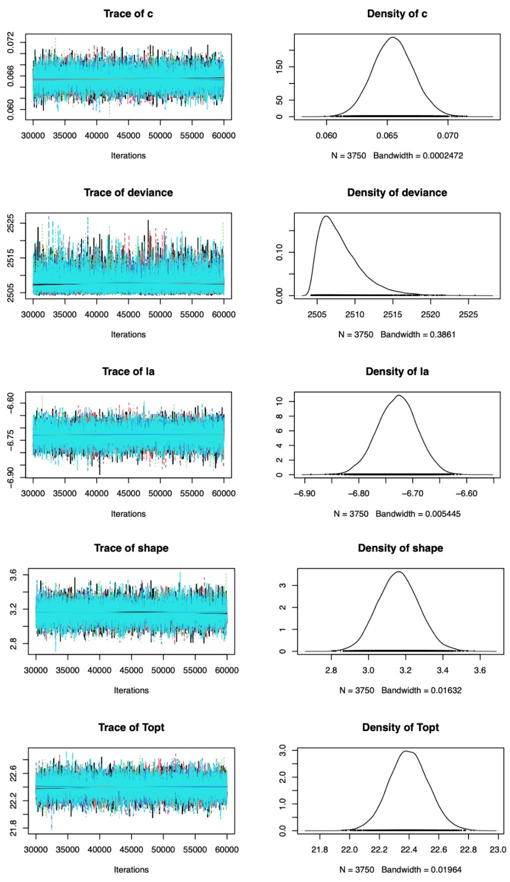


(b) Narrow temperature interval

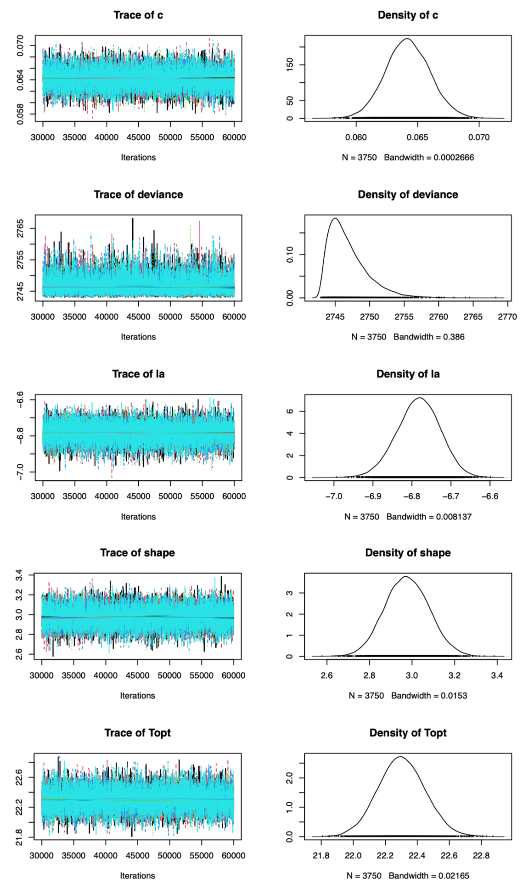


Figure S35: The trace plots for a randomly chosen dataset when modeling using individual-level data and a Weibull model, generated from a Weibull distribution with a wide temperature interval (S35a) and a narrow temperature interval (S35b).

##### 5.2.2 Individual Data, Exponential Model

(a) Wide temperature interval (b) Narrow temperature interval

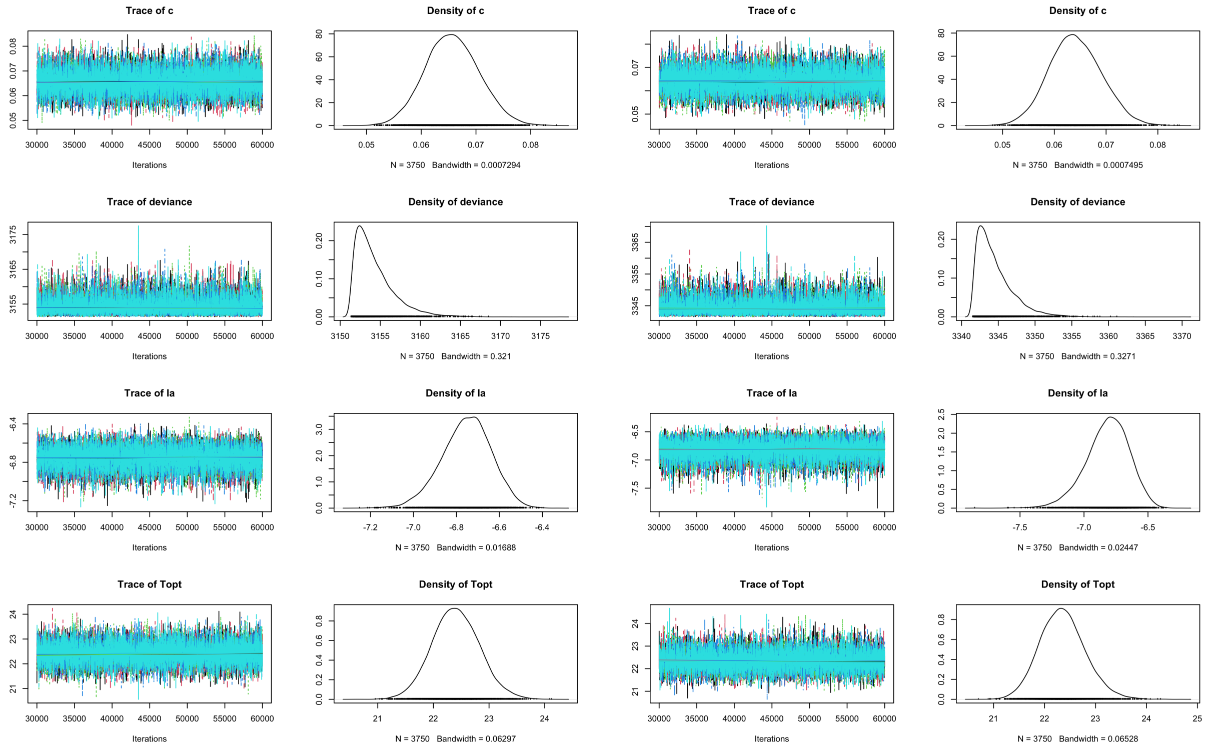


Figure S36: The trace plots for a randomly chosen dataset when modeling using individual-level data and an exponential model, generated from a Weibull distribution with a wide temperature interval (S36a) and a narrow temperature interval (S36b).

##### 5.2.3 Mean Data

(a) Wide temperature interval

(b) Narrow temperature interval

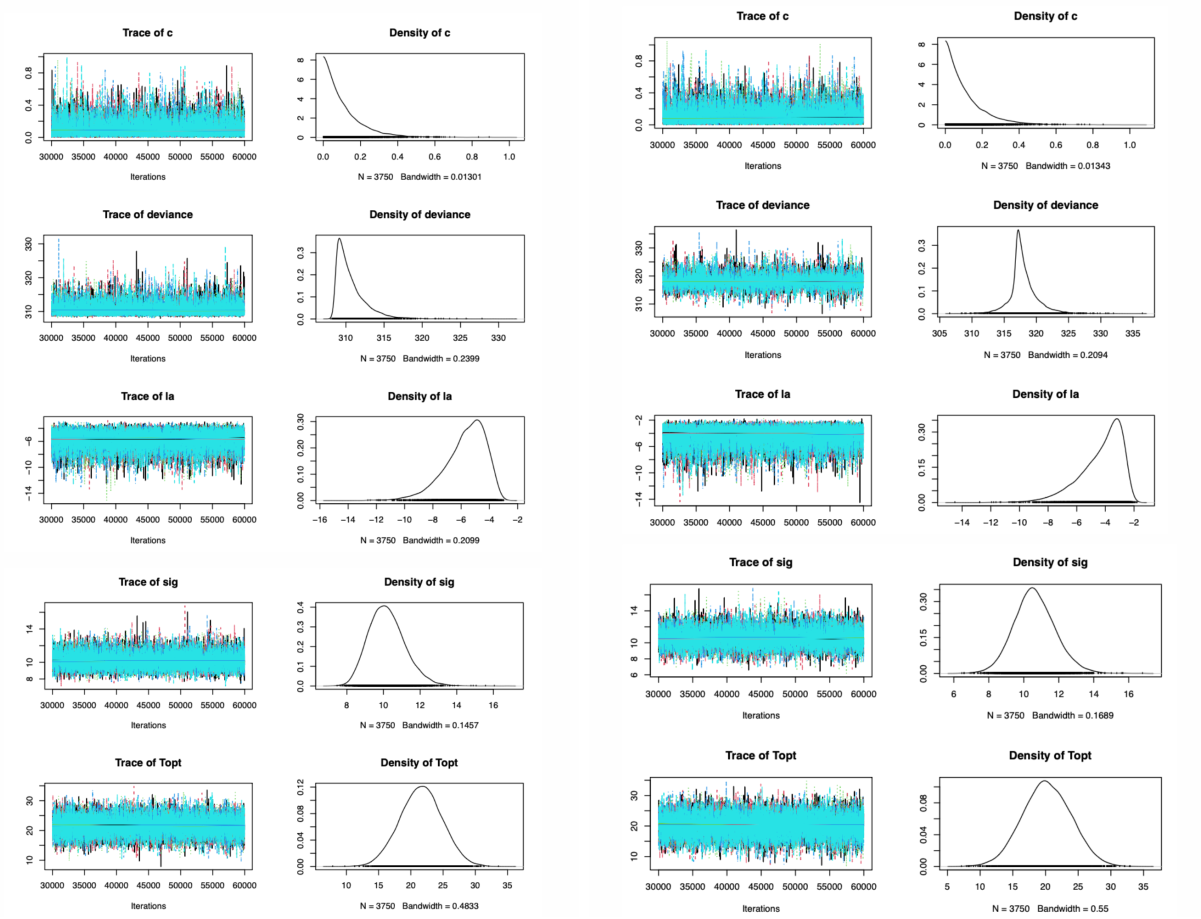


Figure S37: The trace plots for a randomly chosen dataset when modeling using mean data, generated from a Weibull distribution with a wide temperature interval (S37a) and a narrow temperature interval (S37b).

##### 5.2.4 Inverse Data

(a) Wide temperature interval (b) Narrow temperature interval

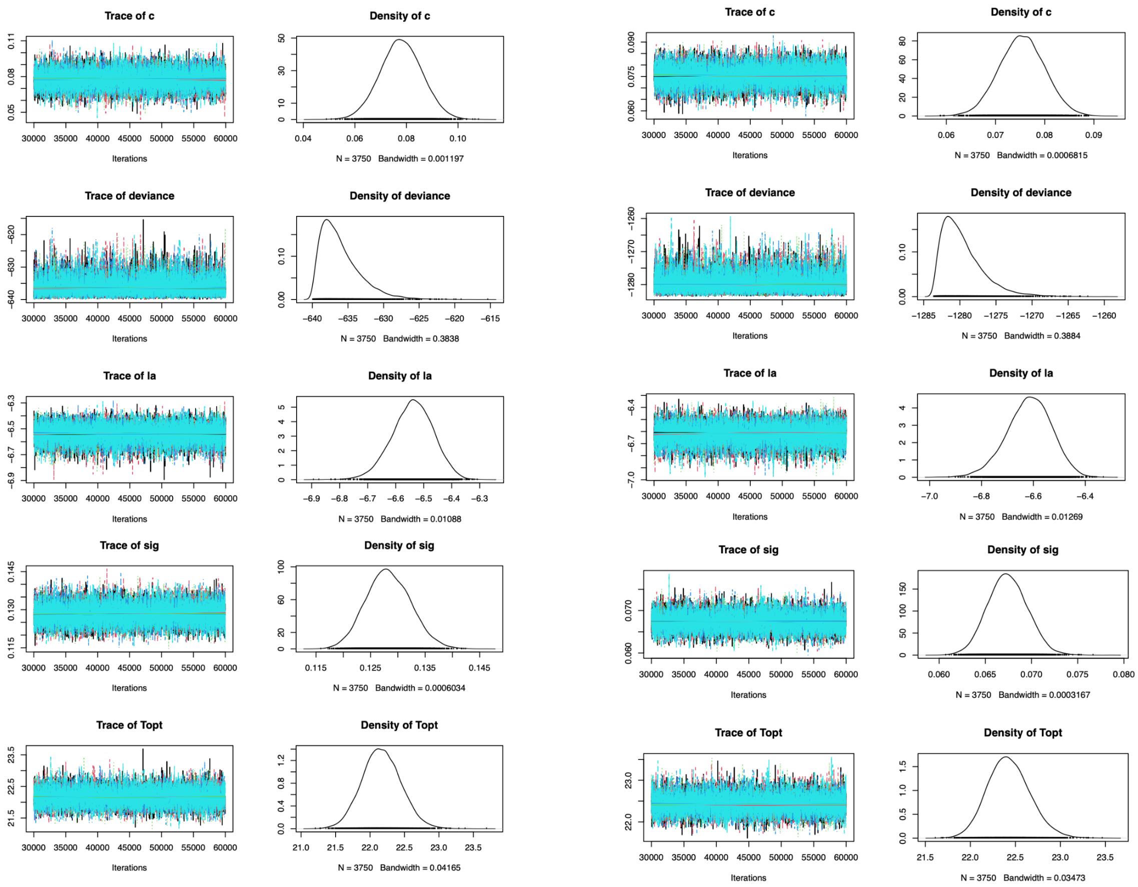


Figure S38: The trace plots for a randomly chosen dataset when modeling using individual inverse data, generated from a Weibull distribution with a wide temperature interval (S38a) and a narrow temperature interval (S38b).

##### 5.2.5 Inverse of Mean Data

(a) Wide temperature interval (b) Narrow temperature interval

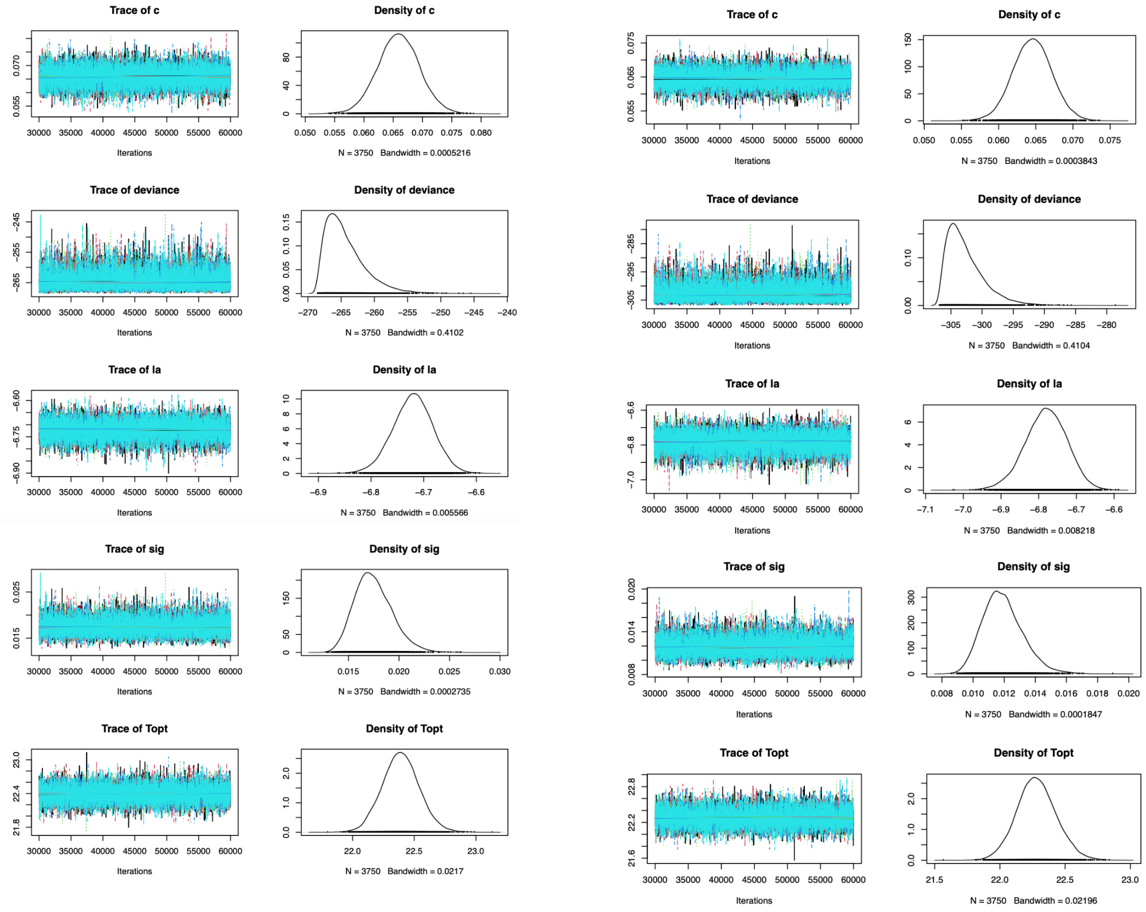


Figure S39: The trace plots for a randomly chosen dataset when modeling using the inverse of the mean data, generated from a Weibull distribution with a wide temperature interval (S39a) and a narrow temperature interval (S39b).

##### 5.2.6 Mean of Inverse Data

(a) Wide temperature interval

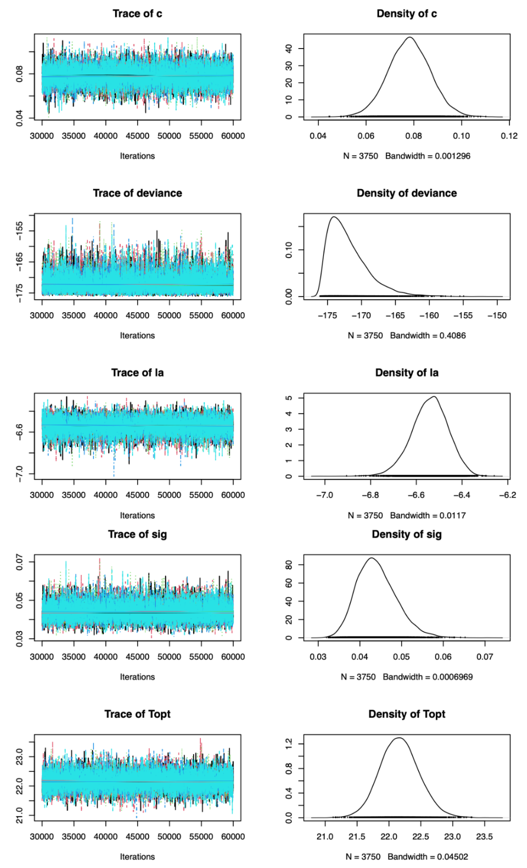


(b) Narrow temperature interval

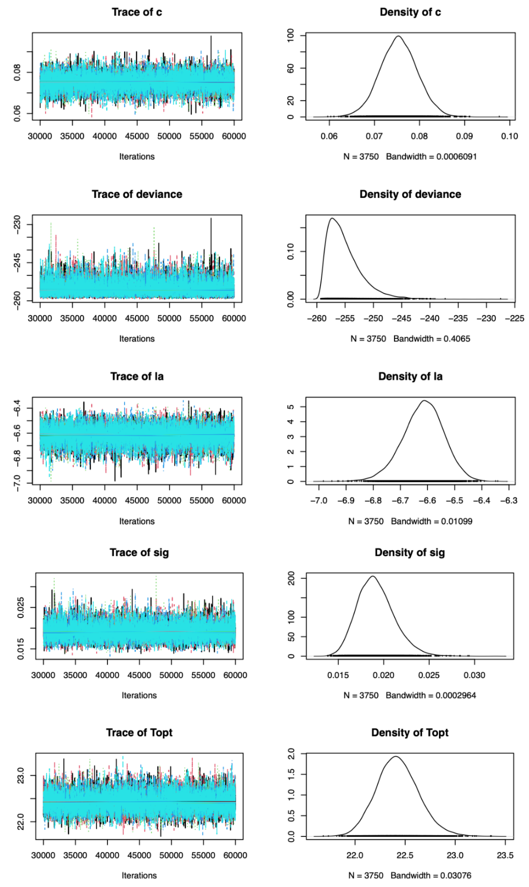


Figure S40: The trace plots for a randomly chosen dataset when modeling using the mean of the inverse data, generated from a Weibull distribution with a wide temperature interval (S40a) and a narrow temperature interval (S40b).
